# Multi-conflict islands are a widespread trend within *Serratia* spp

**DOI:** 10.1101/2024.01.28.577318

**Authors:** Thomas Cummins, Stephen R. Garrett, Tim R. Blower, Giuseppina Mariano

## Abstract

Bacteria carry numerous anti-phage systems in ‘defence islands’ or hotspots. Recent studies have delineated the content and boundaries of these islands in various species, revealing instances of islands that encode additional factors, including antibiotic resistance, stress genes, Type VI Secretion System (T6SS)-dependent effectors, and virulence factors.

Our study identifies three defence islands in the *Serratia* genus with a mixed cargo of anti-phage systems, virulence factors and different types of anti-bacterial modules, revealing a widespread trend of co-accumulation that extends beyond T6SS-dependent effectors to colicins and contact-dependent inhibition systems. We report the identification of four distinct anti-phage system/subtypes, including a previously unreported Toll/IL-1 receptor (TIR)-domain-containing system with population-wide immunity, and two loci co-opting a predicted T6SS-related protein for phage defence. This study enhances our understanding of the protein domains that can be co-opted for phage defence and of the diverse combinations in which known anti-phage proteins can be assembled, resulting in a highly diversified anti-phage arsenal.

## INTRODUCTION

Bacteria utilize hundreds of diverse anti-phage systems to overcome the evolutionary pressure posed by their viral predator, bacteriophages (phages). Many of these novel systems were only recently discovered and have shown remarkable diversity in their mechanisms of phage inhibition^1–5^. Interestingly, a considerable number of newly-discovered anti-phage systems show homology and resemblance to components of the innate immunity pathways of eukaryotes ^6–11^. For example, several anti-phage systems harbour a Toll/IL-1 receptor (TIR) domain ^4,5,12,13^. In the Thoeris system this domain is used for the production of the isomer of cyclic adenosine 5’-diphosphate-ribose (cADPR) signalling molecule whereas in other instances TIR domains can harbour NADase activity that leads to impaired survival of phage-infected cells^1,4,12,14^. The Bil system represents another clear example wherein innate immunity pathways first evolved in bacteria. This operon was shown to conjugate ubiquitin to released viral particles following phage infection, rendering them less infectious and unable to further propagate within the bacterial population^7,8^.

The tendency of anti-phage systems to cluster together in genomic ‘defence islands’ or hotspots has been extensively exploited in efforts to discover new players in bacterial immunity ^2,4,15–17^. Recently more and more studies have demonstrated that anti-phage systems and defence islands can be carried on mobile genetic elements (MGE) such as P2- and P4-like prophages and phage-inducible chromosomal islands (PICIs)^3,17–19^. The type of defence arsenal carried is highly variable even within the same genus ^3,18^. An interesting emerging trend suggests that in some genera, including *Salmonella* and *Vibrio*, defence hotspots frequently carry a mixed cargo^20,21^. In *Salmonella,* genomic hotspots carry a variable cargo of anti-phage systems, antibiotic resistance genes and virulence factor ^20^. Similarly, the gamma-mobile-trio (GMT) islands, initially discovered in *Vibrio parahaemolyticus*, represents the first reported example of co-localisation of anti-phage systems and Type VI Secretion System (T6SS)-mediated anti-bacterial effectors^21^. Nevertheless, the ratio of anti-phage and anti-bacterial modules of GMT islands exhibit a genus-specific variation^21^. The marked diversity observed in genomic hotspots, flanking genes, and substantial variation in cargo genes across different bacterial species highlights the importance for targeted investigations of these islands within specific genera. Such studies can reveal intricate details about the mobilisation and clustering of defence systems, anti-bacterial toxins, and virulence factors. Moreover, these focused studies may uncover novel types or subtypes of anti-phage systems.

The *Serratia* spp. pangenome demonstrates a notable degree of variability and plasticity^22^. Among the publicly available *Serratia* spp. genomes, *Serratia marcescens* is over-represented within the genus, reflecting its increasing relevance in clinical settings as an opportunistic pathogen^23,24^. *S. marcescens* strains are also extensively used to investigate several biological processes, including Type VI Secretion System (T6SS)-mediated anti-bacterial competition^25^.

In this work, we provide insights in the anti-phage arsenal and the type of defence islands found in the genus *Serratia.* By analysing the British Society for Antimicrobial chemotherapy (BSAC) collection of *S. marcescens* isolates^26^, we identify three defence hotspots conserved within the *Serratia* genus. Our findings demonstrate that the three identified islands harbour a mix of anti-phage systems and distinct types of virulence factors and anti-bacterial proteins. The type of virulence or anti-bacterial effector encoded is specific for each genomic island.

Through further investigation of the widespread LptG-YjiA island, which exhibits the highest variability in anti-phage system cargo, we identified previously unreported anti-phage systems/ subtypes, renamed **S**erratia **D**efence **I**sland **C**andidates (SDIC1-4). SDIC4 represent the first examples of an anti-phage system showing structural similarity to a predicted T6SS-related protein, VasI. The anti-phage system SDIC1 harbours a TIR-like domain and a second component with structural similarity to a putative ubiquitin ligase. We show that SDIC1 elicits protection against phage infection through population-wide immunity through SDIC1A, while SDIC1B alleviates the toxicity exerted by SDIC1A.

In summary, our data reveal a prevalent trend of defence hotspots acting as ‘multi-conflict’ islands in *Serratia* spp., accumulating a diverse array of conflict systems, including anti-phage systems, anti-bacterial proteins, and virulence effectors. This phenomenon extends beyond T6SS-dependent effectors to other anti-bacterial systems, such as colicins and contact-dependent inhibition systems (Cdi). Our study further expands the repertoire of known anti-phage systems and subtypes, underscoring the widespread tendency of known defence proteins to be re-shuffled in various combinations, defining numerous and distinct anti-phage systems and subtypes.

## RESULTS

### Identification of defence islands in the *Serratia*

To identify defence islands in *S. marcescens*, we started from the genomes of a group of strains collected by the British Society for Antimicrobial chemotherapy (BSAC). These represent 205 clinical isolates that were isolated from the bloodstream of patients^26^. *S. marcescens* BSAC strains are closely related, nevertheless, they encode a relatively high diversity of known anti-phage systems (57 systems) (Figure S1a-b, Table S1). Within the group of known anti-phage systems identified, PD-T4-6, Thoeris, and R-M type I are found more frequently (Figure S1b and Table S1). Furthermore, using the latest version of PADLOC^27^, 13 additional PADLOC-specific putative systems, that currently remain unverified experimentally, were detected (Figure S1a-b, Table S1).

Next, 50kb upstream and downstream of experimentally confirmed anti-phage systems were retrieved to allow the inclusions of islands that contain multi-gene systems such as BREX and DISARM. Proteins found in each genomic neighbourhood were clustered together using MMseqs2^28^ and inspected to find putative boundaries of defence islands (Table S2). Putative defence-islands boundaries were used as queries to search a Refseq database with cblaster^29^. This program leverages BLASTp to identify genomes wherein hits are co-localised^29^. Genomic islands were identified as defence-enriched hotspots only if, when searching a Refseq database, they contained known anti-phage systems in at least 30% of *Serratia* spp. genomes.

The first putative defence hotspot identified with this method is delimited by *speG*, encoding a spermidine N1-acetyltransferase, and *pstB,* encoding a phosphate ABC transporter. The genomic region defined by SpeG-PstB is found in most *Serratia* genomes, albeit in some cases this is not occupied by any additional genes (Figure 1a). Interestingly, most instances of occupied SpeG-PstB hotspots carry a Thoeris system or another, unidentified, TIR-containing defence system (Figure 1a and Table S3-S4). Occupied SpeG-PstB hotspots frequently carry a haemolysin-like toxin and its cognate, secreting partner (Figure 1a and Table S3-S4).

**Figure 1.**
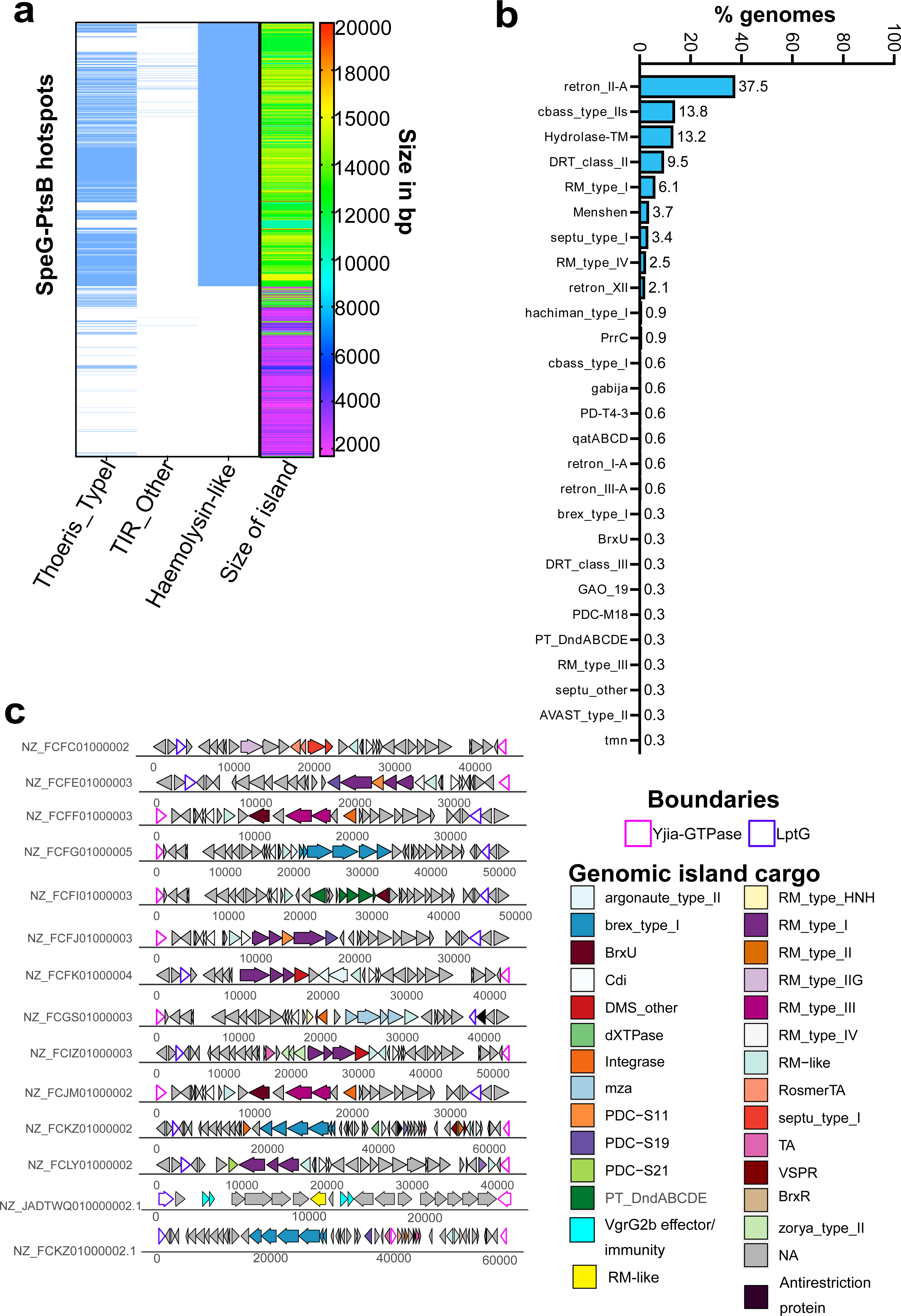
Defence islands identified in the *Serratia* genus. **(a)**Heatmaps showing presence-absence (blue-white) of the Thoeris I system, a TIR domain-containing unknown system and a haemolysin-like toxin system in SpeG-PstB hotspots in relation to the size of the genomic island (shown as a rainbow heatmap on the right). Legend on the right shows the rainbow colour mapping in relation to the size of the island in base-pairs (bp). **(b)** Bar plot summarising the percentage of occurrence of anti-phage systems identified in occupied LysR-esterase hotspots. **(c)** Schematic representation of the genomic organisation of LptG-YjiA island of representative strains of the BSAC collection. LptG and YjiA boundaries are represented with coloured outlines. Anti-phage systems predicted by PADLOC are coloured as indicated in the legend. Integrases, transposases, the WYL-containing regulator BrxR, toxin-antitoxin systems (TA), a predicted T6SS-dependent effector (VgrG2b) and contact-dependent inhibition (Cdi) systems are also coloured according to the legend.

A second hotspot, defined by a LysR transcriptional regulator and an esterase (Table S5), harbours a known anti-phage system in ∼30% of instances (Table S5-S6). The island encodes a diverse array of 27 different anti-phage system, with retron IIA, CBASS IIa, Hydrolase-TM and DRT II being the most abundant (Table S6-Figure 1b).

Finally, we identified a third defence-hotspot encoded between LptG and a GTPase. We note that the protein annotated as GTPase is homologous to YjiA, a protein that, together with YjiO, defines the boundaries of a defence island in *Escherichia coli*^17^. Unlike *E. coli*, we did not find YjiO homologues in *S. marcescens* strains (Figure 1c and Table S2). In BSAC strains, the LptG-YjiA hotspot carries a variable defence cargo that accounts for 30% (17/57) of the known anti-phage systems identified in the BSAC collection (Figure S1a-b, Figure 1b and Table S7), with type I RM, type IV RM and type II argonaute being the most abundant systems (Figure S1a-b, Table S7).

Within the LptG-YjiA islands of the BSAC collection we frequently found Type II toxin-antitoxin systems, Cdi systems, and a protein annotated as ‘Rhs IV effector’ (Table S2). We used PFAM and HHPRED^30,31^ to characterise the latter and found it harbours structural homology with a predicted ‘VgrG2b C-terminal effector domain’ (Figure 1b). This domain was first reported in *P. aeruginosa* as a specialised T6SS effector with metallopeptidase activity, normally found fused with the structural protein VgrG^32^. In the LptG-YjiA islands, the VgrG2b toxic domain is found as cargo effector (not fused to T6SS structural proteins) and encoded next to its predicted immunity lipoprotein (Figure 1b, Table S2)^32^.

By using cblaster with relaxed search parameters that allow to report regions that are split over two contigs, we found that LptG-YjiA islands are present in the majority of *Serratia* spp. (Table S8). Nevertheless, further analysis was only performed on LptG-YjiA islands that span one contig (Table S9). Anti-phage systems predictions highlighted that LptG-YjiA islands contain at least 1 known anti-phage system in 90% instances and that they carry ∼60 different anti-phage systems (Table S10). Consistent with what observed with the BSAC collection, type I RM, type IV RM and type II argonautes remain enriched in *Serratia* LptG-YjiA hotspots (Figure 2a and Table S10). Furthermore, extending the analysis to all *Serratia* spp. revealed that type II RM, mza, type I BREX and Zorya II are also overrepresented (Figure 2a-b). We further report that the islands defined by LptG and YjiA are always occupied by several genes and no instances of empty (unoccupied) islands could be detected (Figure 2c).

**Figure 2.**
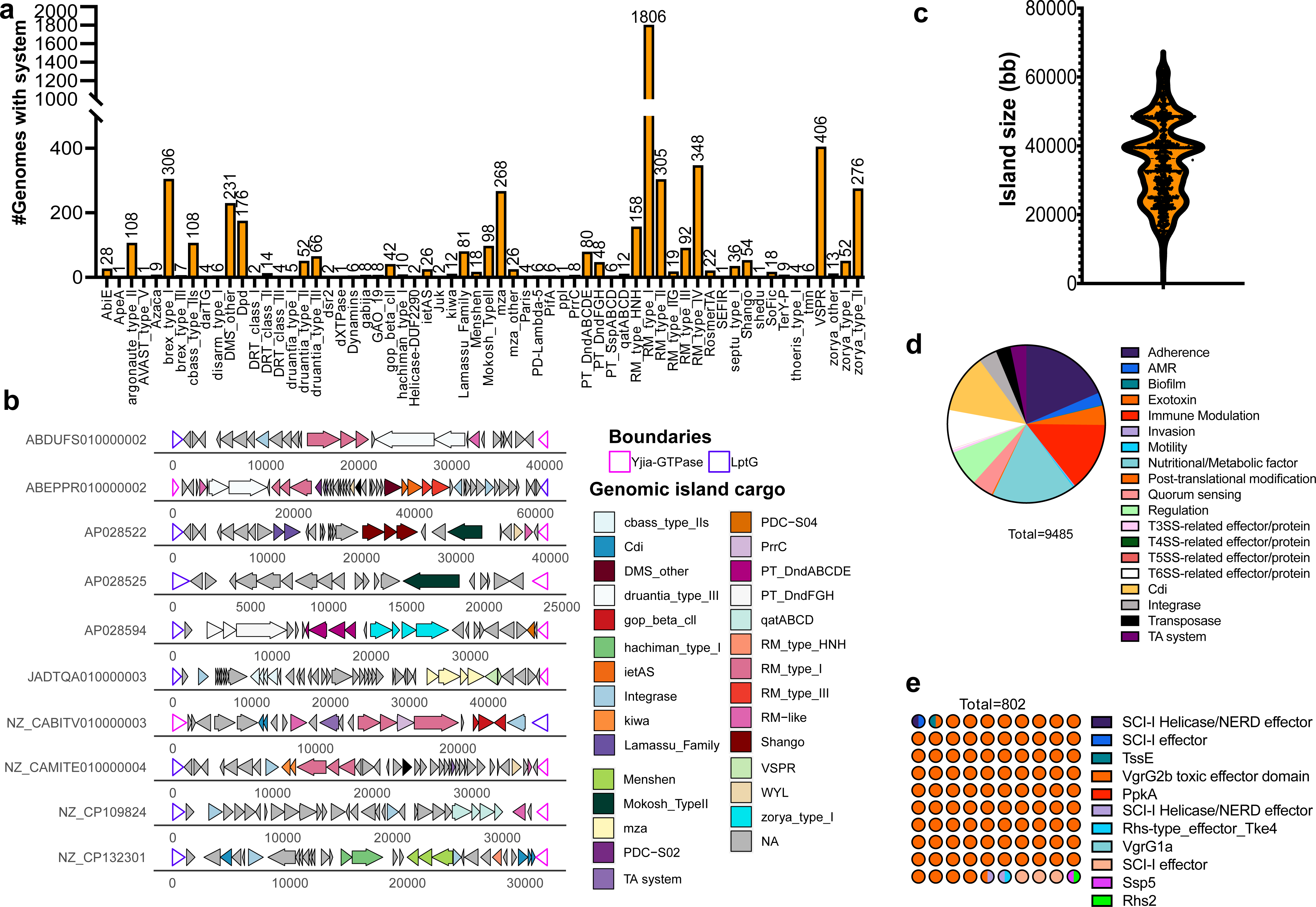
LptG-YjiA islands across the *Serratia* genus carry a high diversity of defence systems. **(a)** Prevalence of anti-phage systems across *Serratia* species with a complete LptG-YjiA island localised across one contig. **(b)** Length of LptG-YjiA islands found across *Serratia* species. **(c)** Schematic representation of the genomic organisation of representative LptG-YjiA islands chosen across the *Serratia* genus. Boundaries are represented with coloured outline and anti-phage systems, anti-bacterial effectors, TAs, BrxR, integrases and transposases are coloured as indicated in the legend. **(d)** Distribution of predicted virulence factors, predicted T3SS-dependent effectors, predicted T4SS-dependent effectors, predicted T5SS two-partner systems, predicted T6SS-dependent effectors and core components, contact-dependent inhibition systems (Cdi), TA systems, transposases, and integrases in *Serratia* LptG-YjiA islands. The VFDB database classification was used to categorise virulence factors as i) adherence, ii) biofilm, iii) motility, iv) immune-modulation, v) nutritional/metabolic factor, vi) exotoxins, vii) invasion, viii) quorum sensing and, ix) post-translational regulation. **(e)** Waffle plot showing the prevalence of T6SS-predicted effectors found in *Serratia* LptG-YjiA islands.

### *Serratia* spp. islands are prevalently ‘multi-conflict’ islands

Given the increased variability in the arsenal of defence systems carried by LptG-YjiA islands and the consistent observation of these islands being occupied, we set to further explore their cargo.

To better characterise the protein content of each *Serratia* spp. LptG-YjiA island we used MMseqs2 to cluster proteins encoded in each island and subsequently curated our dataset to remove clusters that were predicted as known defence systems by PADLOC (Table S11). The remaining sequences were analysed using AMRfinder, Secret6, and PSI-BLAST comparisons against the Virulence Factor Database (VFDB) (Table S12-S13)^33–35^. With these methods we were able to classify 9485 proteins (Table S12-S13). LptG-YjiA islands encode few antibiotic resistance genes, mostly related to fosfomycin and gentamycin resistance (Table S12-S13). When using VFDB, we classified proteins according to the virulence factors categories defined by the database (http://www.mgc.ac.cn/VFs/VFcategory.htm)^34^. Putative virulence factors involved in adherence (fimbriae, pili, adhesins), immune modulation, regulation, and metabolic factors (metal and ions uptake) are found to be overrepresented in LptG-YjiA islands (Figure 2d). Additionally, within LptG-YjiA islands, quorum sensing factors, predicted T6SS effectors/components and Cdi systems are also enriched (Figure 2d, Table S11-13). A closer look to the group of predicted T6SS-dependent effectors revealed that these are mostly represented by homologues of the orphan VgrG2b toxic domain (Figure 2e). We thus wondered whether this could represent an example of toxins adapted for phage defence^36^. We tested a homologue of the VgrG2b toxic domain and its putative immunity protein (WP_108680118.1 and WP_108680119.1) in *E. coli* MG1655 and *S. marcescens* Db10. The effector-immunity pair were expressed under the control of the constitutive Tat promoter in a low-copy number plasmid, and, in both systems, they could not provide immunity against phages (Figure S2a-b), confirming this operon could instead represent an orphan T6SS effector-immunity pair. Further investigation will shed light on the events that have driven the accumulation of this particular pair in *Serratia* LptG-YjiA islands.

HHPRED was additionally used with an e-value threshold (< 0.05) (Table S14) to predict the role of proteins that were not defined with the above methods. This analysis showed that LptG-YjiA islands contain numerous predicted AAA^+^ ATPases, DEAD box helicases, STAND and NACHT domains, ubiquitin-like proteins, Zn^+^ metallo-peptidases, restriction enzymes and several TIR domains that were not recognised as part of known anti-phage systems by PADLOC. These could be part of novel systems or of distinct subtypes of existing anti-phage systems (Table S14). Alternatively, these could represent examples of known defence systems with highly divergent protein sequences, causing the lack of prediction by PADLOC. Inspection with HHPRED confirmed the presence of integrases, transposases and recombinases, additional toxin-antitoxin systems as well as several more predicted T6SS-related proteins (Table S14). In particular, we detected the presence of proteins with a VasI-like fold (Table S14). VasI is a predicted T6SS-associated protein, albeit its function remains poorly understood^37^. Whilst HHPRED predictions indicate that VasI-like proteins found in *Serratia* LptG-YjiA hotspots are structural homologues of the T6SS-dependent VasI, they do not share sequence identity by BLAST. VasI-like proteins carried within LptG-YjiA islands are often encoded close to the WYL-containing protein BrxR, transcriptional regulators strongly linked to anti-phage defence (Figure S2c)^38,39^, thus these might represent an example wherein a protein domain, normally involved in a different function, has been co-opted for phage defence. We additionally found various predicted multidrug-efflux pumps, stress response genes, additional virulence factors (such as VapE, IrmA and hyaluronate lyase) and proteins with predicted roles in transport, metabolism, and modification of sugars (Table S14). The latter could play a role in surface modification and masking of phage receptors. Notably, even following the employment of HHPRED, the function of many proteins in LptG-YjiA islands could not be predicted (Table S14). The investigation of these proteins may reveal new players in i) anti-phage defence, ii) virulence or iii) anti-bacterial competition.

Given the tendency of LptG-YjiA hotspots to accumulate a variable cargo and the over-representation of a haemolysin-like toxin in occupied SpeG-PstB hotspots, we queried whether this is a common pattern within *Serratia* islands. Using a PFAM database, we found that LysR-esterase islands also carries virulence factors (RhuM, Table S15) and anti-bacterial effectors (Table S15). Interestingly, whilst LptG-YjiA islands prevalently carry Cdi and predicted T6SS-dependent proteins, colicins and pyocins are prevalent in LysR-esterase hotspots (Table S15 and Figure S3).

Taken together, our analysis of *S. marcescens* strains from the BSAC collection and the *Serratia* genus uncovered three ‘multi-conflict’ islands wherein anti-phage systems, toxin-antitoxin systems, and proteins involved in anti-bacterial competition and virulence are accumulated. Whilst the anti-phage system composition is mostly variable, the type of virulence or anti-bacterial protein encoded is specific to each island.

### LptG-YjiA islands are widespread multi-conflict islands in Enterobacteria

YjiA was previously identified as the boundary of a defence hotspot in several *E. coli* genomes, with YjiO, a multi-drug efflux transporter, identified as the second boundary of this defence island^17^. Despite the relatedness of *E. coli* and *S. marcescens* and the use of relaxed parameters in cblaster, no homologues of YjiO were detected in *Serratia* LptG-YjiA islands (Table S7, Table S8, Table S16). While other multidrug-efflux pumps are encoded within *Serratia* LptG-YjiA islands, they are not universally present in every island, ruling them out as possible conserved boundaries (Table S14).

Consequently, we sought to understand the overlap between the two genomic islands in *E. coli, Serratia* spp., and other Enterobacteria. Cblaster was run using LptG, YjiA, and YjiO as query sequences (Table S16). These searches confirmed that in *Serratia* islands, YjiO homologues are not present (Figure 3). Islands containing all three LptG-YjiO-YjiA are instead widespread in several Enterobacteria, particularly enriched in *E. coli* and *Salmonella* spp. (Table S16). The prevalence of *E. coli* and *Salmonella* spp. might be influenced by the over-representation of genomes from these organisms on NCBI. Therefore, for further analysis, genomes were filtered using fastANI to eliminate sequence duplications and exclude closely related genomes (90% sequence identity).

**Figure 3.**
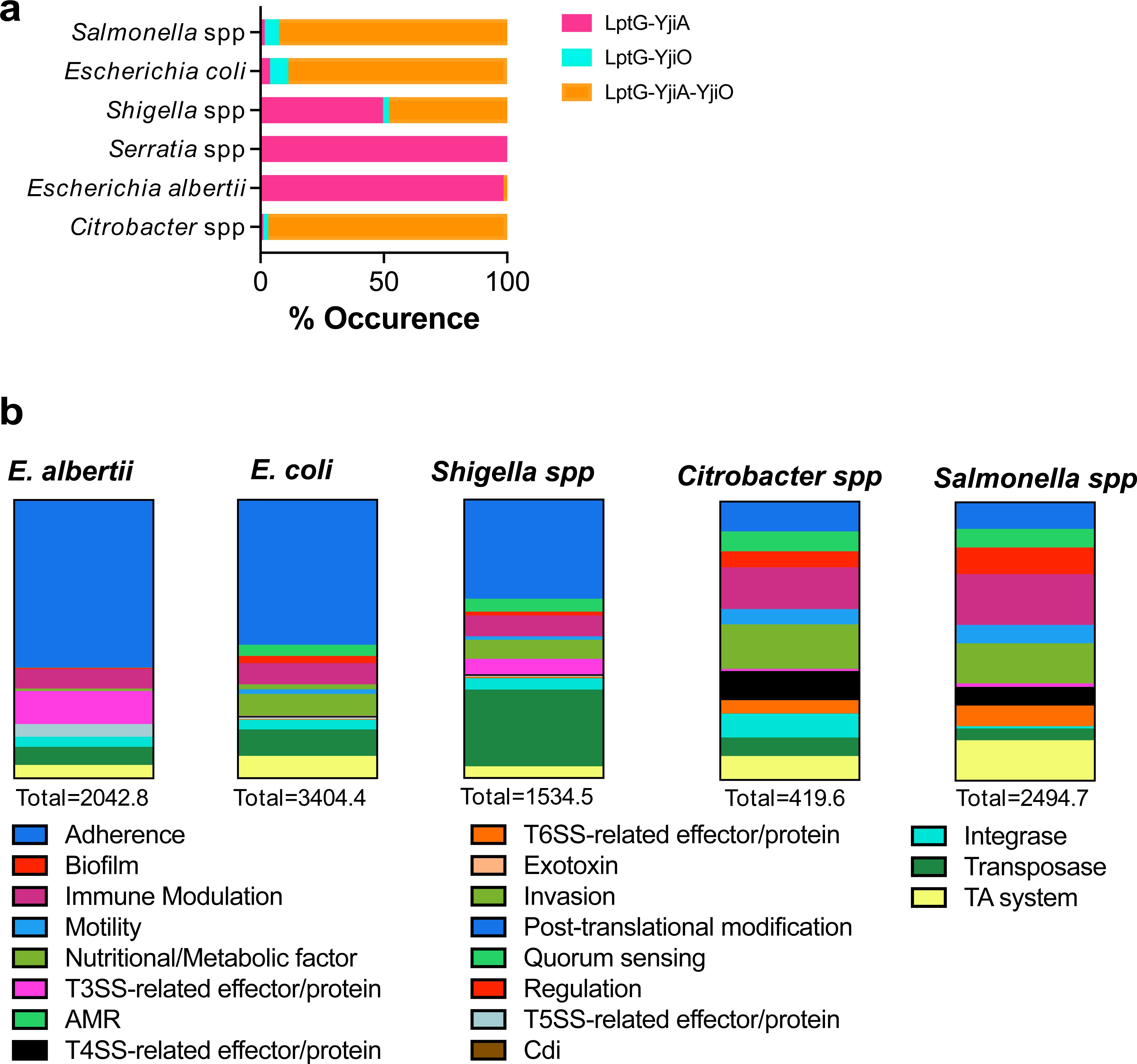
LptG-YjiA and LptG-YjiO-YjiA islands are widespread in Enterobacteria. **(a)** Bar plot showing the prevalence of islands in Enterobacteria with either LptG-YjiA, LptG-YjiO or LptG-YjiO-YjiA as boundaries and insertion points for acquisition of defence systems. **(b)** Distribution of predicted virulence factors, predicted T3SS-dependent effectors, predicted T4SS-dependent effectors, predicted T5SS two-partner systems, predicted T6SS-dependent effectors and core components, contact-dependent inhibition systems (Cdi), TA systems, transposases, and integrases in Enterobacteria LptG-YjiA islands as shown in **panel a.** The VFDB database classification was used to categorise virulence factors as indicated in Figure 2d.

In *Escherichia albertii* strains, detected islands lack YjiO and have LptG-YjiA as boundaries (Figure 3a and Figure S4, Table S17). Conversely, in *E. coli* strains, the most common genomic organization is LptG-YjiO-YjiA, showing considerable variability within genera (Figure 3a and Figure S5). Notably, in *E. coli* strains, known anti-phage systems accumulate both between YjiO-YjiA and between LptG-YjiO, suggesting a high likelihood of recombination events in the intergenic spaces within LptG-YjiO-YjiA and the existence of two insertion points (between LptG-YjiO and YjiO-YjiA) (Figure S5 and Table S18). We note that *E. coli* strains found to carry a LptG-YjiA-YjiO island in our search are mostly distinct from those previously reported to harbour YjiO-YjiA islands, underscoring the high plasticity of defence islands within genera and species (Figure S5 and Table S18)^17^.

Islands carrying LptG-YjiO-YjiA are also prevalent in *Citrobacter* spp. and *Salmonella* spp., showing a similar pattern of anti-phage system accumulation between either LptG-YjiO or YjiO-YjiA (Figure 3, Figure S6, Figure S7, Table S16, Table S19, and Table S20). In the case of *Shigella* spp., approximately 50% of genomes carry an LptG-YjiO-YjiA island, while the remaining 50% have an island with LptG-YjiA as boundaries (Figure 3, Figure S8, and Table S21). Furthermore, rare instances were found where LptG-YjiO represent the boundaries of the defence islands, with YjiA being lost (Figure S4-S8, Table S16-S21). In other instances, the three boundaries are retained, but YjiA and YjiO positions are inverted (LptG-YjiA-YjiO) (Figure S4-S8, Table S16-S21). For simplicity, henceforth in this study, we will refer to these islands indiscriminately as LptG-YjiA islands.

The analysis of non-defence proteins revealed that, like *Serratia*, LptG-YjiA islands in other species can carry diverse genes involved in virulence or inter-bacterial competition (Figure 3b). Interestingly, our data indicate that the type of cargo can vary for each genus, with *Citrobacter* spp. and *Salmonella* spp. harbouring the highest number of predicted T6SS-related and T4SS-related proteins and *E. albertii* carrying the highest number of predicted T3SS-related proteins amongst the genomes analysed (Figure 3b). Interestingly, we did not find Cdi systems in LptG-YjiA islands of species other than *Serratia* (Figure 2 and 3b). While rare, we note that in some *Citrobacter* spp., a full T6SS cluster is found within LptG-YjiA islands (Figure 3b and Figure S7). Similar sporadic instances were also found by manual inspections of Serratia LptG-YjiA islands that were split over two contigs (Table S8) and sporadically in LysR-esterase hotspots (Figure S3b).

Taken together, our data show that LptG-YjiA islands are predominantly ‘multi-conflict’ islands that accumulate a variable cargo across genera.

### *Serratia* LptG-YjiA islands contain previously unidentified anti-phage systems

Manual screening of the group of defence domains and proteins with unknown functions found by HHPRED in LptG-YjiA islands (Table S14) revealed the presence of four candidate anti-phage systems/subtypes, which were renamed **S**erratia **D**efence **I**sland **C**andidates (SDIC1-4)(Table 1 and Figure 4).The domain composition of SDIC1-4, predicted with HHPRED, is summarised in Table 1 and Figure 4.

**Figure 4.**
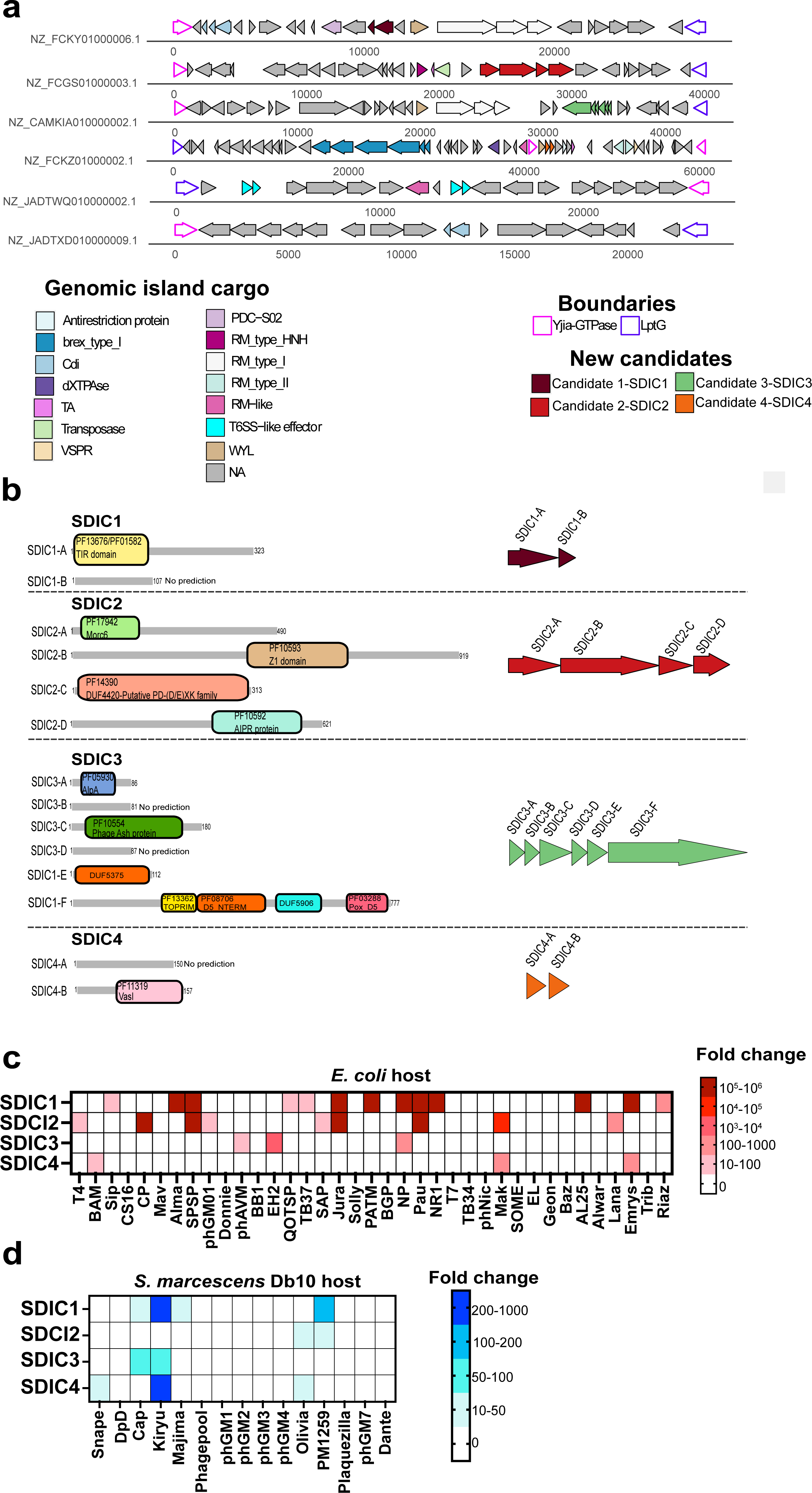
Novel anti-phage systems/subtypes are encoded in *Serratia* LptG-YjiA islands. **(a)**Schematic representation of the genomic neighbourhood of the four *Serratia* defence island candidates (SDIC1-4). **(b)**Schematic representation of the HHPRED domain predictions for the single components of SDCI1-4. **(c)** Evaluation of SDIC1-4 fold protection against a suite of coliphages when expressed in *E. coli* MG1655. Fold protection was calculated as the ratio of the efficiency of plating (EOP) calculated for a strain expressing one of SDIC1-4 candidates and the EOP value of a strain carrying the empty vector (VC, pQE60-Tat)(n= 3 biological replicates). **(d)** Evaluation of SDIC1-4 fold protection when expressed in *S. marcescens* Db10 and challenged against a group of *Serratia* phages (n= 3 biological replicates). Fold protection was calculated as in **(b).**

**Table 1.** Summary of the gene composition and domain prediction for the novel candidate anti-phage systems found in the Serratia defence islands.

| Candidate system | Genes | Representative protein ID | HHPRED |
| --- | --- | --- | --- |
| SDIC 1 | SDIC1A | WP_060442246.1 | TIR domain (PF13676.9) |
|  | SDIC1B | WP_060442245.1 | No prediction |
| SDIC2 | SDIC2A | WP_072270133.1 | Morc6_S5; Morc6 ribosomal protein S5 domain 2-like(PF17942.4) |
|  | SDIC2B | WP_072270134.1 | SWI2/SNF2 ATPase-like protein (7TN2,5JXR); AAA_34 -P-loop containing NTP hydrolase pore-1(PF13872.9)<br>Z1 domain(PF10593.12 ) |
|  | SDIC2C | WP_072270135.1 | DUF4420; Putative PD-(D/E)XK family member, (PF14390.9) |
|  | SDIC2D | WP_080437564.1 | AIPR protein(PF10592.12 ) |
| SDIC3 | SDIC3A | WP_261092931.1 | AlpA regulator |
|  | SDIC3B | WP_261145910.1 | No prediction |
|  | SDIC3C | WP_261145913.1 | host cell division inhibitor lcd-like protein<br>PF10554-Phage Ash protein |
|  | SDIC3D | WP_261145916.1 | No prediction |
|  | SDIC3E | WP_261145919.1 | DUF5375 family protein |
|  | SDIC3F | WP_261145922.1 | Poxvirus D5 protein (PF03288)<br>TOPRIM (PF13362.9)<br>D5 N-terminal like (PF08706.14)<br>DUF5906 (PF19263.2) |
| SDIC4 | SDIC4A | WP_077264947.1 | No prediction |
|  | SDIC4B | WP_060451809.1 | Vasl (PF11319.12) |

SDIC1 is a two-gene system that contains a TIR domain (SDIC1A) and a protein of unknown function (SDIC1B) that could not be defined by HHPRED (Table 1 and Figure 4a-b). SDIC2 has a similar but not identical domain composition to mza, and likely represents a novel mza subtype ^5,40^(Figure 4b). SDIC3 harbours some similarity to a prophage-encoded anti-phage system in ECOR61 identified during a transposon screen (Figure 4b)^5,40^. SDIC3C and SDIC3F have 27% and 17% sequence identity to their counterparts in ECOR61 respectively, whereas the other components are unique to SDIC3. Thus SDIC3 might represent a different subtype of the ECOR61 anti-phage system^40^. Finally, as a fourth candidate (SDIC4) we investigated the locus carrying the novel VasI-like protein (Table 1, Figure 4a-b, Table S14). The 4 candidates were cloned in a low copy plasmid, under the control of a constitutive promoter compatible with expression in *E. coli* and *S. marcescens.* In *E. coli,* SDIC1, SDIC2 and SDIC3 confer strong levels of protection against several phages (≥ 1000 fold), including those belonging to the Durham collection (Figure 4c)^41^, whereas SDIC4-mediated anti-phage activity is more modest (≥ 10-200-fold) but still significant (Figure 4c). To further confirm the role of SDIC1-4 in defence against phages, the same constructs were also tested in *S. marcescens* Db10 (which natively lacks the tested systems). All tested systems show the ability to reduce the efficiency of plating of several *Serratia* phages (Figure 4d), albeit with a lower amplitude than in *E. coli.* The different degrees of protection that SDIC1-4 confer in the two hosts could be related to their level of expression or the reduced size and diversity of the *Serratia* phage panel used.

In summary, our data reveal that inspection of unclassified defence proteins and proteins of unknown function in *Serratia* LptG-YjiA islands can reveal previously unreported anti-phage systems or subtypes.

### VasI-like proteins define two anti-phage system subtypes

To investigate if similar instances of VasI-like proteins involved in defence against phages can be found in other species, SDIC4B homologues were retrieved using PSI-BLAST and their genomic neighbourhood was analysed by flanking gene analysis (FlaGs)^42^.

This analysis highlighted that closely related homologues of SDIC4B can be found in numerous species and confirmed they are not routinely found in T6SS clusters. Instead, they are often associated to a BrxR homologue (Figure S9, Table S23), an indication that they do not represent a divergent T6SS-dependent protein but rather an example of a shared fold co-opted for phage defence (Figure S9, Table S23). The use of cblaster highlighted that SDIC4 homologues are encoded in ∼18,000 genomes in Genbank and mostly predominant in Enterobacteria, and particularly in *Klebsiella* spp., *Salmonella* spp., *Citrobacter* spp., *Enterobacter* spp. and *E. coli* strains (Figure 5a and Table S24).

**Figure 5.**
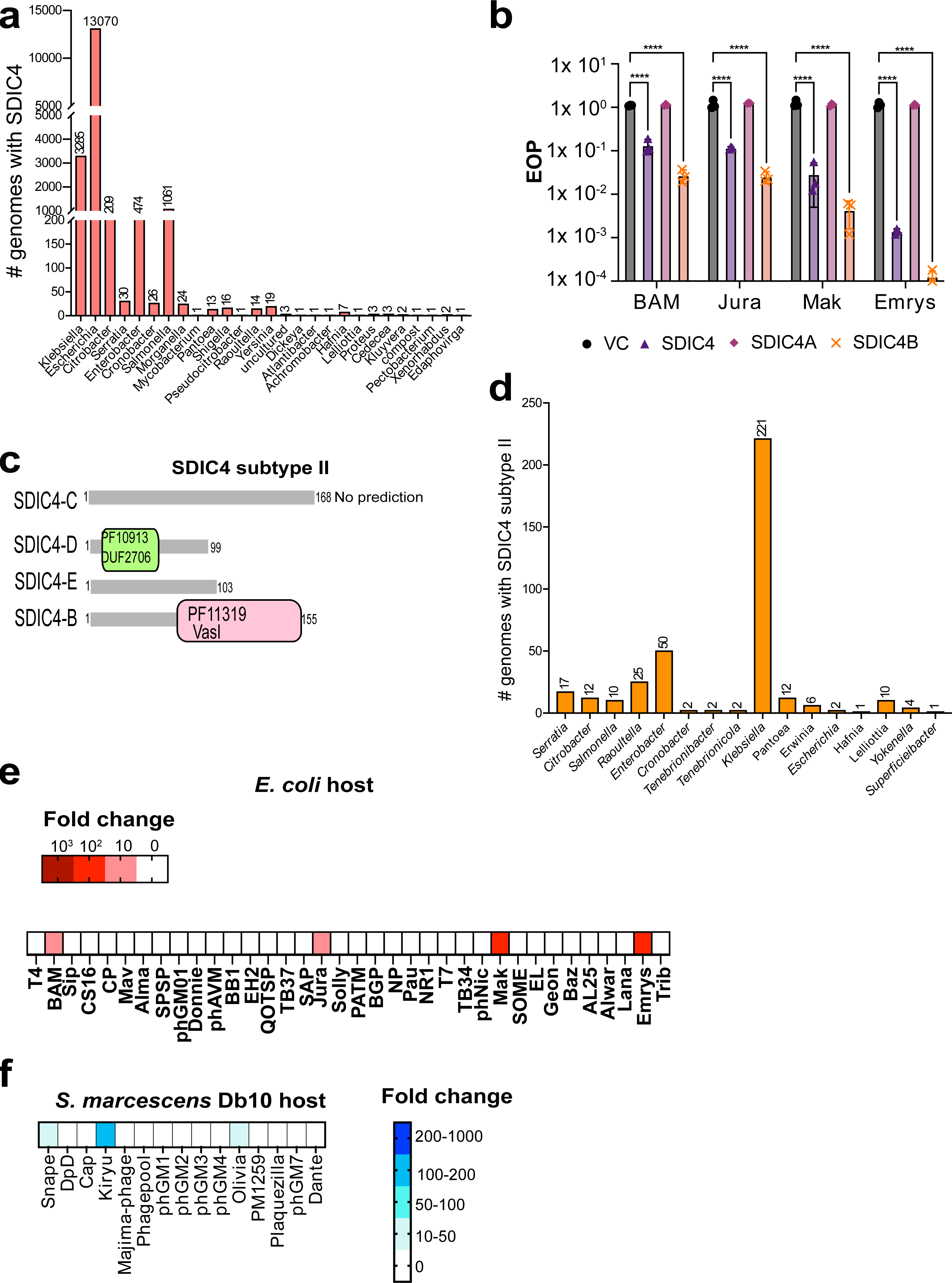
Proteins with VasI-like fold define two SDIC4 subtypes. **(a)** Distribution of SDIC4 systems across different genera. **(b)** Efficiency of plating (EOP) measurement for *E. coli* MG1655 carrying empty vector (VC, pQE60-Tat) or the same plasmid encoding SDIC4, SDIC4A or SDIC4B when challenged with phages BAM, Jura, Mak and Emrys. Points show mean +/− SEM (n = 3 biological replicates). Statistical relevance was measured using one-way ANOVA with Dunnett’s multiple comparison test. No significance was detected, unless indicated (*p ≤ 0.05). **(c)**Schematic representation of HHPRED domain predictions for the second SDIC4 subtype. **(d)** Distribution of SDIC4 subtype II across different bacterial genera. **(e)** Evaluation of SDIC4 subtype II fold protection against a suite of coliphages when expressed in *E. coli* MG1655 (n= 3 biological replicates). **(f)** Evaluation of SDIC4 subtype II fold protection against a suite of *Serratia* phages when expressed in *S. marcescens* Db10 (n=3 biological replicates). For **e-f**, fold protection was calculated as above.

We next set to explore the role that the single components SDIC4A and SDIC4B play in the defence process, using phages BAM, Jura, Mak and Emrys. We consistently observed that, for all tested phages, SDIC4B elicits higher levels of protection compared to the full-length locus. Additionally, SDIC4A cannot prevent phage infection (Figure 5b), demonstrating that SDIC4B is central to the phage defence mechanism of SDIC4.

Our FlaGs analysis further revealed a second operon, containing SDIC4B and associated with BrxR, which defines an additional subtype of SDIC4—SDIC4 subtype II (Figure 5c, Figure S9, Table S23). SDIC4 subtype II comprises four genes (SDIC4C-SDIC4D-SDIC4E-SDIC4B) and is less widespread, found in only 377 genomes in GenBank (Figure 5d and Table S25). It is primarily enriched in *Klebsiella* spp. (Figure 5d and Table S25). Except for SDIC4B, protein domain prediction was only possible for SDIC4E, showing the presence of a DUF2706 (Figure 5c). When expressed in *E. coli* and *S. marcescens,* SDIC4 subtype II elicits protection against a similar spectrum of phages compared to SDIC4 subtype I, suggesting SDIC4B homologues may play a central role in driving specificity to phage types (Figure 5d-e).

### SDIC1 mechanism of phage defence depends on SDIC1A

The increasing number of discovered and characterized anti-phage systems reveals a conserved tendency of shared defence domains^13^. These domains can be assembled in various configurations, contributing to a broad and diverse defence arsenal. TIR domains exemplify this trend, as they are associated or fused with various other defence proteins, forming distinct anti-phage systems and subtypes with diversified specificity and mechanisms of phage inhibition ^11,12,14,43^.

To better understand where SDIC1 fits within the repertoire of TIR-harbouring anti-phage systems, we verified that SDIC1A shares no sequence similarity with known TIR-containing defence proteins through pairwise comparisons. Furthermore, in line with PADLOC predictions, SDIC1 is not identified as a known anti-phage system using Defense-finder (Table S26)^44^. With the employment of Alphafold2 and Foldseek ^45^, we found that the TIR domain of SDIC1A exhibits a high degree of structural similarity to its closest homologue, the human TIRAP adaptor ^46^, while no matching fold was found for its C-terminal domain (Figure 6a and Figure S10).

**Figure 6.**
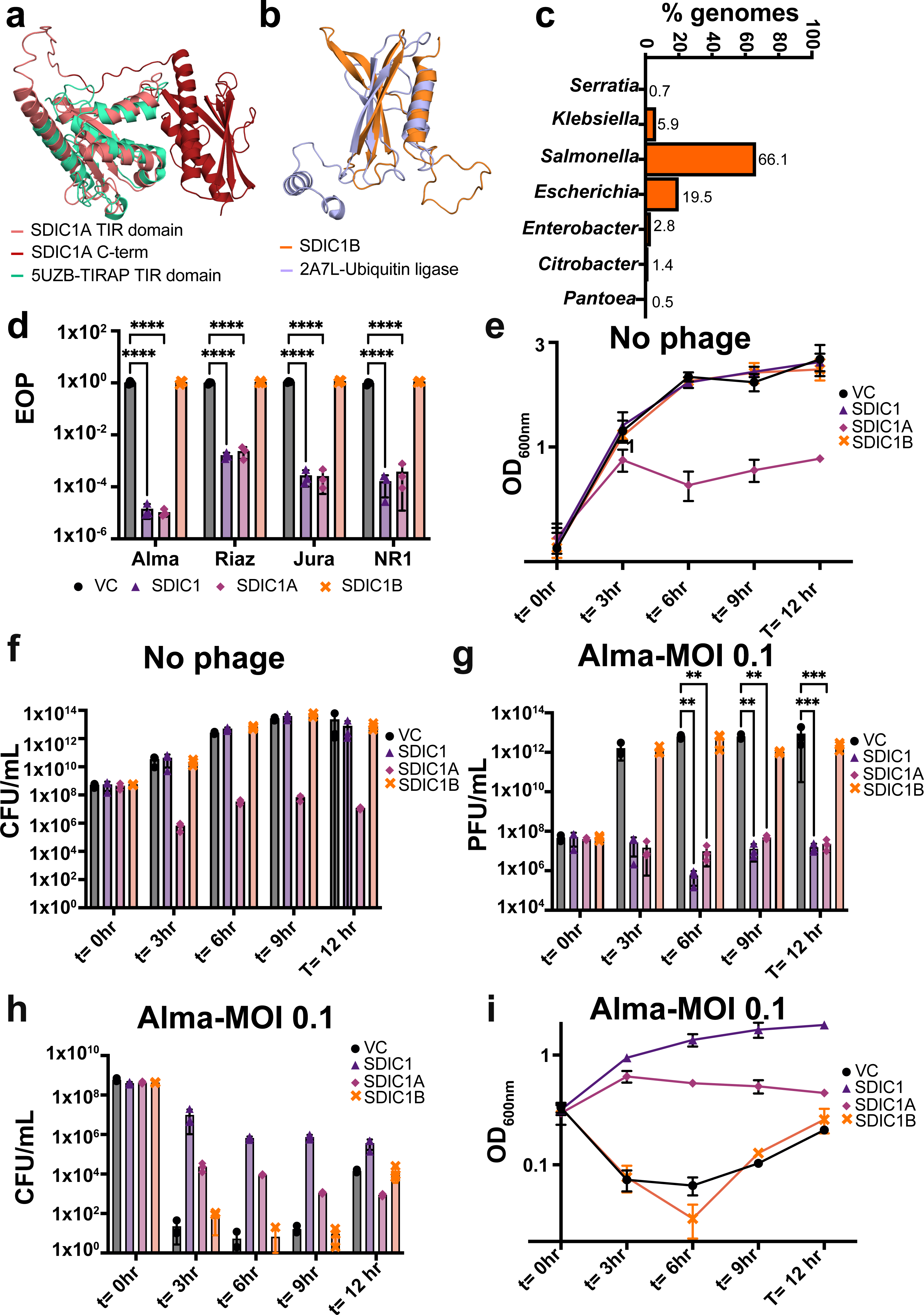
SDIC1 elicits protection against phages through population-wide immunity. **(a)** Pymol structural alignment (RMSD=2.67 Å) between the Alphafold2 model of SDIC1A and its closest structural homologue, the TIR-containing human protein TIRAP (PDBID: 5UZB). **(b)** Pymol structural alignment (RMSD=4.1 Å) between the Alphafold2 model of SDIC1B and a putative human ubiquitin ligase (PDBID: 2A7L). Structural homologues in **a-b** were obtained with Foldseek. **(c)** Distribution of SDIC1 across strains. **(d)**Efficiency of plating (EOP) measurement of *E. coli* MG1655 expressing empty vector(VC, pQE60-Tat) or the same vector encoding SDIC1, SDIC1A or SDIC1B when challenged with phages as reported in panel **c**. **(e-f)** Growth rate (OD_600nm_) **(e)** and cell counts (CFU/mL) **(f)** of *E. coli* MG1655 expressing empty vector (VC, pQE60-Tat) or the same vector encoding SDIC1, SDIC1A or SDIC1B in absence of phage infection. **(g-i)** *E. coli* MG1655 carrying empty vector, SDIC1, SDIC1A or SDIC1B were infected with Alma at MOI 0.1 and the released phages (titre, PFU/mL) **(g)**, cell counts (CFU/mL), **(h)** and growth rate (OD_600nm_)**(i)** was measured over the course of 12 hr post-infection. Statistical analysis for panel **c** and **f** was performed with GraphPad using one-way ANOVA with Dunnett’s multiple comparison test. No significance was detected, unless indicated (*p ≤ 0.05).

The closest homologue of the predicted structure of SDICB is instead a human hypothetical ubiquitin-conjugating enzyme. Despite a higher RMSD value (4.1 Å), these two proteins show a good degree of structural overlap (Figure 6b). Whilst both ubiquitin ligases and TIR domains have been associated with phage defence, the combination of the two has yet to be reported^7,8,12^.

Thus, we concluded SDIC1 represents a novel TIR-containing anti-phage system and we set to explore the role of the single components in phage defence. Homologues of SDIC1 are found in ∼2000 genomes and are particularly enriched in *Salmonella* and *Escherichia* genera. (Figure 6c and Table S27). When we compared the EOP of several phages on strains expressing full-length SDIC1, SDIC1A or SDIC1B only, we found that SDIC1A is necessary to elicit protection against phages, whereas SDIC1B is dispensable (Figure 6d).

We next assessed the progression of phage Alma infection over a 12-hour period, measuring PFU/mL, CFU/mL and OD_600nm_. In absence of a phage threat, cells harbouring full-length SDIC1 exhibit a normal growth rate and CFU/mL over time, whereas SDIC1A-only cells show impaired growth on both solid and liquid media (Figure 6e-f). In presence of a phage threat, cells containing an empty vector or SDIC1B-only encounter a rapid culture collapse and release a high number of phages (Figure 6g-i). Cells carrying SDIC1 retain an unaltered growth rate in liquid, comparable to that of non-infected cells, but decreased cell counts (Figure 6g-h). Nevertheless, they release a significantly lower number of phages compared to vector control over the course of 12 hrs (Figure 6h), indicating SDIC1 operates through an abortive infection phenotype or population-wide immunity. In the case of SDIC1A-only, the growth rate and CFU counts remains highly impaired, however the number of released phages is also lower than vector control (Figure 6g-h).

Overall our data show that, as other TIR-containing defence systems, SDIC1 acts through population-wide immunity, impairing the fitness of infected cells to protect the bacterial population. Our data further suggest that whilst SDIC1A mediates this phenotype, SDIC1B’s role may consist in alleviating SDIC1A toxic effect.

### SDIC1A confers population-wide immunity through membrane depolarisation

Our structural prediction of SDIC1A shows two clear and separate domains: an N-terminal TIR domain and a C-terminal domain with an unknown function. To determine which domain directly mediates SDIC1A’s anti-phage activity, we generated SDIC1A_ΔTIR_ (deletion of aa 1-103) and SDIC1A_ΔC-term_ (deletion of aa104-323) in presence or absence of SDIC1B. Deletion of each SDIC1A domain abolished anti-phage activity and SDIC1A-mediated toxicity on bacterial growth (Figure 7a-b), indicating both SDIC1A domains are central for its activity.

**Figure 7.**
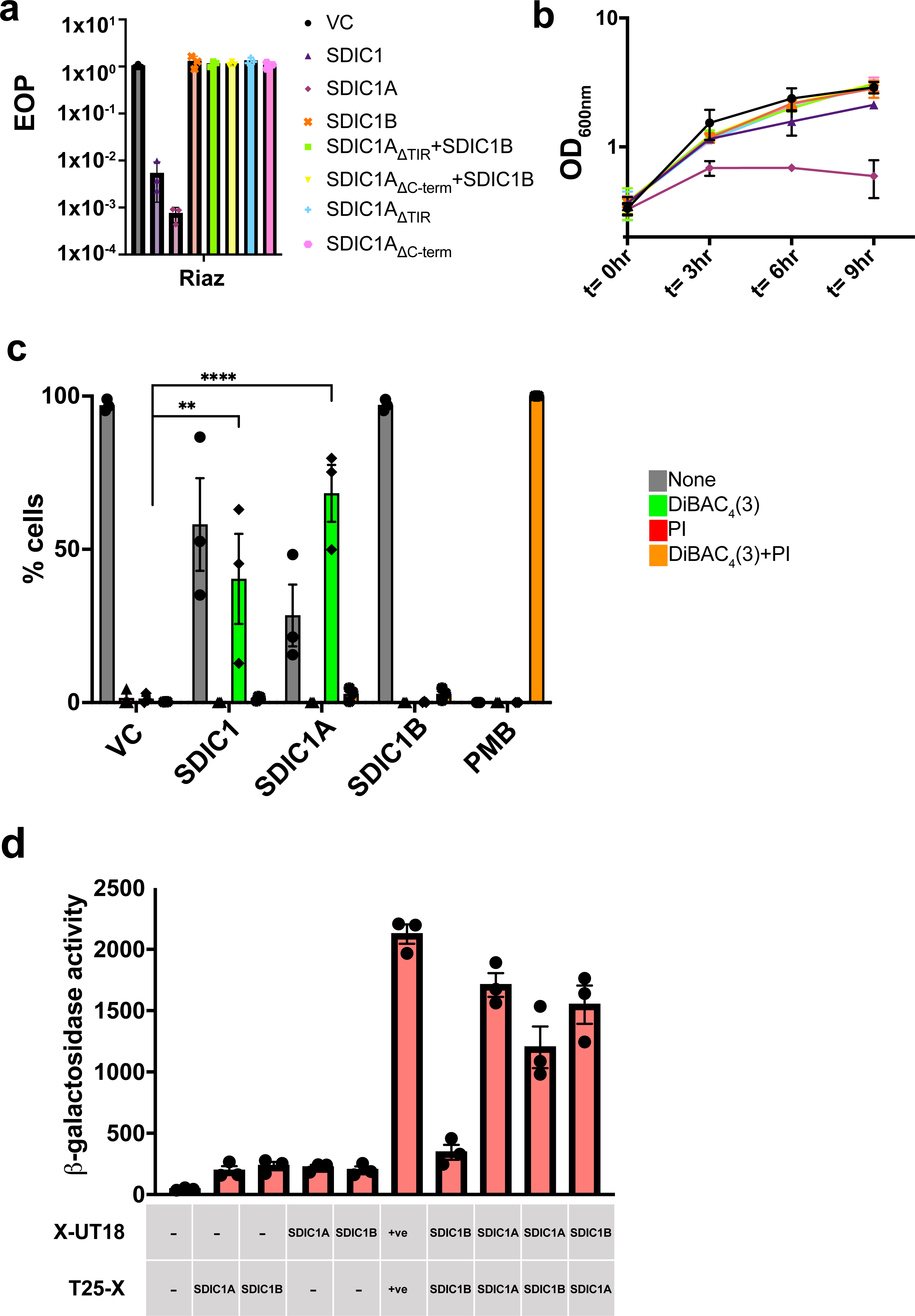
SDIC1 causes membrane depolarisation. **(a)** Efficiency of plating (EOP) measurement of *E. coli* MG1655 expressing empty vector(VC, pQE60-Tat) or the same vector encoding SDIC1, SDIC1A, SDIC1B, SDIC1A_ΔTIR_+SDIC1B, SDIC1A_ΔC-term_+SDIC1B, SDIC1A_ΔTIR_ or SDIC1A_ΔC-term_ when challenged with phage Riaz. Points show mean +/− SEM (n = 3 biological replicates). **(b)** Growth rate (OD_600nm_) for strains as in panel **a** in absence of phage infection. **(c)** *E. coli* MG1655 expressing empty vector (VC, pQE60-Tat) or the same vector encoding SDIC1, SDIC1A or SDIC1B were stained with DiBAC_4_(3) or propidium iodide (PI) and analysed by flow cytometry. The percentage of cells that were depolarised (DiBAC_4_(3) positive), permeabilised (PI positive), or both is reported. Points show mean +/− SEM (n = 3 biological replicates). Polymyxin B (PMB) was used as the depolarisation control. **(d)** Bacterial two-hybrid assay to detect interactions between SDIC1A and SDIC1B. Negative controls were provided by the empty vectors, whereas the positive control was provided by ShdB ^52^(+ve). Interactions were quantified with β-galactosidase activity assays, expressed as Δ405/min/ml/OD_600_. Points show mean +/− SEM (n = 3 biological replicates).

We next set to determine the mechanism by which SDIC1 causes population-wide immunity. Examination with the DNA-specific stain 4′,6-diamidino-2-phenylindole (DAPI) indicated no damage to the bacterial chromosome (Figure S10c). Next, cells were stained using DiBAC_4_(3) and propidium iodide (PI). DiBAC_4_(3) is a negatively-charged dye that can only enter depolarised cells, resulting in green fluorescence. PI is unable to penetrate intact cells, but can cross highly-damaged membranes, emitting red fluorescence. Depolarised cells are detectable in both SDIC1 and SDIC1A populations, but the percentage of DiBAC_4_(3)-positive cells is higher for SDIC1A only, in agreement with what observed in growth experiments (Figure 7c, Figure S10d). TIR domains often provide protection against phages through NAD^+^ degradation^47^. NAD^+^ depletion is normally accompanied by the loss of membrane potential ^56^; thus, we wondered if SDIC1-mediated depolarisation was a secondary effect of NAD^+^ degradation caused by the SDIC1A TIR domain. Overall, while we observed no significant changes in whole-cell levels of NAD^+^, there was a decrease in NAD^+^ in SDIC1A-only cells compared to the control (Figure S10e). Nevertheless, this difference was not found to be statistically significant (Figure S10e). Thus, we were unable to fully discern whether SDIC1-mediated depolarisation is caused by NAD^+^ depletion or whether slight changes of NAD^+^ levels are a secondary consequence of the depolarisation. Lastly, given the increased toxicity of SDIC1A expression in absence of SDIC1B, we used a bacterial two-hybrid assay to investigate their possible interaction^48^. This yielded a clear interaction between SDIC1A and SDIC1B (Figure 7d) and a self-interaction between SDIC1A monomers (Figure 7d).

Taken together, our data show that the mechanism of population immunity mediated by SDIC1 requires both SDIC1A domains and includes the disruption of membrane potential. Furthermore, we demonstrate that oligomerisation is a part of the SDIC1A mechanism, and that SDIC1B-mediated inhibition of its toxicity might involve an interaction between the two proteins.

## DISCUSSION

An increasing amount of evidence has shown that anti-phage systems are clustered in ‘defence islands’ and very often localised on the mobilome of bacteria, likely to facilitate their rapid transfer during bacteria-phages arms race ^3,4,15–17,19,49^.

More recently, a less conservative analysis of defence islands content evidenced that these regions often show a high degree of plasticity and that anti-phage systems are frequently neighbouring other resistance factors, such as stress genes, antibiotic resistance, and virulence factors ^20,50,51^. Importantly, in *Vibrio* spp., Mahata *et al.* highlighted GMT mobile islands as the first example of co-localisation of anti-phage systems with T6SS-dependent effectors^51^. In this study, through the analysis of *Serratia* genomes, we show that this phenomenon is widespread to other defence islands and encompasses several types of offensive and anti-bacterial tools, including colicins, bacteriocins and Cdi systems (Figure 1,2 and Figure S3). Our analysis revealed that, within the three identified islands, the anti-phage system cargo varies significantly across *Serratia* species. However, there is a predisposition to acquire a specific type of anti-bacterial or virulence effector for each island. The SpeG-PstB island stands out as the only exception, predominantly harbouring a Thoeris I system, a haemolysin-like toxin system, or both (Figure 1).

The *Serratia* LptG-YjiA we report partially overlaps a previously identified genomic hotspot in *E. coli* strains^17^. In this study, the multi-drug efflux transporter YjiO was identified as the flanking gene together with YjiA^17^. Our search demonstrates that while in *Serratia* spp. and *E. albertii,* LptG-YjiA islands do not encompass YjiO, in several strains of *Salmonella, Citrobacter, Shigella* and *E. coli* this genomic hotspot is predominantly organised as LptG-YjiO-YjiA and that anti-phage systems, anti-bacterial effectors and virulence factors can be inserted indiscriminately between LptG-YjiA or YjiO-YjiA. *E. coli* strains found to carry LptG-YjiA islands in our study are mostly distinct from those reported to carry a YjiO-YjiA hotspot ^17^, implying a high likelihood of recombination events in these hotspots and high variability even across strains. The frequent recombination events will ultimately result in variability of the flanking genes that delimit defence islands, even across the same genera, as we observe for *E. coli* and *E. albertii,* or across *Salmonella* serovars, as reported recently^20^. It is plausible that, even within *E. coli* strains, some defence islands are organised as LptG-YjiO-YjiA, while others are organised as YjiO-YjiA, possibly due to recombination events. This variability should be considered as efforts increase to identify defence islands flanking genes, aiming to facilitate the discovery of novel anti-phage systems.

A high percentage of genes found in the *Serratia* LptG-YjiA islands had either no predicted function, or they carried known defence domains that were, however, not classified by PADLOC as known defence systems. We show that a closer look at these proteins will enable to define distinct classes or types of anti-phage systems, wherein known defence proteins have been re-shuffled to assemble a highly diversified arsenal of anti-phage systems, as in the case of SDIC1, SDIC2 and SDIC3. This tendency is consistent with what has been extensively reported for many known anti-phage systems, which often share similar defence domains, assembled in different combinations^3,4,13,27^. Importantly, in our study, we demonstrate that our focused investigation of *Serratia* islands’ cargo led to the discovery of anti-phage systems (SDIC1 and SDIC4) present across many species, emphasising the scalability of genus-focused studies (Figure 5-6).

Our investigation of the LptG-YjiA islands’ cargo further led to the intriguing revelation that the anti-phage system SDIC4, found in two different subtypes, co-opted the same fold as a poorly-characterised predicted T6SS-related protein for phage defence (Figure 4-5). This discovery aligns with a growing body of reports documenting domains of diverse origins, including housekeeping proteins, being co-opted for defence against phages^3,4,13,36,52^. Furthermore, it corresponds to observed overlaps between toxin-antitoxin systems and anti-phage defence mechanisms ^36,53–55^. It is plausible to speculate that this evolutionary or functional connection may extend to other anti-bacterial effectors or domains of different nature^3,4,36,53,54^.

Significantly, in the case of SDIC1, we uncover a new type of anti-phage system characterised by the presence of a TIR domain and its association with a putative ubiquitin ligase—a combination not previously documented in other anti-phage systems (Figure 6). SDIC1A oligomerises and mediates phage inhibition by reducing the survival of infected cells, likely through disruption of membrane potential (Figure 7). In our study, while depolarisation caused by SDIC1A was consistent, we only observed a slight change in NAD^+^ levels in SDIC1A-only cells. Whilst these observations point to depolarisation being the primary effect of SDIC1A intoxication, we were not able to fully discriminate between the two often connected phenotypes^56^. The observation that both the C-terminal domain and the TIR domain of SDIC1A are necessary for its activity further points to depolarisation as the primary consequence of the SDIC1A-mediated mechanism (Fig.7a-b). Nevertheless, future biochemical investigation of purified SDIC1A will establish whether NAD^+^ degradation or depolarisation is the primary phenotype.

SDIC1B exhibits distant similarity to a putative ubiquitin ligase and although it is not essential for eliciting phage protection, it plays a central role in reducing the toxicity of SDIC1A, likely through their interaction (Figure 6-7). Unlike the Bil and CBASS systems, where ubiquitylation-like modifications directly impede phage assembly or the release of virulent phage particles^7,8,57^, SDIC1B’s primary function seems to revolve around alleviation of SDIC1A levels of toxicity. A regulative role for ubiquitination in TIR receptors signalling in eukaryotes has been previously documented^58^. Future investigations will aim to uncover the mechanism by which SDIC1B can modulate SDIC1A-mediated toxicity and discern the specific role it plays in the defence mechanism.

In future, the analysis of existing and new defence islands is poised to reveal additional instances of co-accumulation of anti-phage systems, virulence factors, and anti-bacterial effectors. Such exploration holds the potential to identify the evolutionary and environmental factors influencing the acquisition of specific anti-bacterial, virulence and anti-phage modules or their unique combinations within each island. These genomic islands additionally represent a valuable resource that promises to uncover novel anti-bacterial effectors, anti-phage systems, and virulence factors with new and exciting biological mechanisms.

## MATERIAL AND METHODS

### Bacterial strains, plasmids and culture conditions

Expression of novel anti-phage systems in *E. coli* was performed using the MG1655 strain. MG1655 cells were grown at 37°C on either solid media or liquid culture, shaking at 200 rpm. Growth in liquid media was performed using the LB Miller Broth (Formedium) whereas for solid media, LB was supplemented with 1.5% (w/v) or with 0.35% (w/v) agar to obtain solid or soft agar, respectively.

Expression in *S. marcescens* was performed using the *S. marcescens* Db10 strain and cells were grown at 30°C LB Lennox Broth (Formedium). Agar was added at 1.5% (w/v) or at 0.5% (w/v) to obtain solid or soft agar plates. When required, LB was supplemented with ampicillin (Amp, 50 μg/mL) for plasmid selection in *E. coli* or carbenicillin (200 μg/mL) for selection in *S. marcescens.* Strains and plasmids used in this study are listed in Table S28

### Phage propagation and lysate preparation

Coliphages used were derived from the Durham collection, except for phages reported in Table S28. *Serratia* phages are reported in table Table S28.

Coliphages lysates were propagated in *E*. *coli* DH5α and prepared in phage buffer (10 mM Tris–HCl pH 7.4, 10 mM MgSO4, 0.1% gelatin). *Serratia* phages were instead propagated in *S. marcescens* Db10 and stored in phage buffer.

Neat lysates or their 10x fold dilutions were added to 200 μL of *E*. *coli* DH5α or *S. marcescens* Db10 and incubated 5 min at room temperature. Five mL of soft agar were then added and the mixture poured on LB agar plates. Plates were then incubated overnight at 37 °C for *E*. *coli* and 30 °C for *S. marcescens.* Following incubation, top agar containing confluent plaques was scraped off and added to 3 mL of phage buffer and 500 μL of chloroform. Samples were mixed for 2 minutes by vortexing, incubated at 4°C for 30 min and subsequently centrifuged at 4000 x g for 20 min. The supernatant was collected and added to 100 μL/mL (v/v) of chloroform for storage.

### DNA manipulation and cloning

Plasmids (Table S28) were synthetised through Genscript (https://www.genscript.com/) unless stated otherwise. Inserts of interest were cloned by Genscript in a low copy vector (pQE60) modified to harbour a constitutive Tat promoter to replace its native T5 promoter. For insert cloning, plasmid backbones and inserts were amplified using Verifi hot start polymerase (PCR Biosystems). PCR products and plasmids were purified using a Zymo DNA Clean-up and Concentration and a Zymo Zippy plasmid kit, respectively (Cambridge Biosciences). Overlapping primers for amplification were designed using the NEBuilder assembly tool (https://nebuilderv1.neb.com/) (Table S29). Plasmids and inserts were assembled using NEBuilder HiFi DNA Assembly (NEB), followed by incubation at 50°C for 30 min. NEBuilder reactions were treated with 1 μL DpnI for 30 min at 37 °C and subsequently transformed in chemically competent DH5α. Deletions of single genes in plasmid pGM255 were performed using the KLD enzyme mix (NEB).

### Efficiency of plating measurement and fold protection calculation

To measure the efficiency of plating (EOP), *E*. *coli* MG1655 or *S. marcescens* Db10 strains harbouring an empty vector or constructs encoding anti-phage systems of interest were grown to exponential phage (OD_600nm_=0.6). Ten microliters of phage lysates was added to 200 μL of each bacterial culture and incubated 5 min at room temperature. Five mL of soft agar was added to each culture and poured onto LB agar plates. As a control strain, plasmid-free MG1655 or plasmid-free *S. marcescens* Db10 were used. EOP was measured as the PFU mL^-1^ of a test strain divided by the number of PFU/mL of a plasmid-free control strain. Fold protection was calculated as the ratio between the EOP value of empty vector and the EOP value of each candidate system.

### Measurement of PFU/mL, CFU/mL and growth rate

For SDIC1, *E*. *coli* MG1655 harbouring empty vector, full-length SDIC1 or SDIC1A or SDIC1B were grown in LB with ampicillin to an OD_600nm_ of ∼ 0.6. Cultures were infected with phage Pau at MOI of 0.1. An aliquot of each culture was collected at t = 0 hr, t = 2 hr, t = 4 hr, t = 6 hr, t = 8 hr, t = 10 hr and t = 12 hr. Each aliquot was used to measure OD_600nm,_ CFU/mL and measure phage titre (PFU/mL). Measurement of the phage titre at each timepoint was obtained by plating onto *E*. *coli* DH5α top lawns.

### Bacterial two hybrid assay

Bacterial two-hybrid assays were performed as previously reported ^48,59^. SDIC1A and SDIC1B were fused to two fragments of the adenylate cyclase encoded on plasmids pUT18 and pT25 (Table S28-29). Derived plasmids were co-transformed in *E. coli* MG1655 Δ*cyaA* and plated on MacConkey agar plates with Amp or chloramphenicol (Cml, 25 μg/mL) and 1% maltose. Plates were incubated for 48 hrs at 30°C and subsequent 24 hrs at 37°C. Positive interactions were identified as dark red colonies. For quantification, a β-galactosidase assay was performed on 3 biological replicates. Cells were normalised to the same OD_600nm_, permeabilised with toluene and resuspended in a buffer containing 60 mM Na_2_HPO_4_, 40 mM NaH_2_PO_4_, 10 mM KCl, 1 mM MgSO_4_, 36 mM β-mercaptoethanol and 1.1 mg/mL o-nitrophenyl β-galactoside. The reaction was quantified as the rate of increase of A_405_ (ΔA_405_/min) at 37°C using a CLARIOstar Plus Microplate Reader (BMG Labtech).

### Flow cytometry assessment of membrane depolarisation

*E. coli* MG1655 carrying empty vector or the same vector carrying full-length SDIC1, SDIC1A or SDIC1B were grown in LB supplemented with Amp in a final volume of 5 mL. As SDIC1A-only cells showed decreased growth rate, all cultures were recovered when empty vector cells reached an OD_600nm_= 1, washed in LB and then normalised to an OD_600nm_= 0.3 (Corresponding to the lowest OD_600nm_ of SDIC1A cells). DiBAC_4_(3) (Bis-[1,3-Dibutylbarbituric Acid] Trimethine Oxonol; Thermo) and propidium iodide (Thermo) were added at a final concentration of 10 μM and 1 μM, respectively. Cells were incubated in the dark for 20 min at 37°C. As a control, cells were treated with 2 μg/mL polymyxin B (Merck). Following staining, cells were analysed on a Attune NxT Acoustic Focusing Cytometer with filters BL1 (Ex = 488 nm, Em = 530/30 nm) and YL1 (Ex = 561 nm, Em = 585/16 nm).

### Whole cells NADase assay

*E. coli* MG1655 cells carry empty vector, SDIC1 or its mutants were grown and normalised as above. NAD^+^ and NADH levels were measured using the EnzyChrom™ NAD/NADH Assay Kit (Universal Biologicals) following the manufacturer instructions. Briefly, for NAD^+^ assessment cells were washed in ice-cold PBS, resuspended in NAD^+^ specific buffer, and incubated at 60 °C for 5 min. Following incubation, cell suspensions were neutralised with NADH-specific buffer and with assay buffer. A calibration curve was built using the kit-provided NAD^+^ standard. 40 μL of each cell suspension or standard was added to a 96-well plate (Sarstedt). For each well, 80 μL of working reagent (containing assay Buffer, Enzyme A, Enzyme B, Lactate and MTT) were added and absorbance (A _565nm_) was recorded after 0 min and 30 min.

### Fluorescence microscopy

*E. coli* MG1655 carrying empty vector or the same vector carrying full-length SDIC1, SDIC1A or SDIC1B were grown until empty vector cells reached an OD_600nm_= 1. Cells were stained with 4′,6-diamidino-2-phenylindole (DAPI) at a final concentration of 5 μg /mL. Cells mixed with DAPI were incubated at 37°C for 15 min and then 1 μL of each culture was transferred on a microscope slide with a pad of 1% UltraPure agarose (Invitrogen) in H_2_O. Images were collected on a Zeiss LSM980 Microscope equipped with Widefield Camera Axiocam 705 mono and Light Source Colibri 5 Type RGB-UV - 4-channel fluorescence light source with integrated control unit and a Plan Apochromat 63x objective.

### Anti-phage systems predictions

Anti-phage systems were predicted using PADLOC v2.0.0 with PADLOC database v2.0.0^27^ unless stated otherwise. Additionally, Defense Finder v1.2.0 was also run BSAC genomes to confirm that SDIC1 represents a new instance of anti-phage system harbouring a TIR domain.

### Identification of defence islands and boundaries in the BSAC strains collection

The coordinates of the genomic neighbourhood of each anti-phage system predicted by PADLOC was retrieved with the use of pyfaidx v0.7.2.1 and BEDtools suite v2.30.0 ^60,61^, using the ‘slop’ function. Genomic regions were then retrieved using efetch from the entrez utilities ^62^. Proteins sequences in the genomic neighbourhoods were clustered with MMseqs2 linclust using the following parameters: cov-mode=0, c=0.5 and min-seq-id=0.6^28^. Protein clusters were curated manually to identify those that were more frequently associated with anti-phage systems in BSAC strains and thus, to identify defence island boundaries. Genomic islands were scored as defence-enriched loci only if, upon scaling their search to all *Serratia* genomes, they carried known defence systems in > 10% instances.

Gene clusters were visualised and annotated using RStudio (R v4.3.1), ggplot2 and gggenes v0.5.1^63^. Genbank files were converted to a gggenes-compatible format using biopython SeqIO and SeqFeature^64^. Predictions parallelisation was obtained using GNU parallel ^65^. Custom scripts used for analysis and to generate figures are found at: https://github.com/GM110Z/Serratia_multiconflict.

### Analysis of gene clusters and genomic islands

To find defence islands in all *Serratia* genomes or in other species, the protein sequence of the two boundaries was searched against Genbank using the program cblaster v1.3.12^29^ with the following parameters: -min id=30 and -min cov = 50. An intergenic space of >50,000 bp was allowed.

Cblaster was additionally used with -min id=30 and -min cov = 50 and an intergenic space < 70 bp to identify SDIC1, SDIC4 subtype I and subtype II instances in Genbank genomes. Only operons present on the same contigs were allowed.

### Protein domain analysis of genomic islands

To characterise the non-defence content of LptG-YjiA islands in all *Serratia* genomes, proteins were clustered as above. One representative sequence for each cluster was first analysed using the SecReT6, AMRFinderPlus and Virulence Factor Database (VFDB)^33–35^. Comparisons with VFDB were performed with PSI-BLAST, including hits with bitscore>50. For hits with bitscore<50, these were only included if they had a e-value< 0.01 and if the alignment covered at least 70% of the query and subject sequence length. Proteins whose function could not be determined through these programs were further analysed using HHPRED against a PFAM and PDB database. Only hits with an e-value< 0.05 were included.

To investigate the anti-phage systems content of LptG-YjiA islands in other bacterial species, genomes were filtered using fastANI in order to eliminate duplicate or very closely related genomes ^66^. PADLOC was used as above to predict anti-phage systems in the representative genomes obtained. Non-defence proteins were analysed with VFDB as above.

For analysis of the cargo of the LysR-esterase hotspots HMMscan from the HMMER 3 suite was run against a local PFAM database^30^.

### Protein structure predictions

Protein structure prediction was performed using Alphafold2 with MMseqs2 (v2.2.4) through Colab notebooks. Obtained models were used as input in FoldSeek to find structural homologues (https://search.foldseek.com/search)^45,67^. Protein structures were aligned on Pymol with the cealign option.

### Computational analysis of the genomic neighbourhood of SDIC4B (VasI-like proteins) homologues

SDIC4B homologues were identified using PSI-BLAST, with a threshold of e-value=0.01. sequence identity = 30% and query coverage = 50%. The genomic neighbourhood of the obtained hits was retrieved and clustered using the FlaGs2.py program ^42^

## AUTHOR CONTRIBUTIONS

G.M conceptualised the study. T.C. and G.M. performed experiments. G.M and S.R.G performed the bioinformatic analysis. G.M., T.M., S.R.G and T.R.B analysed data. G.M. wrote the manuscript with input from T.C., S.R.G, and T.R.B.

## ACKNOWLEDGEMENTS

The authors thank Dr Emmanuele Severi for useful discussion on the use of the Tat promoter. The authors additionally thank Dr Franklin Nobrega and Prof Sarah Coulthurst for critical reading of the manuscript.

