## Supplementary-Figures for "Multi-conflict islands are a widespread trend within *Serratia* spp"

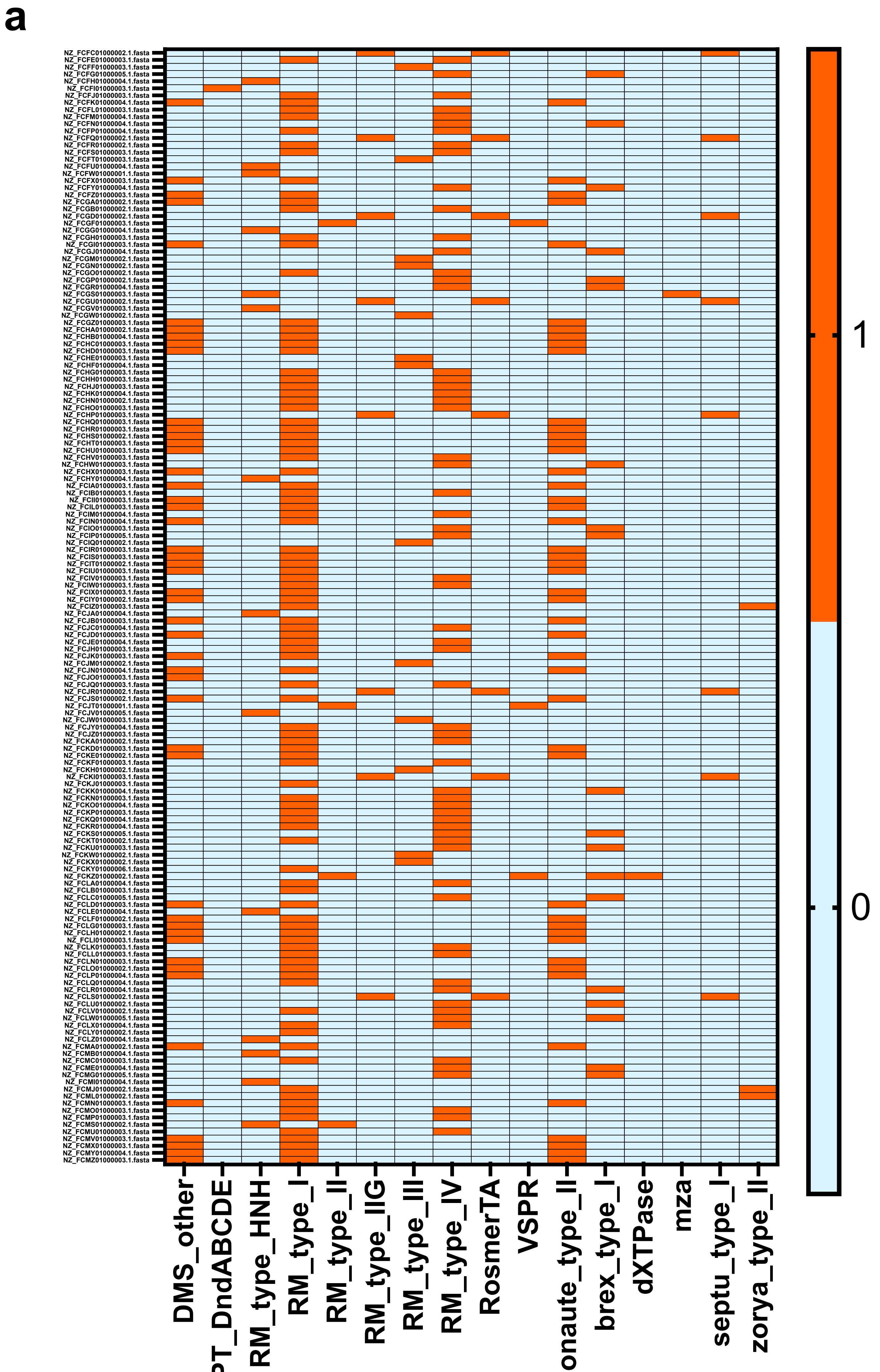
a
1
0
DMS_other
PT_DndABCDE
RM_type_HNH
RM_type_I
RM_type_II
RM_type_IIG
RM_type_III
RM_type_IV
RosmerTA
VSPR
onaute_type_II
brex_type_I
dXTPase
mza
septu_type_I
zorya_type_II

**b**

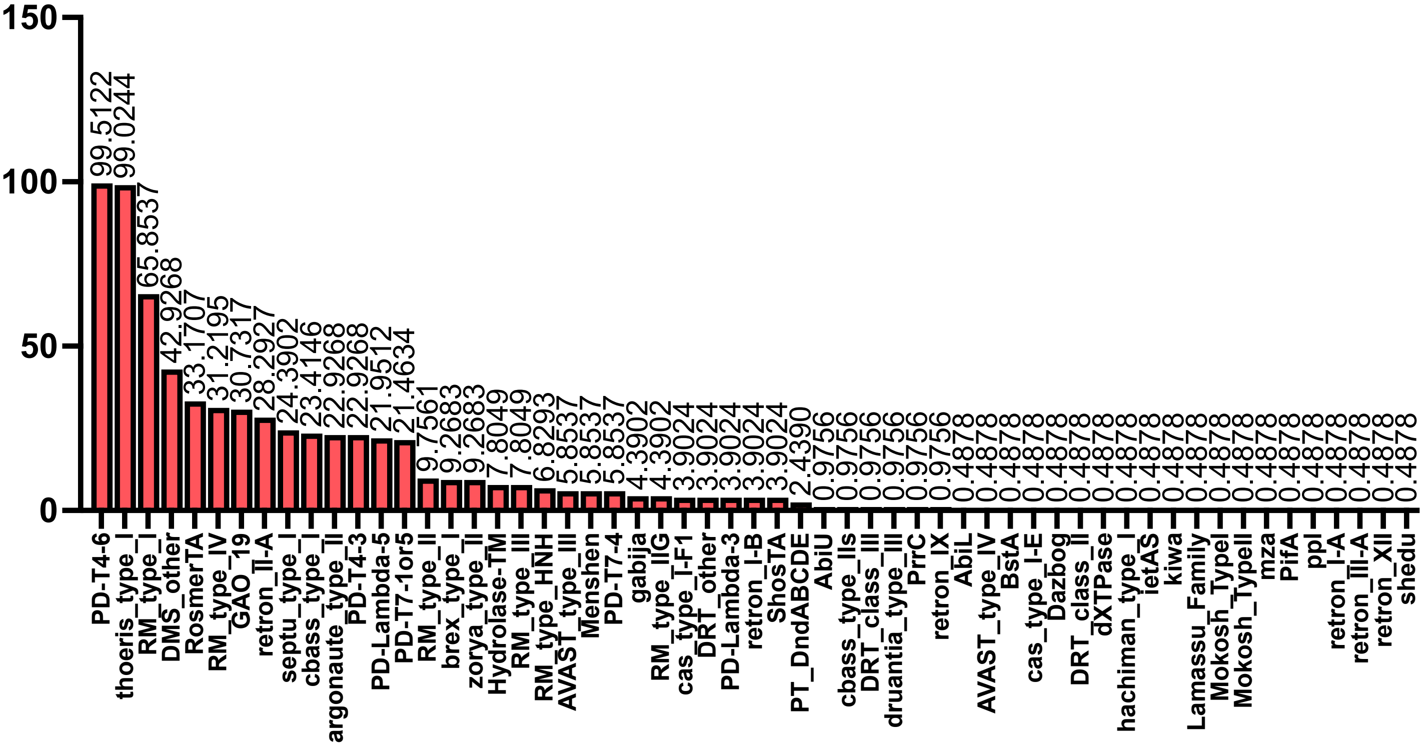


**Figure S1. Anti-phage system arsenal on LptG-YjiA islands of the BSAC collection. (a)** Heatmap showing presence-absence of anti-phage systems predicted by PADLOC in the *S. marcescens* strains of the BSAC collection. **(b)** Prevalence of anti-phage systems predicted by PADLOC in the *S. marcescens* strains of the BSAC collection.

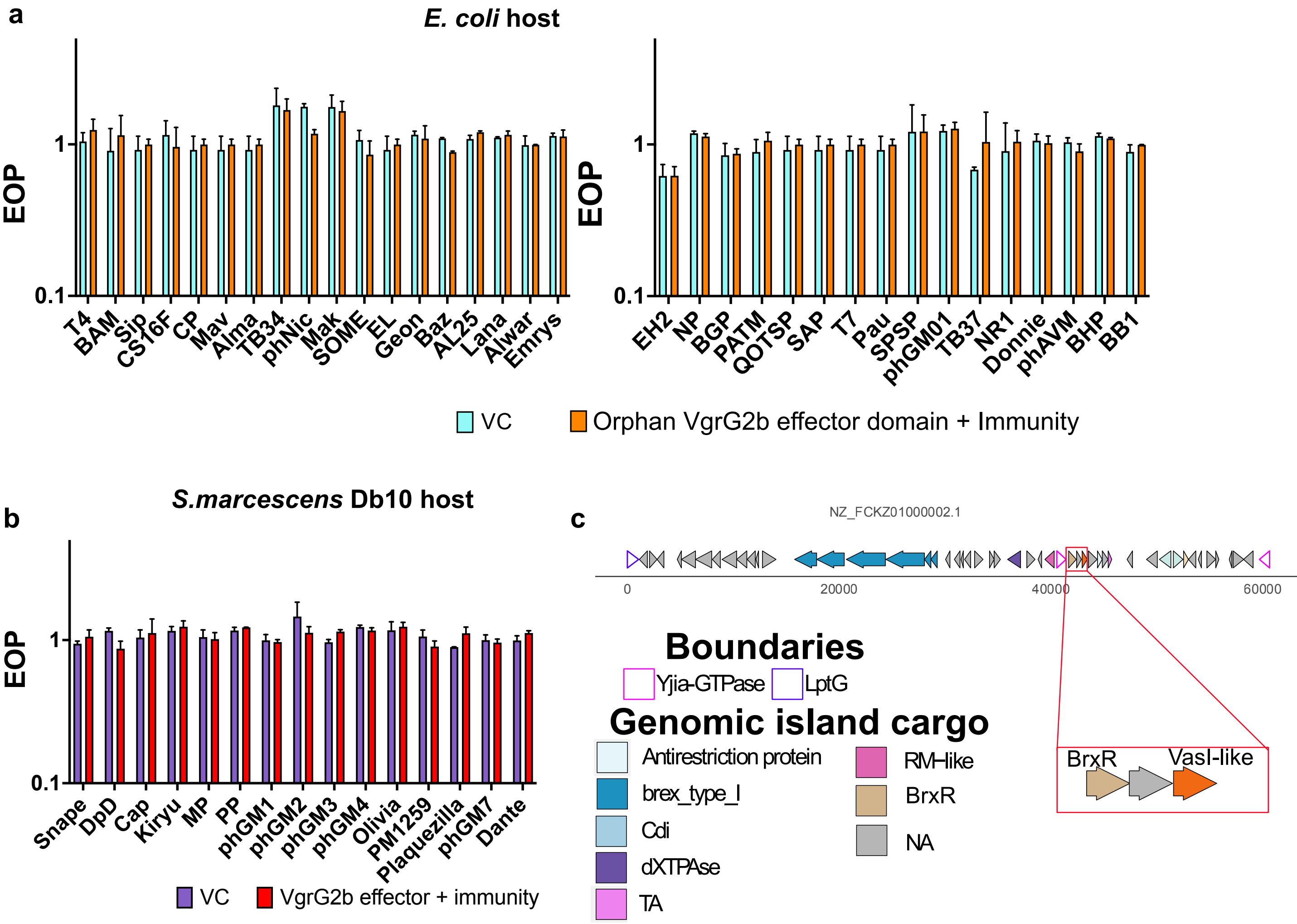


**Figure S2. The orphan VgrG2b toxic effector and its cognate immunity do not play a role in phage defence. (a)** Efficiency of plating (EOP) measurement for *E. coli* MG1655 carrying empty vector (VC, pQE60-Tat) or the same plasmid encoding the VgrG2b toxic effector-immunity pair (VgrG2b effector domain + immunity) when challenged with a panel of coliphages as shown in panel **a**. Efficiency of plating (EOP) measurement for *S. marcescens* Db10 harbouring empty vector (VC, pQE60-Tat) or the same plasmid encoding the VgrG2b toxic effector-immunity pair (VgrG2b effector domain + immunity) when challenged with a panel of *Serratia* phages as shown in panel b. For panel **a-b** Points show mean +/− SEM (n = 3 biological replicates). **(c)** Schematic representation of a LptG-YjiA island harbouring a VasI-like protein encoded next to BrxR.

a

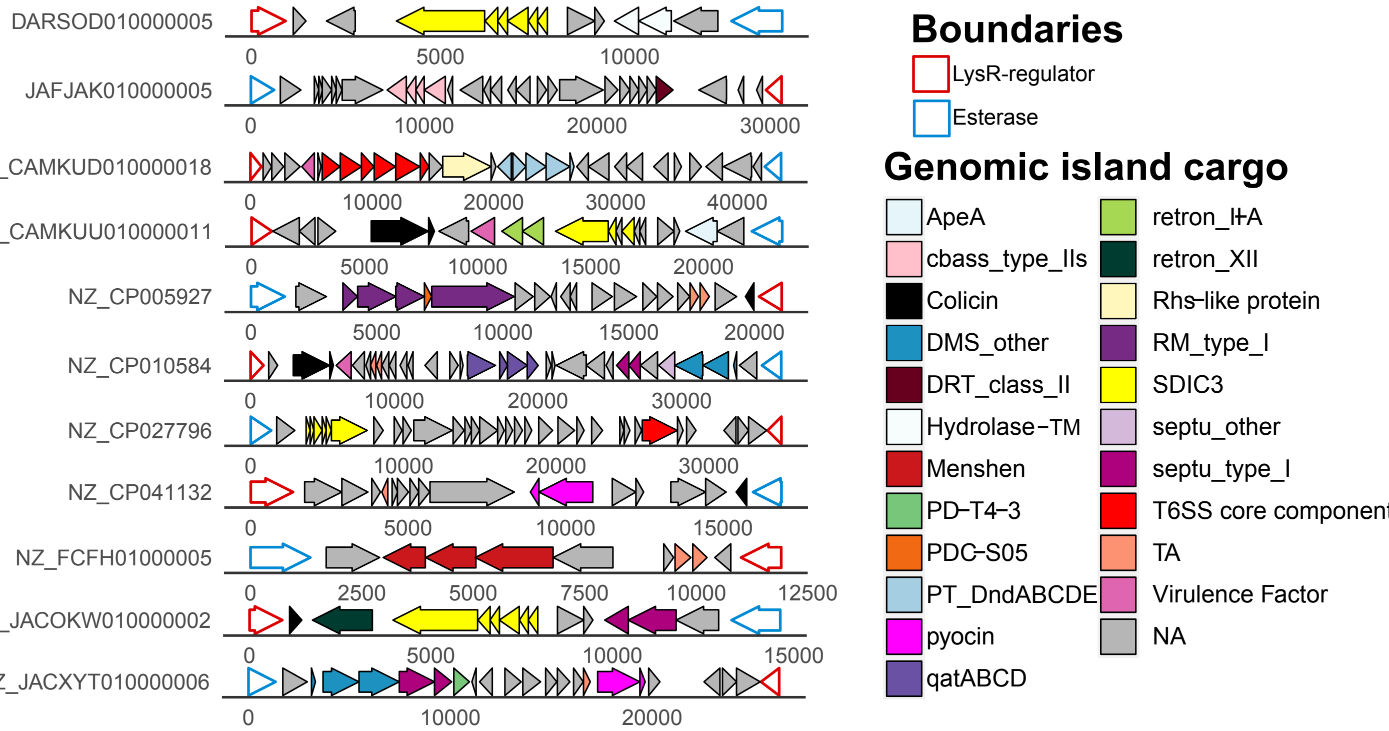


b

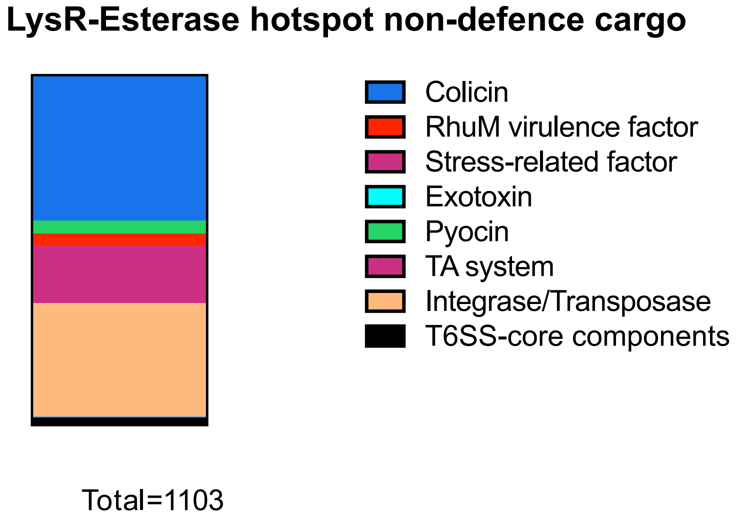


**Figure S3. LysR-esterase hotspots often carry colicins. (a)** Schematic representation of representative LysR-Esterase islands. Island boundaries are represented with a coloured outline. Anti-phage systems predicted by PADLOC, integrases, transposases, T6SS-dependent effectors/core components, colicins and pyocins are coloured according to the legend. **(b)** Abundance of the non-defence cargo of LysR-esterase hotspots.

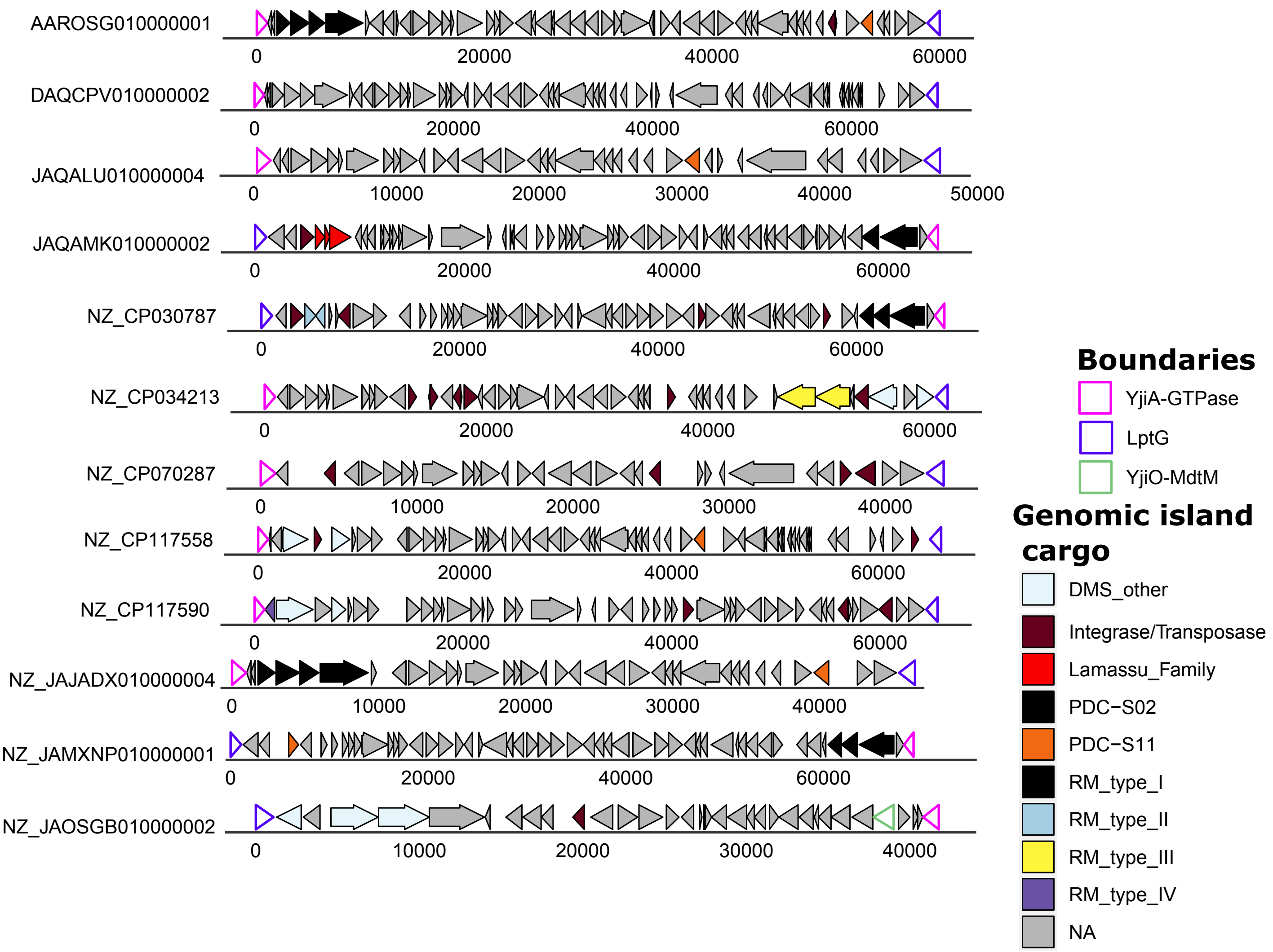


**Figure S4. LptG-YjiA islands in *E. albertii* strains.** Schematic representation of representative *E. albertii* LptG-YjiA islands. LptG and YjiA boundaries are represented with coloured outlines. Anti-phage systems predicted by PADLOC are coloured as indicated in the legend. Integrases, transposases, toxin-antitoxin systems (TA), predicted T6SS-dependent effectors are also coloured according to the legend.

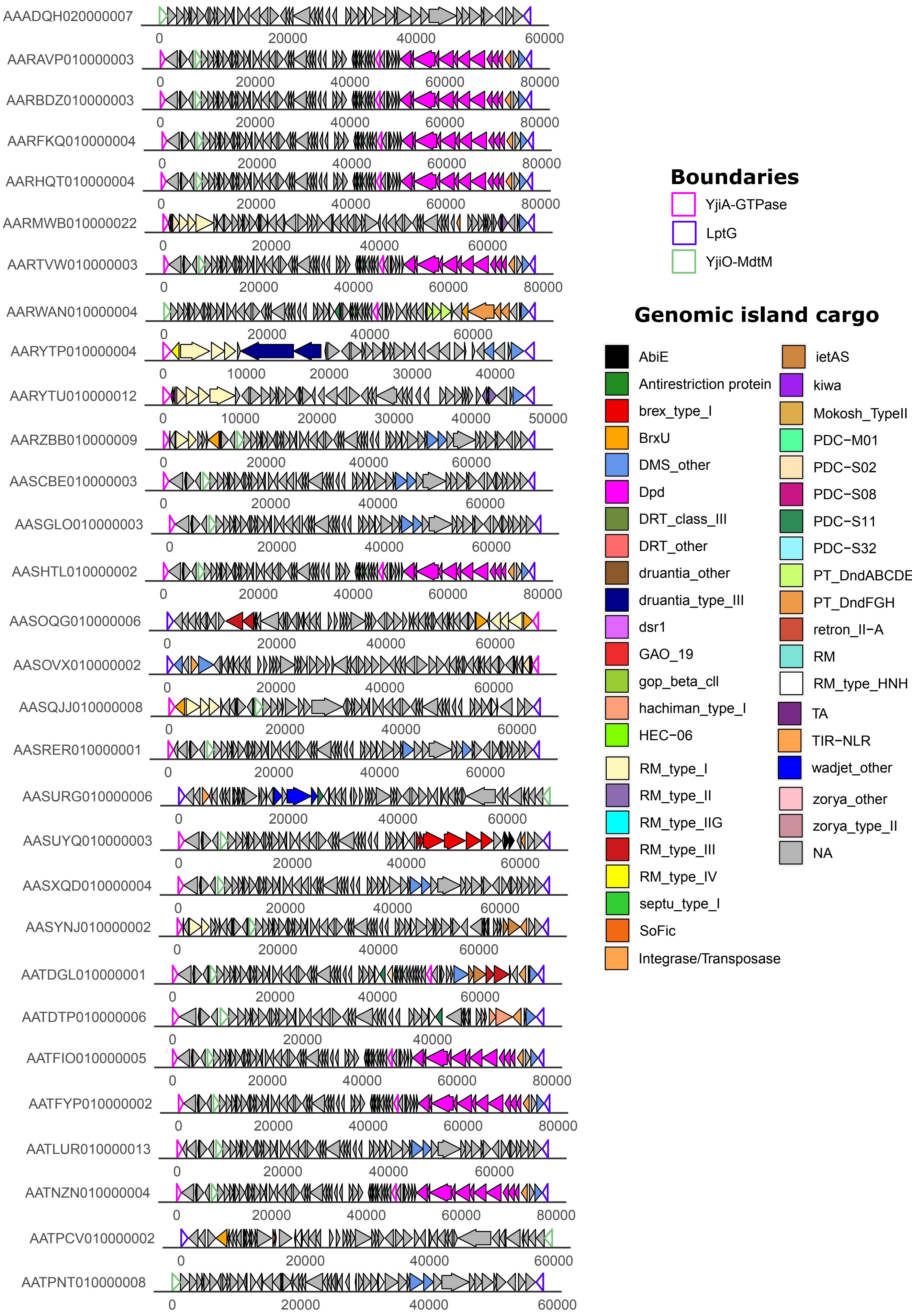

AAADQH020000007
AARAVP010000003
AARBDZ010000003
AARFKQ010000004
AARHQT010000004
AARMWB010000022
AARTVW010000003
AARWAN010000004
AARYTP010000004
AARYTU010000012
AARZBB010000009
AASCBE010000003
AASGLO010000003
AASHTL010000002
AASOQG010000006
AASOVX010000002
AASQJJ010000008
AASRER010000001
AASURG010000006
AASUYQ010000003
AASXQD010000004
AASYNJ010000002
AATDGL010000001
AATDTP010000006
AATFIO010000005
AATFYP010000002
AATLUR010000013
AATNZN010000004
AATPCV010000002
AATPNT010000008
0
10000
20000
30000
40000
50000
60000
80000
Boundaries
YjiA-GTPase
LptG
YjiO-MdtM
Genomic island cargo
AbiE
Antirestriction protein
brex_type_I
BrxU
DMS_other
Dpd
DRT_class_III
DRT_other
druantia_other
druantia_type_III
dsr1
GAO_19
gop_beta_cll
hachiman_type_I
HEC-06
RM_type_I
RM_type_II
RM_type_IIG
RM_type_III
RM_type_IV
septu_type_I
SoFic
Integrase/Transposase
ietAS
kiwa
Mokosh_TypeII
PDC-M01
PDC-S02
PDC-S08
PDC-S11
PDC-S32
PT_DndABCDE
PT_DndFGH
retron_II-A
RM
RM_type_HNH
TA
TIR-NLR
wadjet_other
zorya_other
zorya_type_II
NA

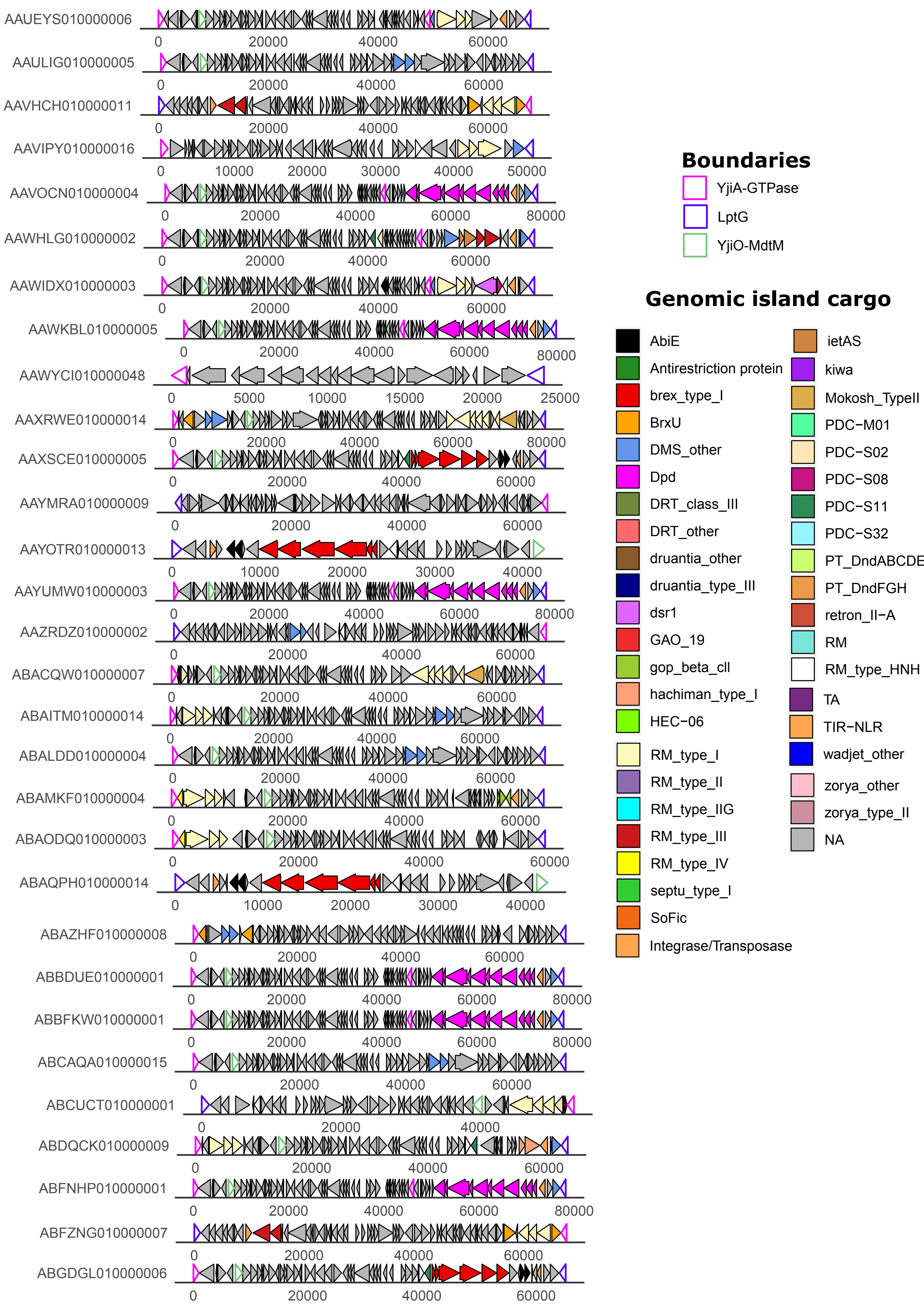
AAUEYS010000006
AAULIG010000005
AAVHCH010000011
AAVIPY010000016
AAVOCN010000004
AAWHLG010000002
AAWIDX010000003
AAWKBL010000005
AAWYCI010000048
AAXRWE010000014
AAXSCE010000005
AAYMRA010000009
AAYOTR010000013
AAYUMW010000003
AAZRDZ010000002
ABACQW010000007
ABAITM010000014
ABALDD010000004
ABAMKF010000004
ABAODQ010000003
ABAQPH010000014
ABAZHF010000008
ABBDUE010000001
ABBFKW010000001
ABCAQA010000015
ABCUCT010000001
ABDQCK010000009
ABFNHP010000001
ABFZNG010000007
ABGDGL010000006
Boundaries
YjiA-GTPase
LptG
YjiO-MdtM
Genomic island cargo
AbiE
Antirestriction protein
brex_type_I
BrxU
DMS_other
Dpd
DRT_class_III
DRT_other
druantia_other
druantia_type_III
dsr1
GAO_19
gop_beta_cll
hachiman_type_I
HEC-06
RM_type_I
RM_type_II
RM_type_IIG
RM_type_III
RM_type_IV
septu_type_I
SoFic
Integrase/Transposase
ietAS
kiwa
Mokosh_TypeII
PDC-M01
PDC-S02
PDC-S08
PDC-S11
PDC-S32
PT_DndABCDE
PT_DndFGH
retron_II-A
RM
RM_type_HNH
TA
TIR-NLR
wadjet_other
zorya_other
zorya_type_II
NA

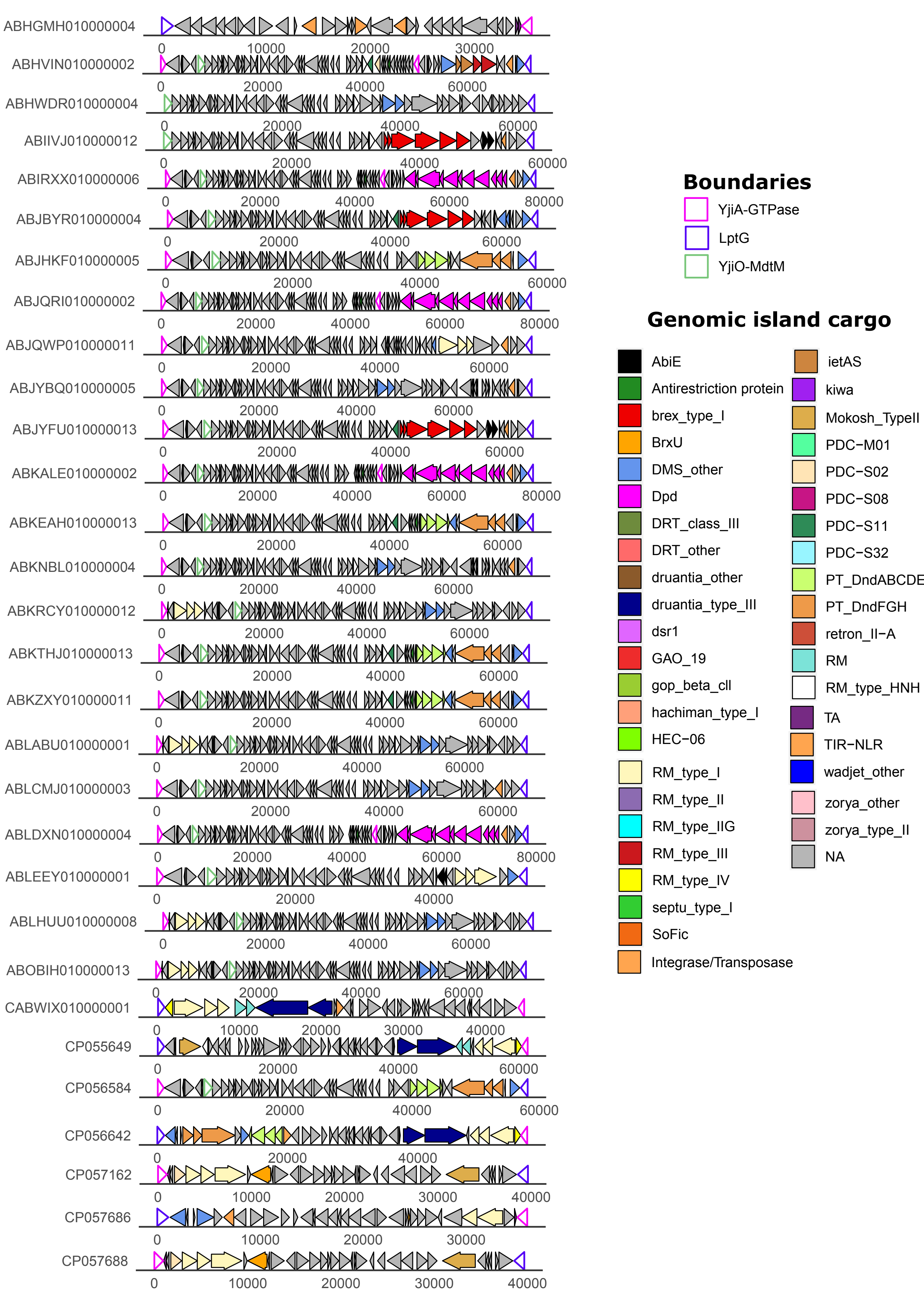

ABHGMH010000004
ABHVIN010000002
ABHWDR010000004
ABIIVJ010000012
ABIRXX010000006
ABJBYR010000004
ABJHKF010000005
ABJQRI010000002
ABJQWP010000011
ABJYBQ010000005
ABJYFU010000013
ABKALE010000002
ABKEAH010000013
ABKNBL010000004
ABKRCY010000012
ABKTHJ010000013
ABKZXY010000011
ABLABU010000001
ABLCMJ010000003
ABLDXN010000004
ABLEEY010000001
ABLHUU010000008
ABOBIH010000013
CABWIX010000001
CP055649
CP056584
CP056642
CP057162
CP057686
CP057688
Boundaries
YjiA-GTPase
LptG
YjiO-MdtM
Genomic island cargo
AbiE
Antirestriction protein
brex_type_I
BrxU
DMS_other
Dpd
DRT_class_III
DRT_other
druantia_other
druantia_type_III
dsr1
GAO_19
gop_beta_cll
hachiman_type_I
HEC-06
RM_type_I
RM_type_II
RM_type_IIG
RM_type_III
RM_type_IV
septu_type_I
SoFic
Integrase/Transposase
ietAS
kiwa
Mokosh_TypeII
PDC-M01
PDC-S02
PDC-S08
PDC-S11
PDC-S32
PT_DndABCDE
PT_DndFGH
retron_II-A
RM
RM_type_HNH
TA
TIR-NLR
wadjet_other
zorya_other
zorya_type_II
NA

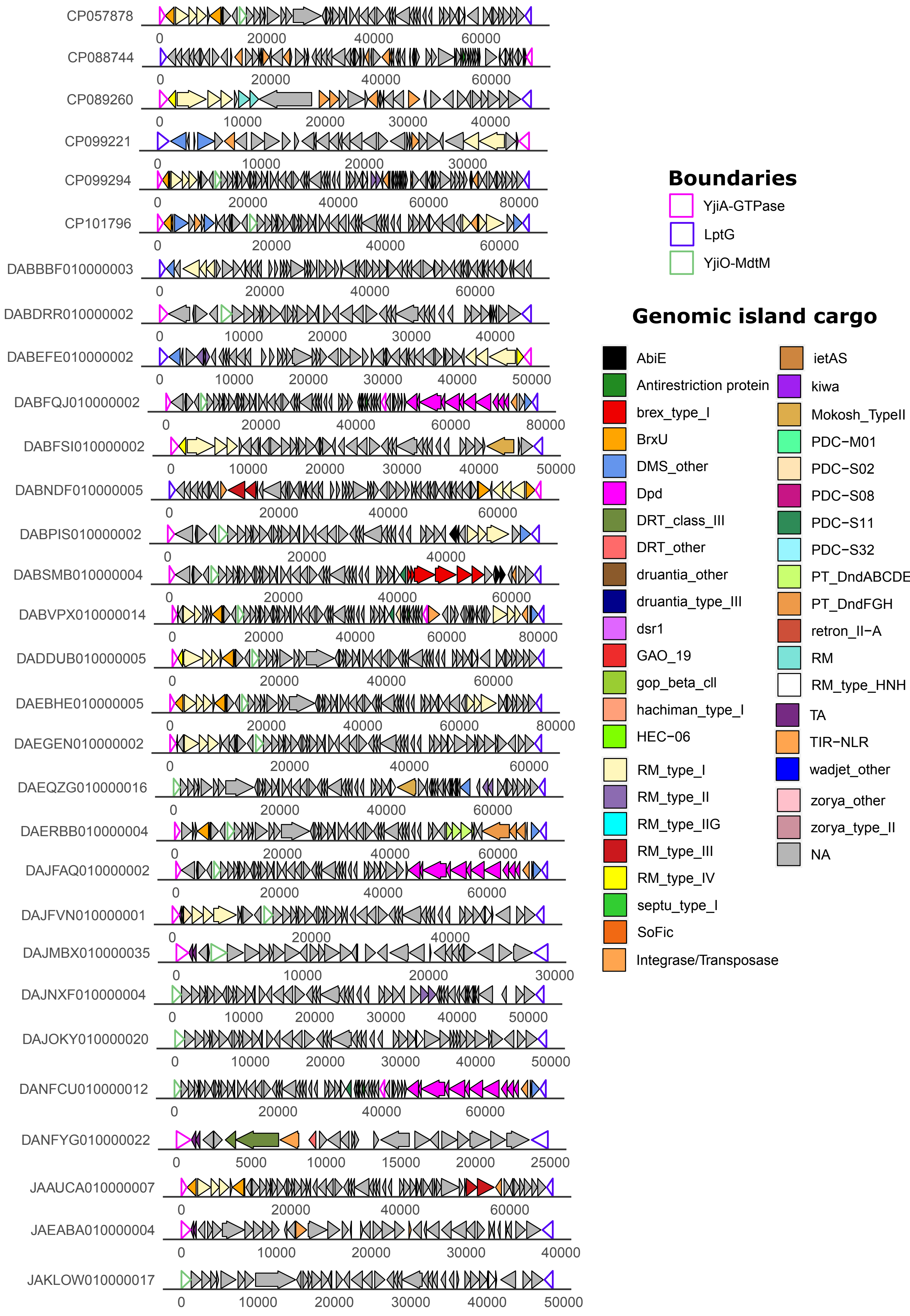

CP057878
CP088744
CP089260
CP099221
CP099294
CP101796
DABBBF010000003
DABDRR010000002
DABEFE010000002
DABFQJ010000002
DABFSI010000002
DABNDF010000005
DABPIS010000002
DABSMB010000004
DABVPX010000014
DADDUB010000005
DAEBHE010000005
DAEGEN010000002
DAEQZG010000016
DAERBB010000004
DAJFAQ010000002
DAJFVN010000001
DAJMBX010000035
DAJNXF010000004
DAJOKY010000020
DANFCU010000012
DANFYG010000022
JAAUCA010000007
JAEABA010000004
JAKLOW010000017
Boundaries
YjiA-GTPase
LptG
YjiO-MdtM
Genomic island cargo
AbiE
Antirestriction protein
brex_type_I
BrxU
DMS_other
Dpd
DRT_class_III
DRT_other
druantia_other
druantia_type_III
dsr1
GAO_19
gop_beta_cll
hachiman_type_I
HEC-06
RM_type_I
RM_type_II
RM_type_IIG
RM_type_III
RM_type_IV
septu_type_I
SoFic
Integrase/Transposase
ietAS
kiwa
Mokosh_TypeII
PDC-M01
PDC-S02
PDC-S08
PDC-S11
PDC-S32
PT_DndABCDE
PT_DndFGH
retron_II-A
RM
RM_type_HNH
TA
TIR-NLR
wadjet_other
zorya_other
zorya_type_II
NA

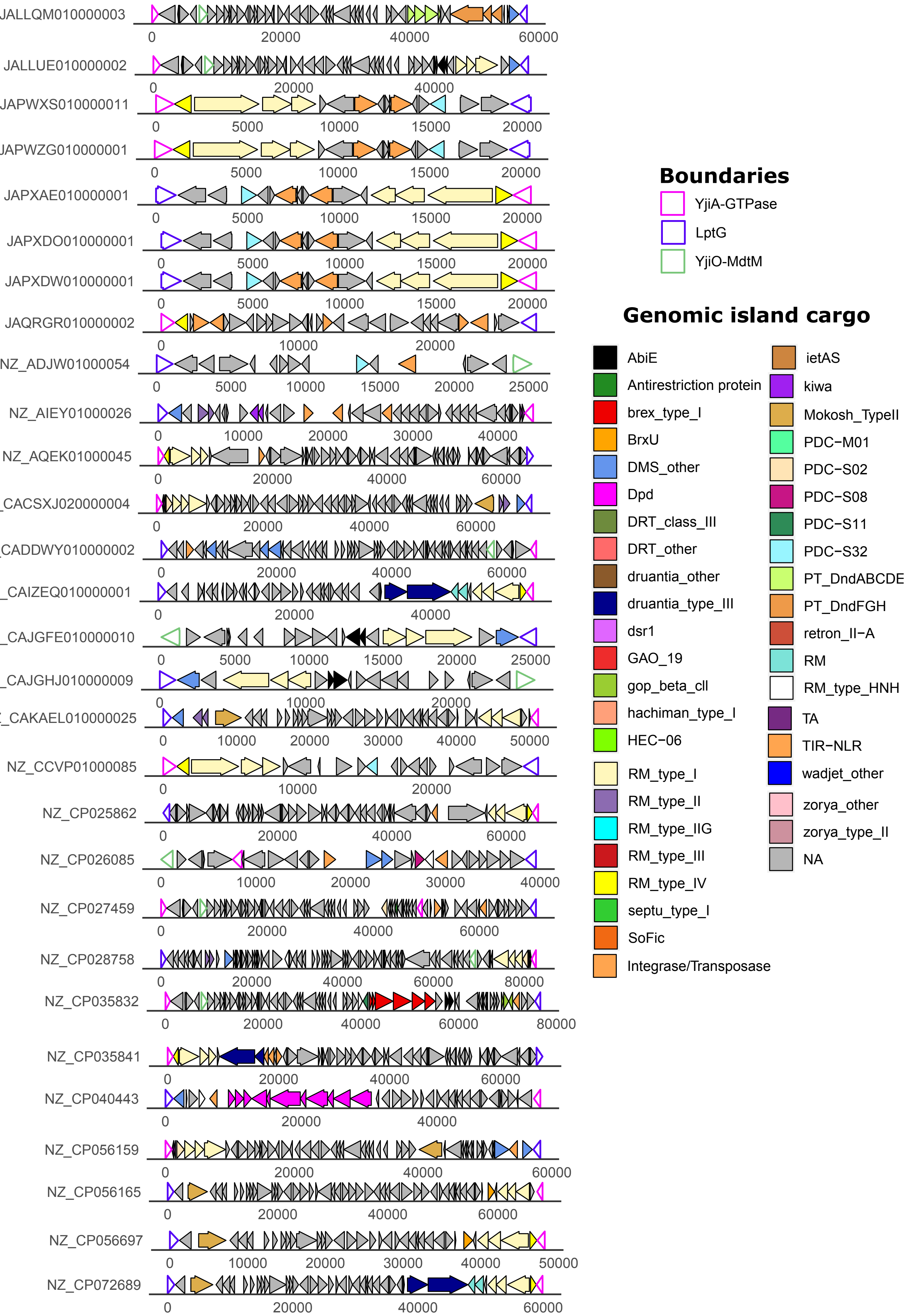

JALLQM010000003
JALLUE010000002
JAPWXS010000011
JAPWZG010000001
JAPXAE010000001
JAPXDO010000001
JAPXDW010000001
JAQRGR010000002
NZ_ADJW01000054
NZ_AIEY01000026
NZ_AQEK01000045
CACSXJ020000004
CADDWY010000002
CAIZEQ010000001
CAJGFE010000010
CAJGHJ010000009
CAKAEL010000025
NZ_CCVP01000085
NZ_CP025862
NZ_CP026085
NZ_CP027459
NZ_CP028758
NZ_CP035832
NZ_CP035841
NZ_CP040443
NZ_CP056159
NZ_CP056165
NZ_CP056697
NZ_CP072689
Boundaries
YjiA-GTPase
LptG
YjiO-MdtM
Genomic island cargo
AbiE
Antirestriction protein
brex_type_I
BrxU
DMS_other
Dpd
DRT_class_III
DRT_other
druantia_other
druantia_type_III
dsr1
GAO_19
gop_beta_cll
hachiman_type_I
HEC-06
RM_type_I
RM_type_II
RM_type_IIG
RM_type_III
RM_type_IV
septu_type_I
SoFic
Integrase/Transposase
ietAS
kiwa
Mokosh_TypeII
PDC-M01
PDC-S02
PDC-S08
PDC-S11
PDC-S32
PT_DndABCDE
PT_DndFGH
retron_II-A
RM
RM_type_HNH
TA
TIR-NLR
wadjet_other
zorya_other
zorya_type_II
NA

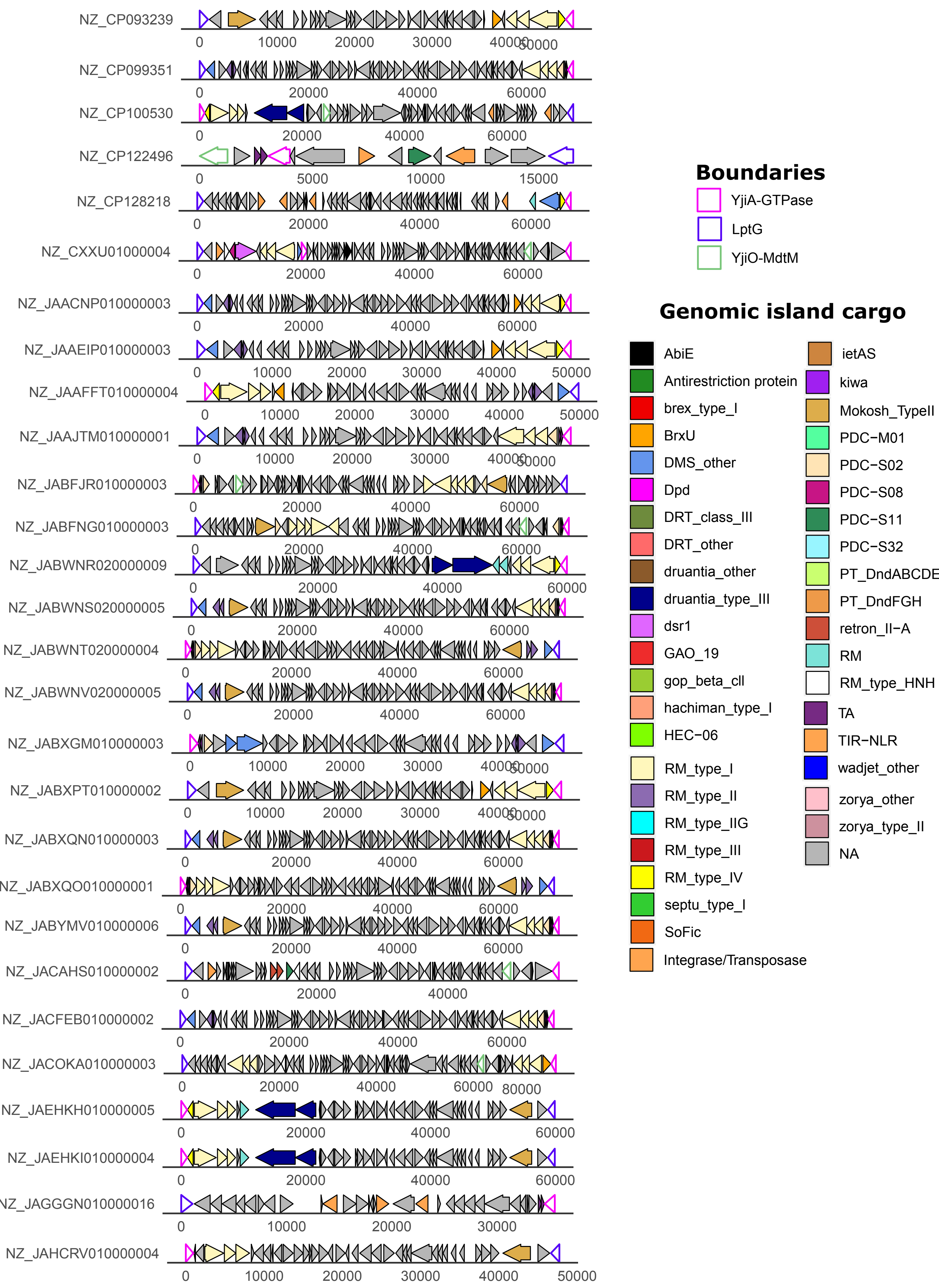

NZ_CP093239
NZ_CP099351
NZ_CP100530
NZ_CP122496
NZ_CP128218
NZ_CXXU01000004
NZ_JAACNP010000003
NZ_JAAEIP010000003
NZ_JAAFFT010000004
NZ_JAAJTM010000001
NZ_JABFJR010000003
NZ_JABFNG010000003
NZ_JABWNR020000009
NZ_JABWNS020000005
NZ_JABWNT020000004
NZ_JABWNV020000005
NZ_JABXGM010000003
NZ_JABXPT010000002
NZ_JABXQN010000003
NZ_JABXQO010000001
NZ_JABYMV010000006
NZ_JACAHS010000002
NZ_JACFEB010000002
NZ_JACOKA010000003
NZ_JAEHKH010000005
NZ_JAEHKI010000004
NZ_JAGGGN010000016
NZ_JAHCRV010000004
Boundaries
YjiA-GTPase
LptG
YjiO-MdtM
Genomic island cargo
AbiE
Antirestriction protein
brex_type_I
BrxU
DMS_other
Dpd
DRT_class_III
DRT_other
druantia_other
druantia_type_III
dsr1
GAO_19
gop_beta_cll
hachiman_type_I
HEC-06
RM_type_I
RM_type_II
RM_type_IIG
RM_type_III
RM_type_IV
septu_type_I
SoFic
Integrase/Transposase
ietAS
kiwa
Mokosh_TypeII
PDC-M01
PDC-S02
PDC-S08
PDC-S11
PDC-S32
PT_DndABCDE
PT_DndFGH
retron_II-A
RM
RM_type_HNH
TA
TIR-NLR
wadjet_other
zorya_other
zorya_type_II
NA

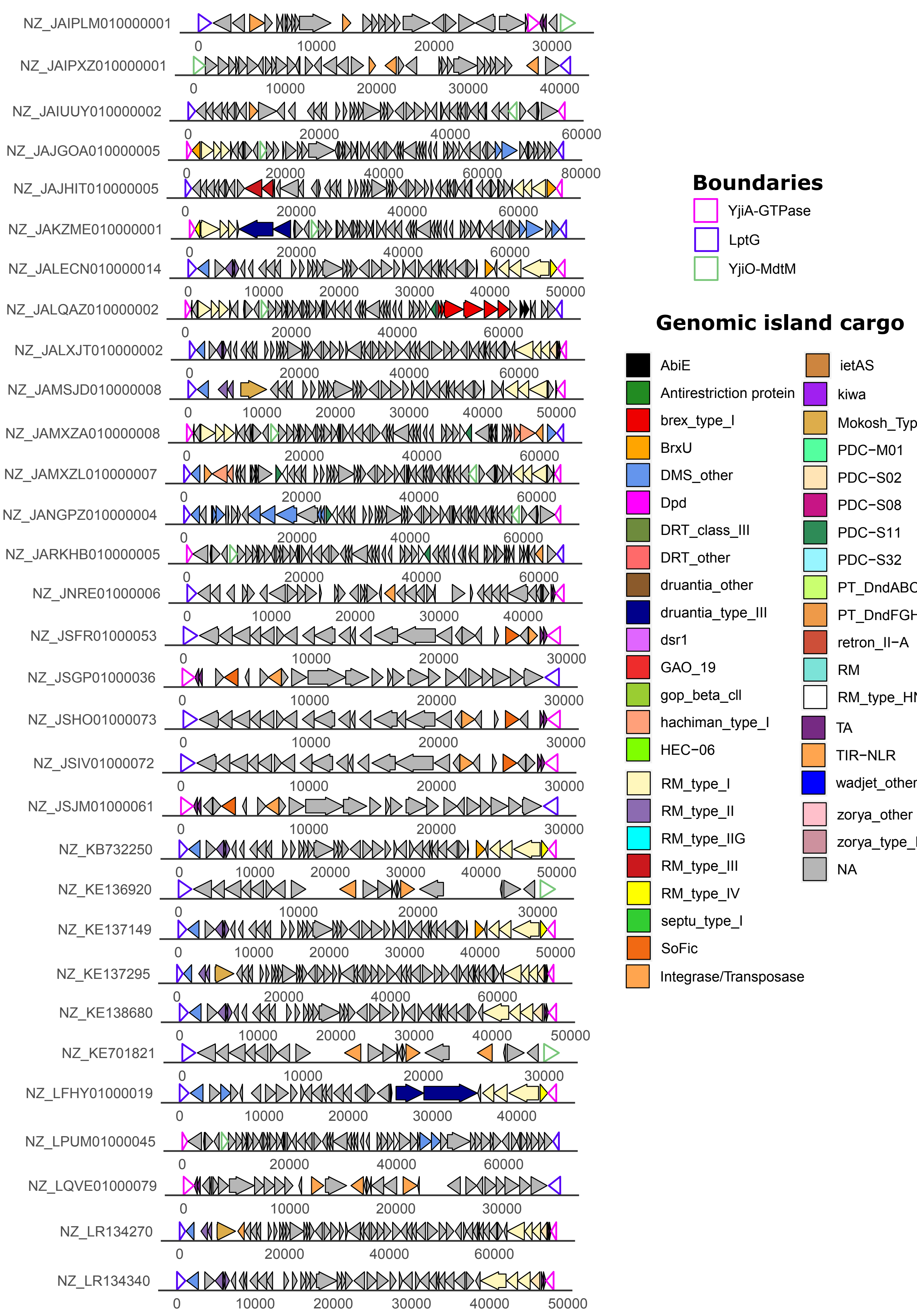

NZ_JAIPLM010000001
NZ_JAIPXZ010000001
NZ_JAIUUY010000002
NZ_JAJGOA010000005
NZ_JAJHIT010000005
NZ_JAKZME010000001
NZ_JALECN010000014
NZ_JALQAZ010000002
NZ_JALXJT010000002
NZ_JAMSJD010000008
NZ_JAMXZA010000008
NZ_JAMXZL010000007
NZ_JANGPZ010000004
NZ_JARKHB010000005
NZ_JNRE01000006
NZ_JSFR01000053
NZ_JSGP01000036
NZ_JSHO01000073
NZ_JSIV01000072
NZ_JSJM01000061
NZ_KB732250
NZ_KE136920
NZ_KE137149
NZ_KE137295
NZ_KE138680
NZ_KE701821
NZ_LFHY01000019
NZ_LPUM01000045
NZ_LQVE01000079
NZ_LR134270
NZ_LR134340
Boundaries
YjiA-GTPase
LptG
YjiO-MdtM
Genomic island cargo
AbiE
Antirestriction protein
brex_type_I
BrxU
DMS_other
Dpd
DRT_class_III
DRT_other
druantia_other
druantia_type_III
dsr1
GAO_19
gop_beta_cll
hachiman_type_I
HEC-06
RM_type_I
RM_type_II
RM_type_IIG
RM_type_III
RM_type_IV
septu_type_I
SoFic
Integrase/Transposase
ietAS
kiwa
PDC-M01
PDC-S02
PDC-S08
PDC-S11
PDC-S32
retron_II-A
RM
TA
TIR-NLR
wadjet_other
zorya_other
NA

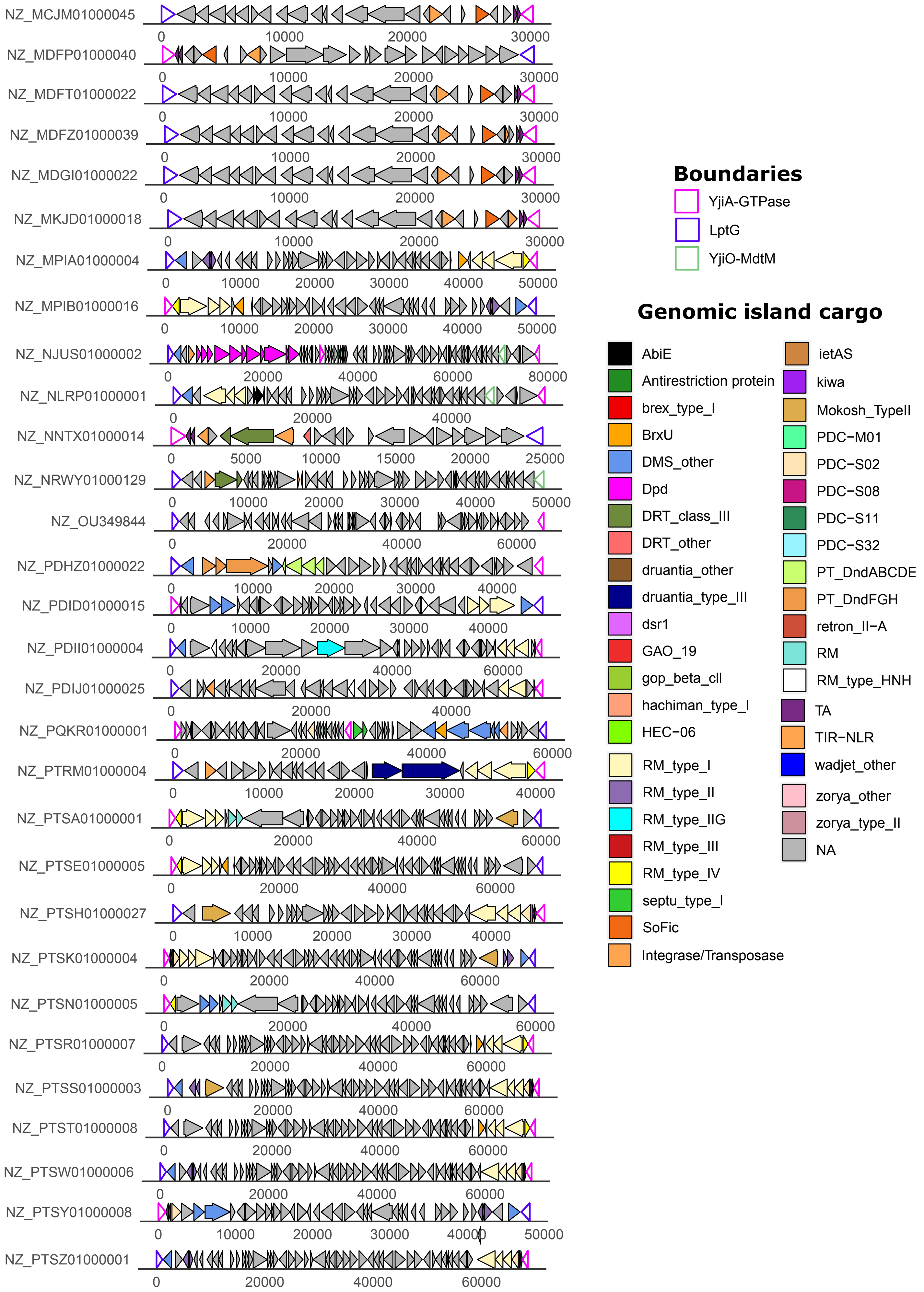

NZ_MCJM01000045
NZ_MDFP01000040
NZ_MDFT01000022
NZ_MDFZ01000039
NZ_MDGI01000022
NZ_MKJD01000018
NZ_MPIA01000004
NZ_MPIB01000016
NZ_NJUS01000002
NZ_NLRP01000001
NZ_NNTX01000014
NZ_NRWY01000129
NZ_OU349844
NZ_PDHZ01000022
NZ_PDID01000015
NZ_PDII01000004
NZ_PDIJ01000025
NZ_PQKR01000001
NZ_PTRM01000004
NZ_PTSA01000001
NZ_PTSE01000005
NZ_PTSH01000027
NZ_PTSK01000004
NZ_PTSN01000005
NZ_PTSR01000007
NZ_PTSS01000003
NZ_PTST01000008
NZ_PTSW01000006
NZ_PTSY01000008
NZ_PTSZ01000001
Boundaries
YjiA-GTPase
LptG
YjiO-MdtM
Genomic island cargo
AbiE
Antirestriction protein
brex_type_I
BrxU
DMS_other
Dpd
DRT_class_III
DRT_other
druantia_other
druantia_type_III
dsr1
GAO_19
gop_beta_cll
hachiman_type_I
HEC-06
RM_type_I
RM_type_II
RM_type_IIG
RM_type_III
RM_type_IV
septu_type_I
SoFic
Integrase/Transposase
ietAS
kiwa
Mokosh_TypeII
PDC-M01
PDC-S02
PDC-S08
PDC-S11
PDC-S32
PT_DndABCDE
PT_DndFGH
retron_II-A
RM
RM_type_HNH
TA
TIR-NLR
wadjet_other
zorya_other
zorya_type_II
NA

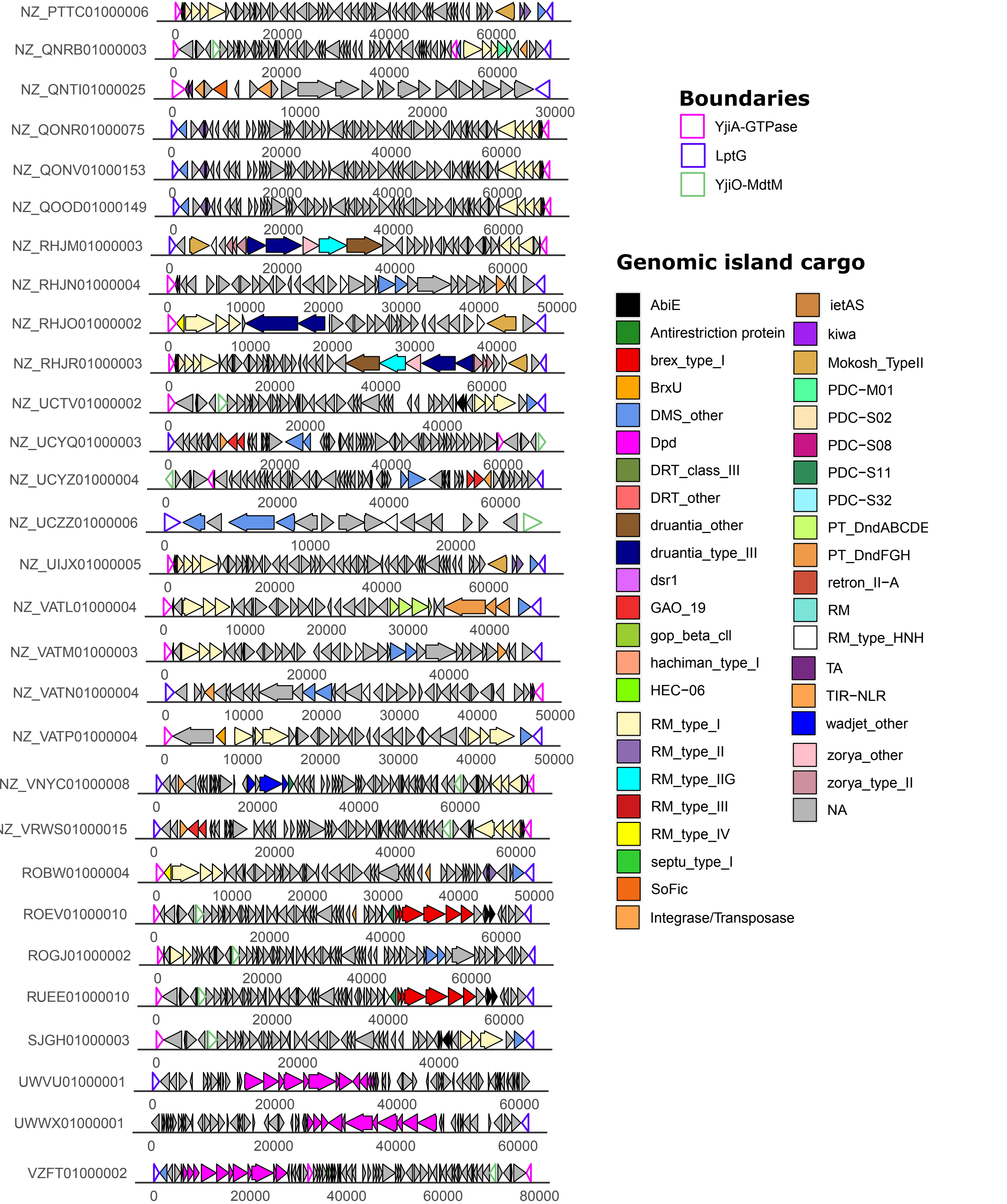


**Figure S5. *E. coli* predominantly carry LptG-YjiO-YjiA with a double insertion point between LptG-YjiO and YjiO-YjiA**. Representative E. coli LptG-YjiO-YjiA were chosen by filtering closely related nucleotide sequences (90% sequence identity threshold) with fastANI. LptG, YjiO and YjiA boundaries are depicted with coloured outlines. Anti-phage systems predicted by PADLOC, integrases, transposases, toxin-antitoxin systems (TA), predicted T6SS-dependent effectors are coloured according to the legend in the figure.

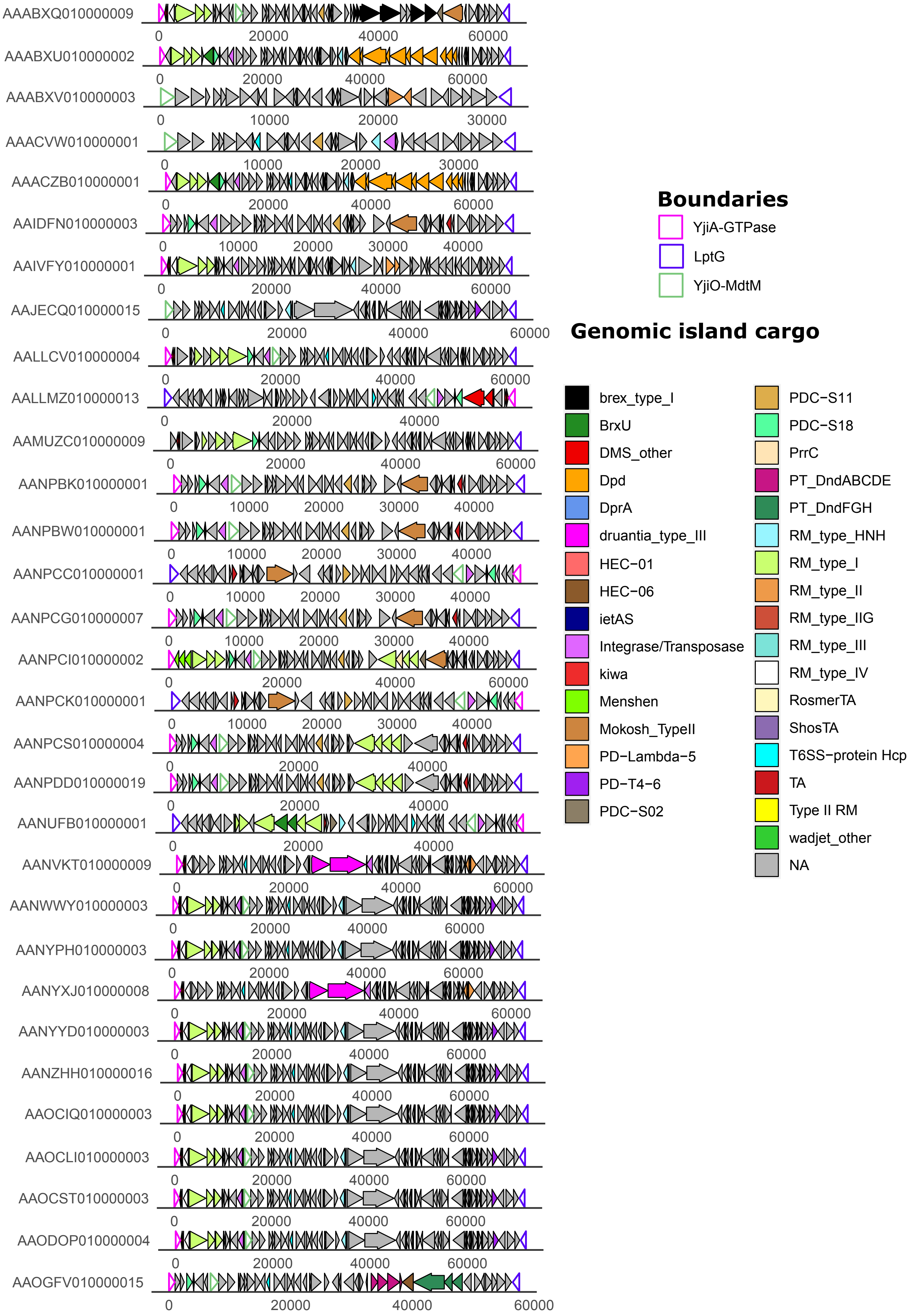
AAABXQ010000009
AAABXU010000002
AAABXV010000003
AAACVW010000001
AAACZB010000001
AAIDFN010000003
AAIVFY010000001
AAJECQ010000015
AALLCV010000004
AALLMZ010000013
AAMUZC010000009
AANPBK010000001
AANPBW010000001
AANPCC010000001
AANPCG010000007
AANPCI010000002
AANPCK010000001
AANPCS010000004
AANPDD010000019
AANUFB010000001
AANVKT010000009
AANWWY010000003
AANYPH010000003
AANYXJ010000008
AANYYD010000003
AANZHH010000016
AAOCIQ010000003
AAOCLI010000003
AAOCST010000003
AAODOP010000004
AAOGFV010000015
Boundaries
YjiA-GTPase
LptG
YjiO-MdtM
Genomic island cargo
brex_type_I
BrxU
DMS_other
Dpd
DprA
druantia_type_III
HEC-01
HEC-06
ietAS
Integrase/Transposase
kiwa
Menshen
Mokosh_TypeII
PD-Lambda-5
PD-T4-6
PDC-S02
PDC-S11
PDC-S18
PrrC
PT_DndABCDE
PT_DndFGH
RM_type_HNH
RM_type_I
RM_type_II
RM_type_IIG
RM_type_III
RM_type_IV
RosmerTA
ShosTA
T6SS-protein Hcp
TA
Type II RM
wadjet_other
NA

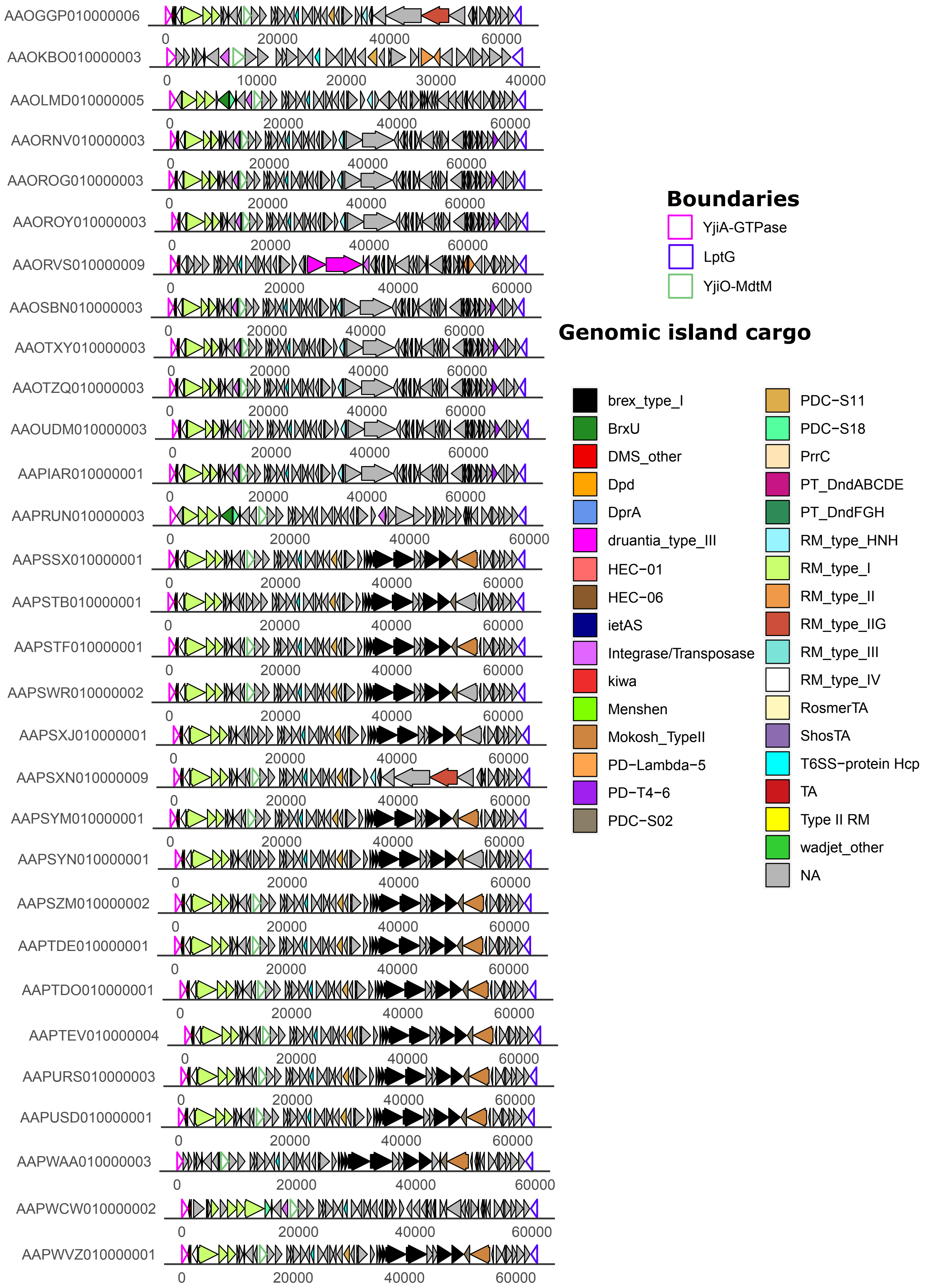
AAOGGP010000006
AAOKBO010000003
AAOLMD010000005
AAORNV010000003
AAOROG010000003
AAOROY010000003
AAORVS010000009
AAOSBN010000003
AAOTXY010000003
AAOTZQ010000003
AAOUDM010000003
AAPIAR010000001
AAPRUN010000003
AAPSSX010000001
AAPSTB010000001
AAPSTF010000001
AAPSWR010000002
AAPSXJ010000001
AAPSXN010000009
AAPSYM010000001
AAPSYN010000001
AAPSZM010000002
AAPTDE010000001
AAPTDO010000001
AAPTEV010000004
AAPURS010000003
AAPUSD010000001
AAPWAA010000003
AAPWCW010000002
AAPWVZ010000001
Boundaries
YjiA-GTPase
LptG
YjiO-MdtM
Genomic island cargo
brex_type_I
BrxU
DMS_other
Dpd
DprA
druantia_type_III
HEC-01
HEC-06
ietAS
Integrase/Transposase
kiwa
Menshen
Mokosh_TypeII
PD-Lambda-5
PD-T4-6
PDC-S02
PDC-S11
PDC-S18
PrrC
PT_DndABCDE
PT_DndFGH
RM_type_HNH
RM_type_I
RM_type_II
RM_type_IIG
RM_type_III
RM_type_IV
RosmerTA
ShosTA
T6SS-protein Hcp
TA
Type II RM
wadjet_other
NA

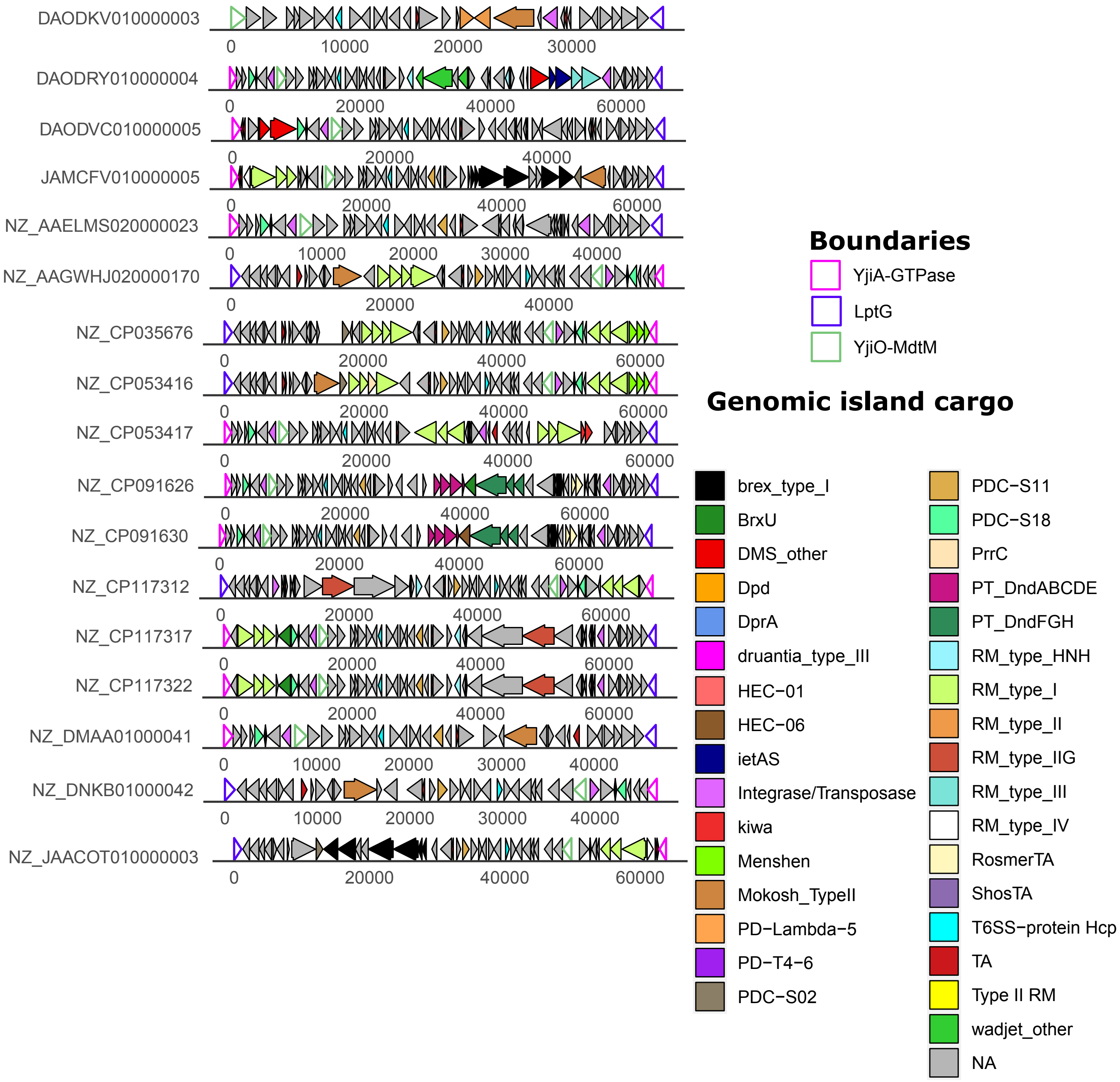


**Figure S6. LptG-YjiO-YjiA are abundant in *Salmonella spp.*** Schematic representation of LptG-YjiO-YjiA islands in *Salmonella* spp. Representative hotspots were chosen by filtering closely related nucleotide sequences (90% sequence identity threshold) with fastANI. LptG, YjiO and YjiA boundaries are represented with coloured outlines. Anti-phage systems predicted by PADLOC are coloured as indicated in the legend. Integrases, transposases, toxin-antitoxin systems (TA), predicted T6SS-dependent effectors are also coloured according to the legend.

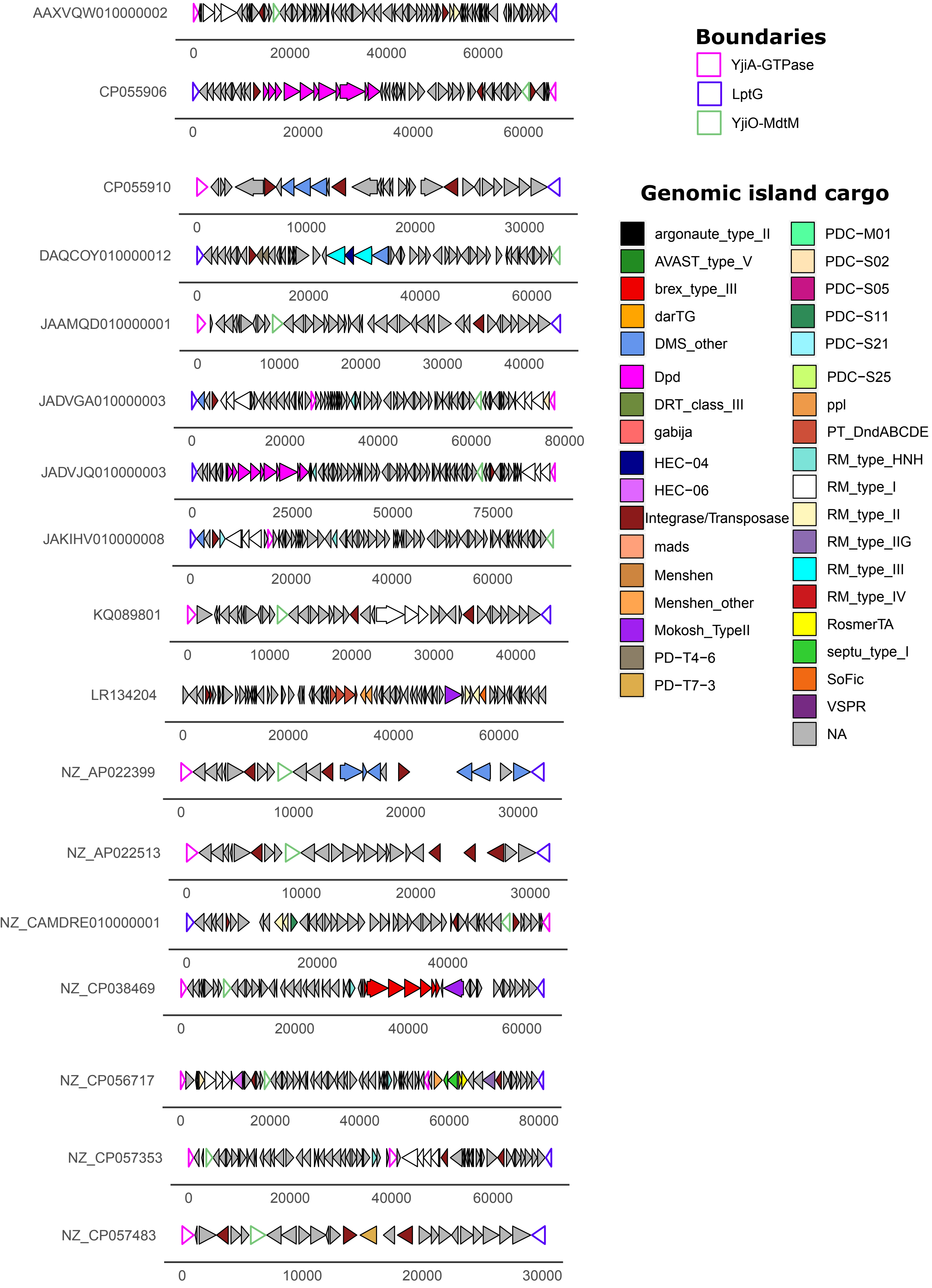

AAXVQW010000002
CP055906
CP055910
DAQCOY010000012
JAAMQD010000001
JADVGA010000003
JADVJQ010000003
JAKIHV010000008
KQ089801
LR134204
NZ_AP022399
NZ_AP022513
NZ_CAMDRE010000001
NZ_CP038469
NZ_CP056717
NZ_CP057353
NZ_CP057483
Boundaries
YjiA-GTPase
LptG
YjiO-MdtM
Genomic island cargo
argonaute_type_II
AVAST_type_V
brex_type_III
darTG
DMS_other
Dpd
DRT_class_III
gabija
HEC−04
HEC−06
Integrase/Transposase
mads
Menshen
Menshen_other
Mokosh_TypeII
PD−T4−6
PD−T7−3
PDC−M01
PDC−S02
PDC−S05
PDC−S11
PDC−S21
PDC−S25
ppl
PT_DndABCDE
RM_type_HNH
RM_type_I
RM_type_II
RM_type_IIG
RM_type_III
RM_type_IV
RosmerTA
septu_type_I
SoFic
VSPR
NA

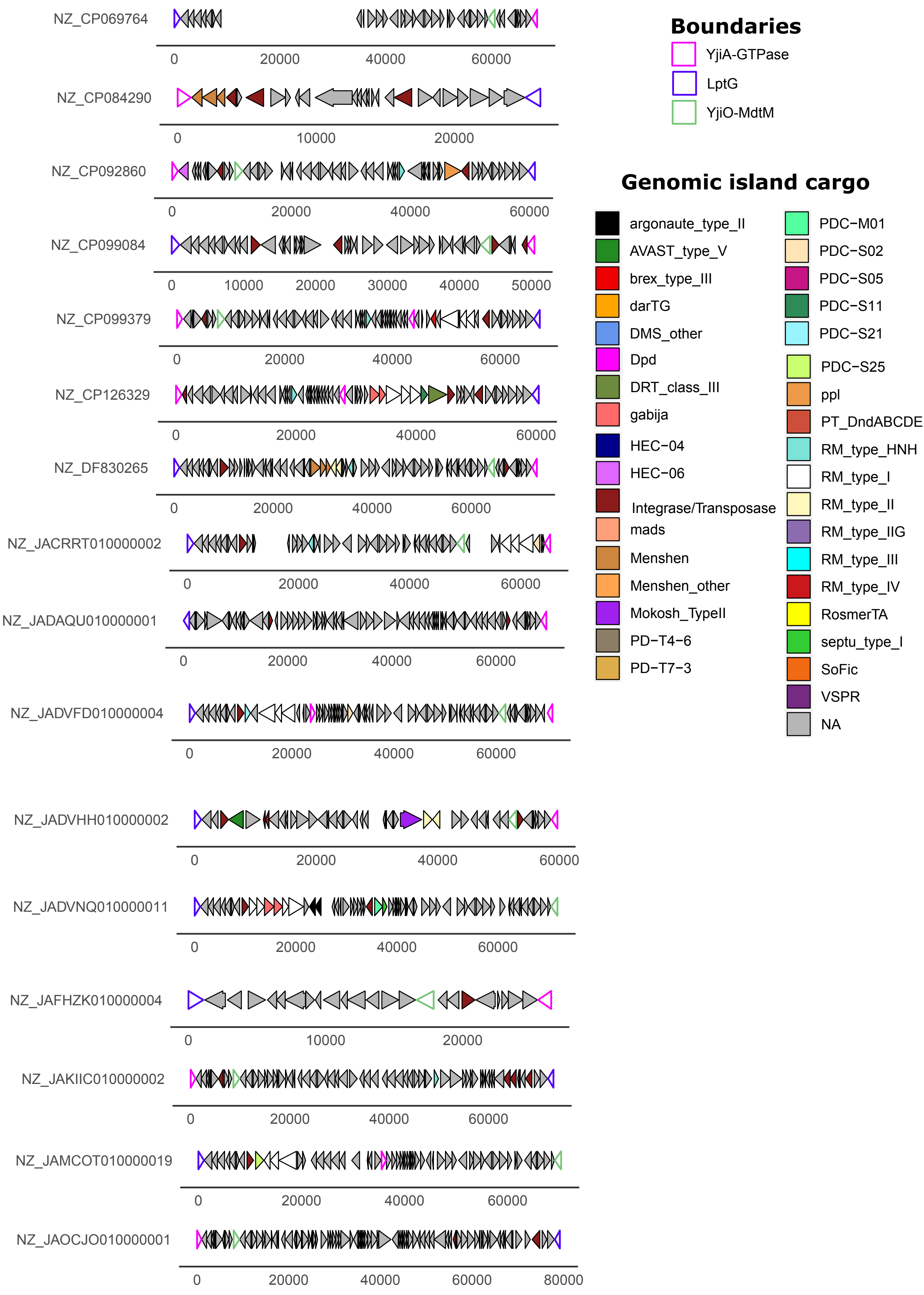
NZ_CP069764
0 20000 40000 60000
NZ_CP084290
0 10000 20000
NZ_CP092860
0 20000 40000 60000
NZ_CP099084
0 10000 20000 30000 40000 50000
NZ_CP099379
0 20000 40000 60000
NZ_CP126329
0 20000 40000 60000
NZ_DF830265
0 20000 40000 60000
NZ_JACRRT010000002
0 20000 40000 60000
NZ_JADAQU010000001
0 20000 40000 60000
NZ_JADVFD010000004
0 20000 40000 60000
NZ_JADVHH010000002
0 20000 40000 60000
NZ_JADVNQ010000011
0 20000 40000 60000
NZ_JAFHZK010000004
0 10000 20000
NZ_JAKIIC010000002
0 20000 40000 60000
NZ_JAMCOT010000019
0 20000 40000 60000
NZ_JAOCJO010000001
0 20000 40000 60000 80000
Boundaries
YjiA-GTPase
LptG
YjiO-MdtM
Genomic island cargo
argonaute_type_II
AVAST_type_V
brex_type_III
darTG
DMS_other
Dpd
DRT_class_III
gabija
HEC−04
HEC−06
Integrase/Transposase
mads
Menshen
Menshen_other
Mokosh_TypeII
PD−T4−6
PD−T7−3
PDC−M01
PDC−S02
PDC−S05
PDC−S11
PDC−S21
PDC−S25
ppl
PT_DndABCDE
RM_type_HNH
RM_type_I
RM_type_II
RM_type_IIG
RM_type_III
RM_type_IV
RosmerTA
septu_type_I
SoFic
VSPR
NA

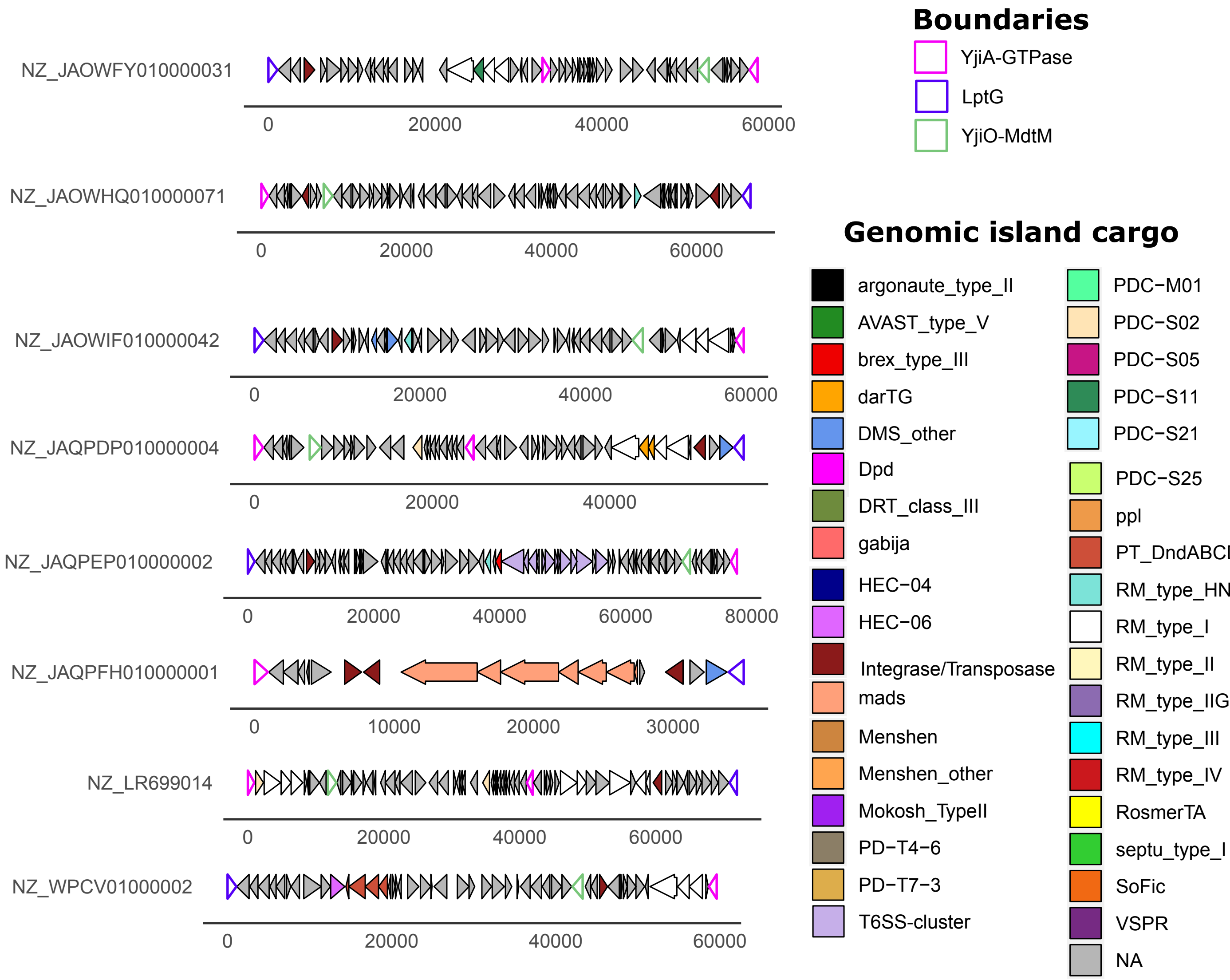


**Figure S7.** ***Citrobacter spp.*** **carry LptG-YjiO-YjiA islands.** Schematic representation of LptG-YjiO-YjiA islands in *Citrobacter* spp. Representative hotspots were chosen by filtering identical nucleotide sequences (90% sequence identity threshold) with fastANI. LptG, YjiO and YjiA boundaries are represented with coloured outlines. Anti-phage systems predicted by PADLOC, integrases, transposases, toxin-antitoxin systems (TA), predicted T6SS-dependent effectors are coloured according to the legend.

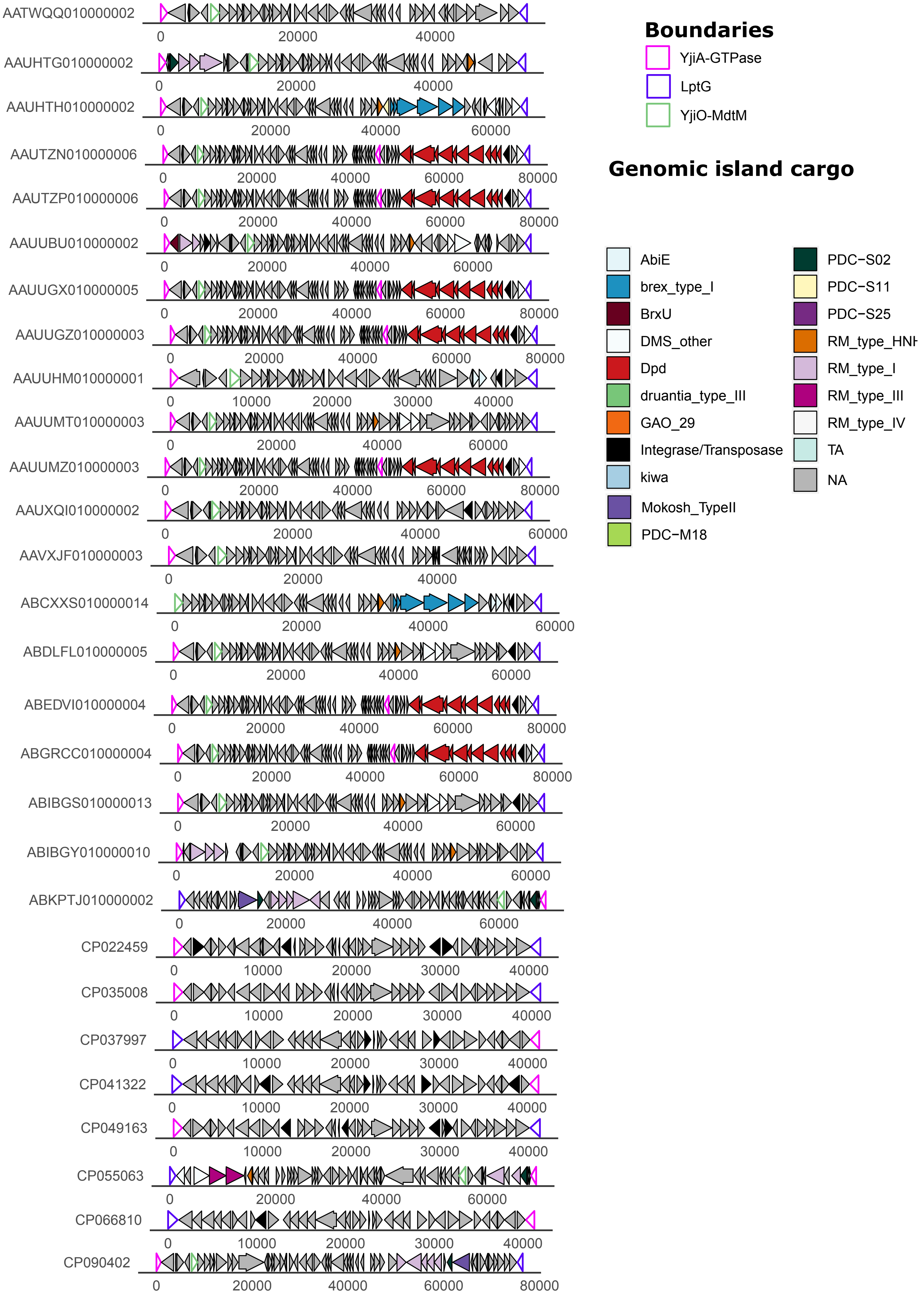
Boundaries
YjiA-GTPase
LptG
YjiO-MdtM
Genomic island cargo
AbiE
brex_type_I
BrxU
DMS_other
Dpd
druantia_type_III
GAO_29
Integrase/Transposase
kiwa
Mokosh_TypeII
PDC−M18
PDC−S02
PDC−S11
PDC−S25
RM_type_HNH
RM_type_I
RM_type_III
RM_type_IV
TA
NA
AATWQQ010000002
AAUHTG010000002
AAUHTH010000002
AAUTZN010000006
AAUTZP010000006
AAUUBU010000002
AAUUGX010000005
AAUUGZ010000003
AAUUHM010000001
AAUUMT010000003
AAUUMZ010000003
AAUXQI010000002
AAVXJF010000003
ABCXXS010000014
ABDLFL010000005
ABEDVI010000004
ABGRCC010000004
ABIBGS010000013
ABIBGY010000010
ABKPTJ010000002
CP022459
CP035008
CP037997
CP041322
CP049163
CP055063
CP066810
CP090402

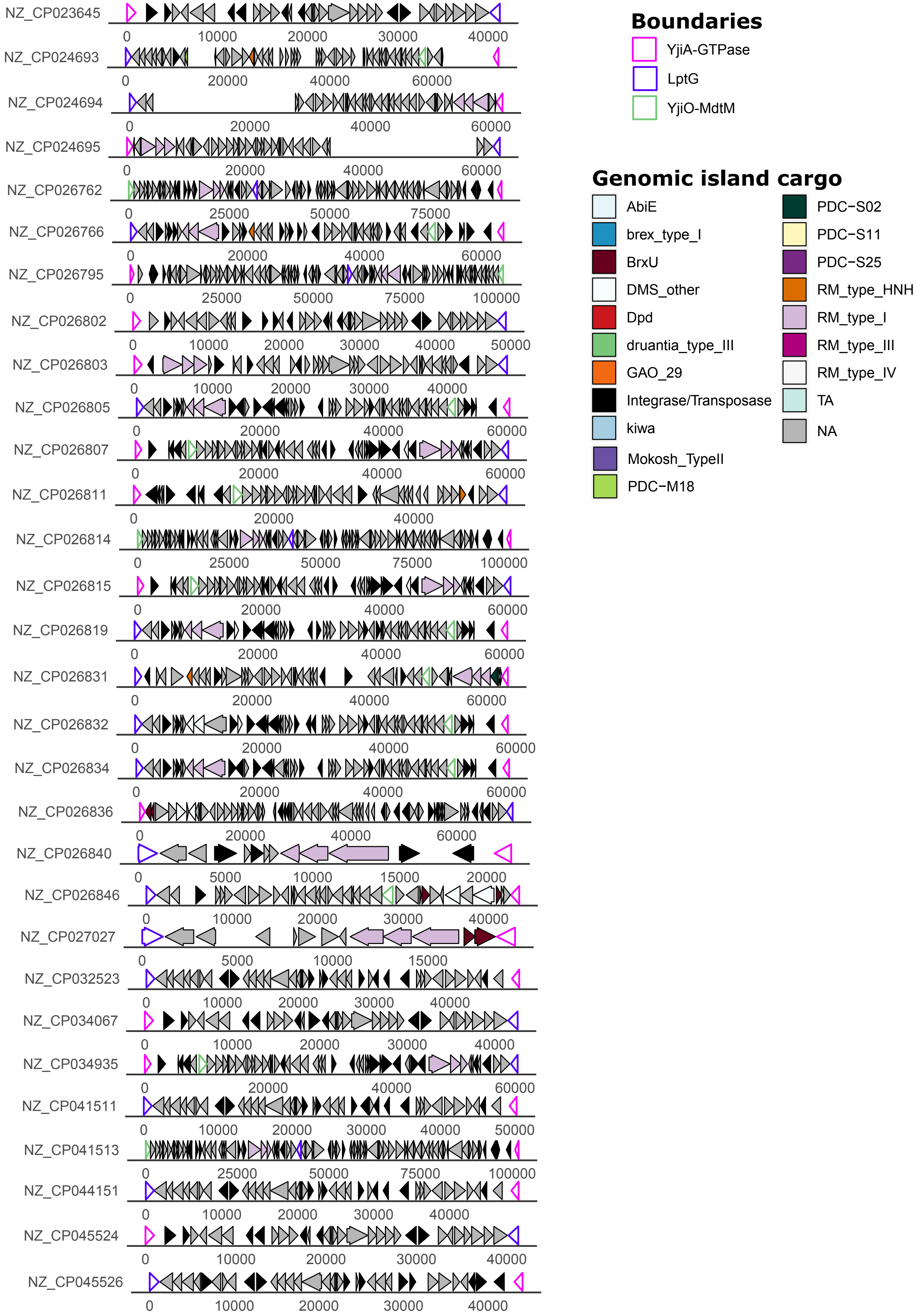
NZ_CP023645
NZ_CP024693
NZ_CP024694
NZ_CP024695
NZ_CP026762
NZ_CP026766
NZ_CP026795
NZ_CP026802
NZ_CP026803
NZ_CP026805
NZ_CP026807
NZ_CP026811
NZ_CP026814
NZ_CP026815
NZ_CP026819
NZ_CP026831
NZ_CP026832
NZ_CP026834
NZ_CP026836
NZ_CP026840
NZ_CP026846
NZ_CP027027
NZ_CP032523
NZ_CP034067
NZ_CP034935
NZ_CP041511
NZ_CP041513
NZ_CP044151
NZ_CP045524
NZ_CP045526
Boundaries
YjiA-GTPase
LptG
YjiO-MdtM
Genomic island cargo
AbiE
brex_type_I
BrxU
DMS_other
Dpd
druantia_type_III
GAO_29
Integrase/Transposase
kiwa
Mokosh_TypeII
PDC−M18
PDC−S02
PDC−S11
PDC−S25
RM_type_HNH
RM_type_I
RM_type_III
RM_type_IV
TA
NA

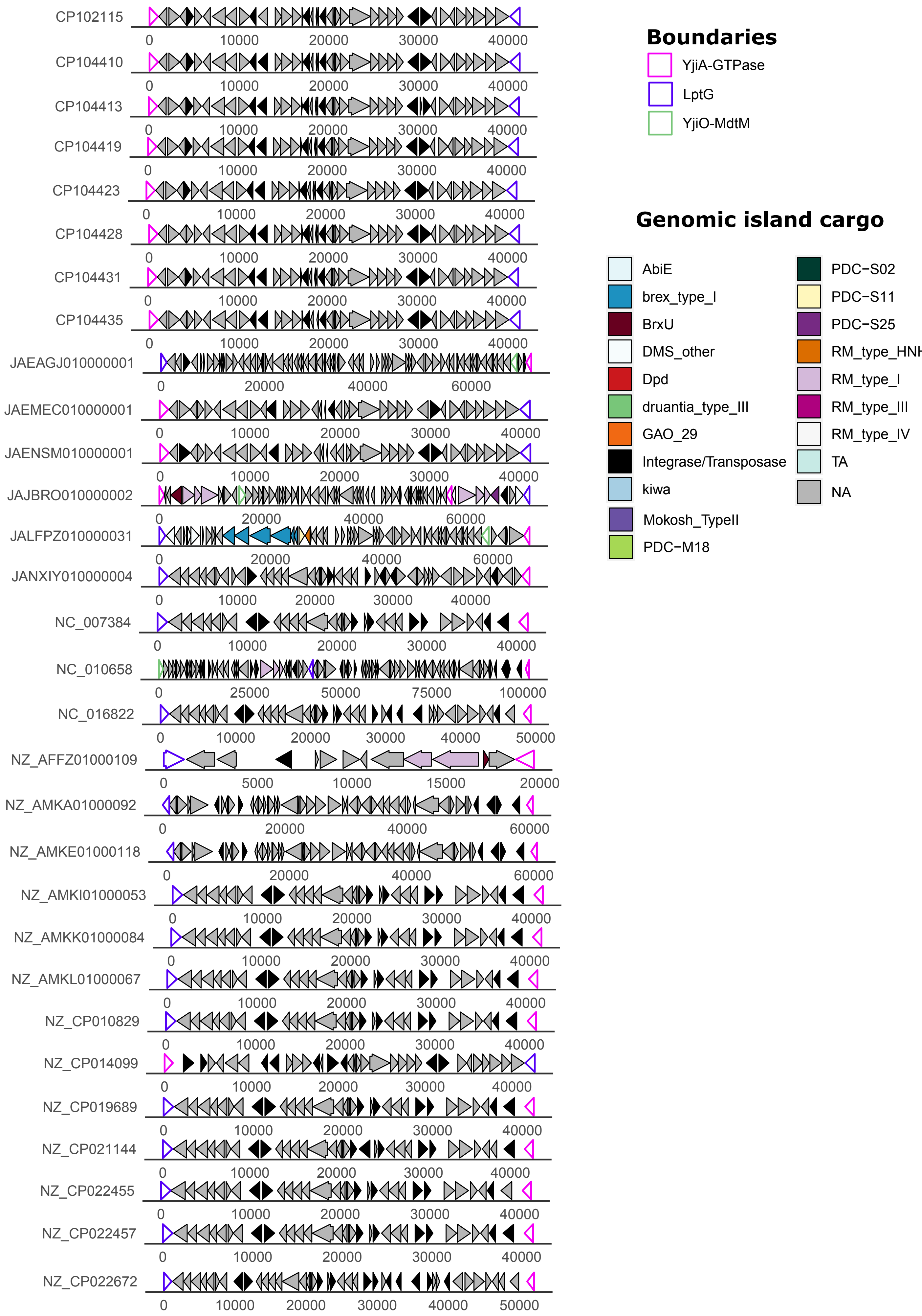
CP102115
CP104410
CP104413
CP104419
CP104423
CP104428
CP104431
CP104435
JAEAGJ010000001
JAEMEC010000001
JAENSM010000001
JAJBRO010000002
JALFPZ010000031
JANXIY010000004
NC_007384
NC_010658
NC_016822
NZ_AFFZ01000109
NZ_AMKA01000092
NZ_AMKE01000118
NZ_AMKI01000053
NZ_AMKK01000084
NZ_AMKL01000067
NZ_CP010829
NZ_CP014099
NZ_CP019689
NZ_CP021144
NZ_CP022455
NZ_CP022457
NZ_CP022672
Boundaries
YjiA-GTPase
LptG
YjiO-MdtM
Genomic island cargo
AbiE
brex_type_I
BrxU
DMS_other
Dpd
druantia_type_III
GAO_29
Integrase/Transposase
kiwa
Mokosh_TypeII
PDC−M18
PDC−S02
PDC−S11
PDC−S25
RM_type_HN
RM_type_I
RM_type_III
RM_type_IV
TA
NA

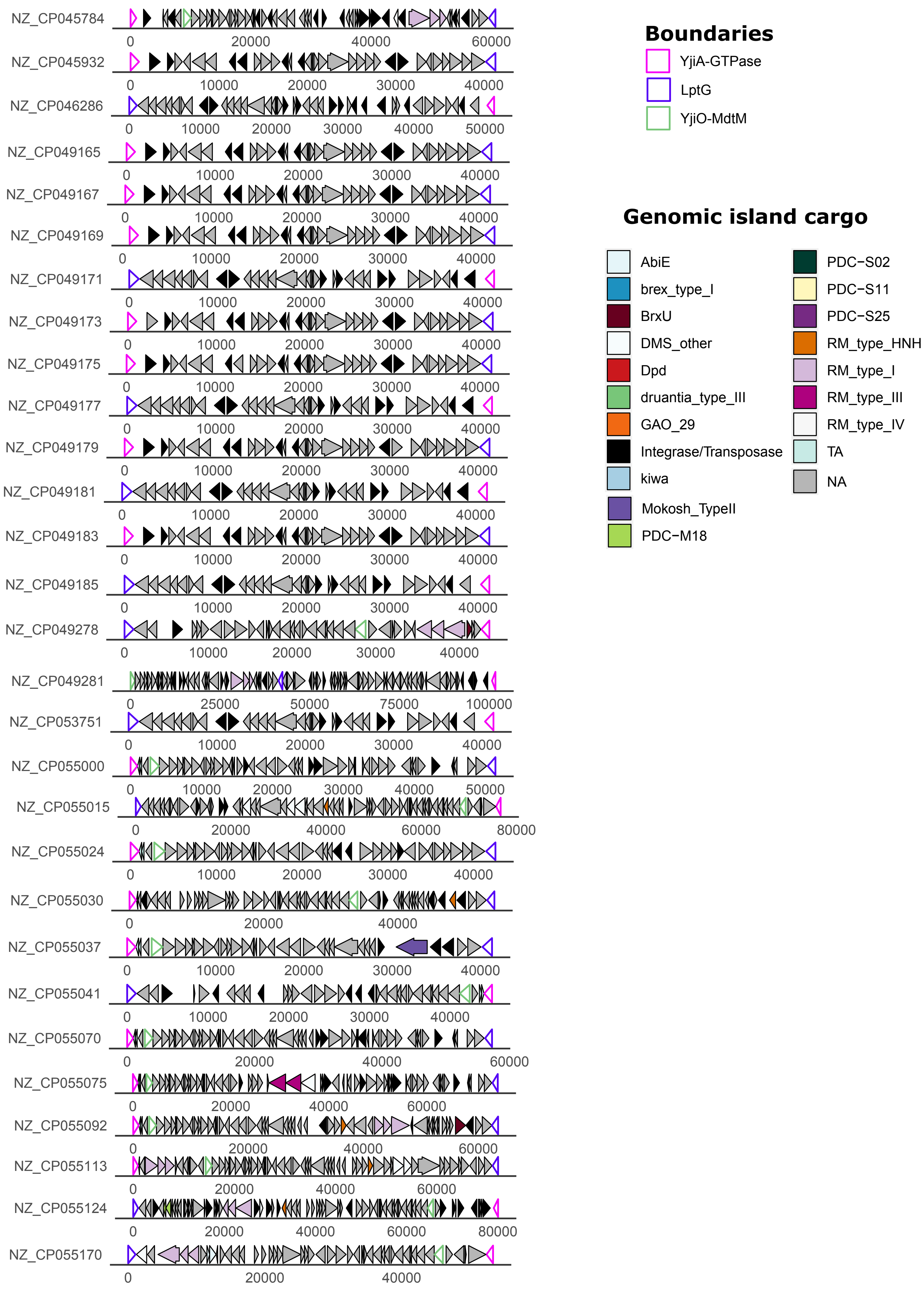

NZ_CP045784
NZ_CP045932
NZ_CP046286
NZ_CP049165
NZ_CP049167
NZ_CP049169
NZ_CP049171
NZ_CP049173
NZ_CP049175
NZ_CP049177
NZ_CP049179
NZ_CP049181
NZ_CP049183
NZ_CP049185
NZ_CP049278
NZ_CP049281
NZ_CP053751
NZ_CP055000
NZ_CP055015
NZ_CP055024
NZ_CP055030
NZ_CP055037
NZ_CP055041
NZ_CP055070
NZ_CP055075
NZ_CP055092
NZ_CP055113
NZ_CP055124
NZ_CP055170
Boundaries
YjiA-GTPase
LptG
YjiO-MdtM
Genomic island cargo
AbiE
brex_type_I
BrxU
DMS_other
Dpd
druantia_type_III
GAO_29
Integrase/Transposase
kiwa
Mokosh_TypeII
PDC-M18
PDC-S02
PDC-S11
PDC-S25
RM_type_HNH
RM_type_I
RM_type_III
RM_type_IV
TA
NA

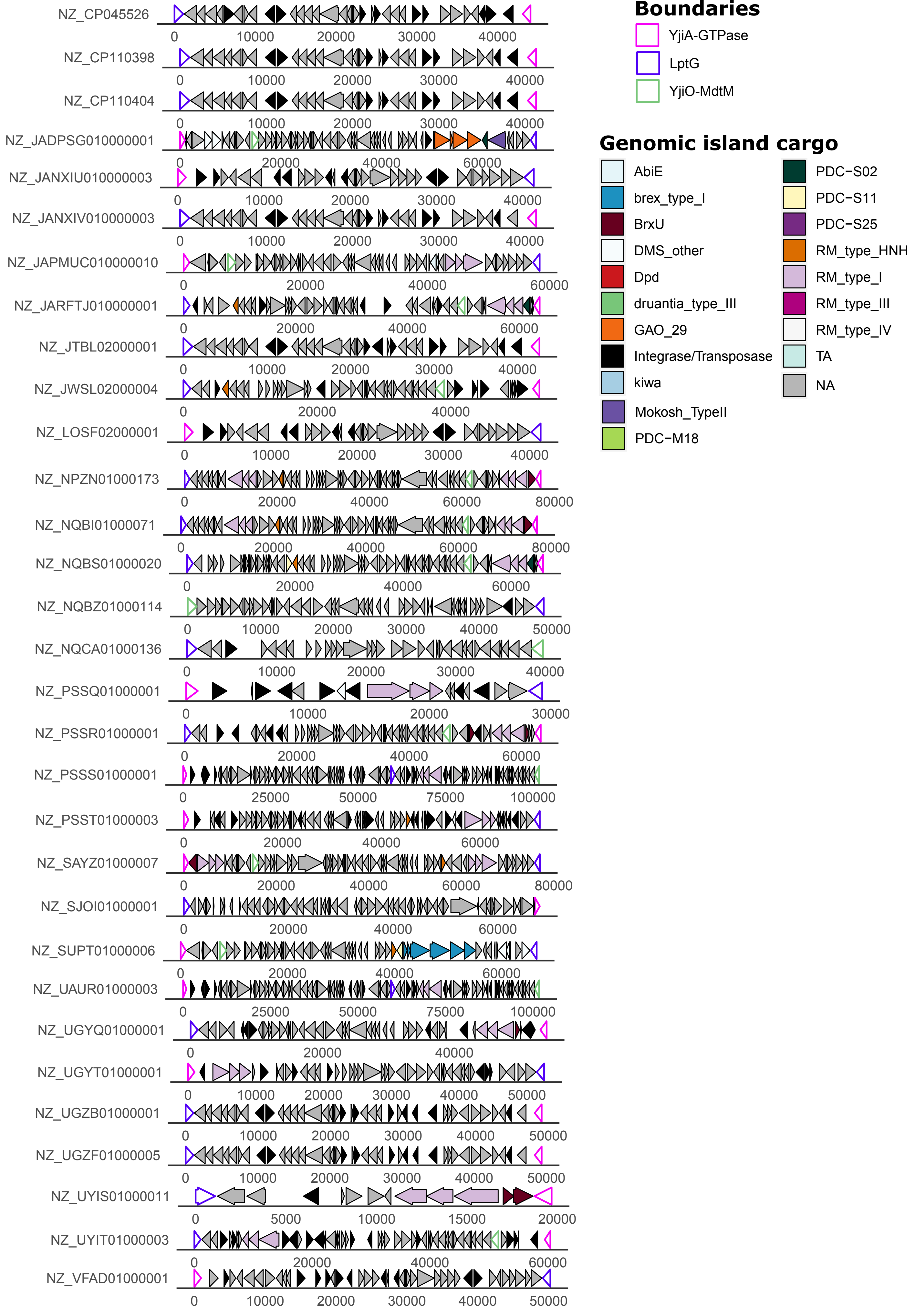

Boundaries
YjiA-GTPase
LptG
YjiO-MdtM
Genomic island cargo
AbiE
brex_type_I
BrxU
DMS_other
Dpd
druantia_type_III
GAO_29
Integrase/Transposase
kiwa
Mokosh_TypeII
PDC-M18
PDC-S02
PDC-S11
PDC-S25
RM_type_HNH
RM_type_I
RM_type_III
RM_type_IV
TA
NA
NZ_CP045526
NZ_CP110398
NZ_CP110404
NZ_JADPSG010000001
NZ_JANXIU010000003
NZ_JANXIV010000003
NZ_JAPMUC010000010
NZ_JARFTJ010000001
NZ_JTBL02000001
NZ_JWSL02000004
NZ_LOSF02000001
NZ_NPZN01000173
NZ_NQBI01000071
NZ_NQBS01000020
NZ_NQBZ01000114
NZ_NQCA01000136
NZ_PSSQ01000001
NZ_PSSR01000001
NZ_PSSS01000001
NZ_PSST01000003
NZ_SAYZ01000007
NZ_SJOI01000001
NZ_SUPT01000006
NZ_UAUR01000003
NZ_UGYQ01000001
NZ_UGYT01000001
NZ_UGZB01000001
NZ_UGZF01000005
NZ_UYIS01000011
NZ_UYIT01000003
NZ_VFAD01000001

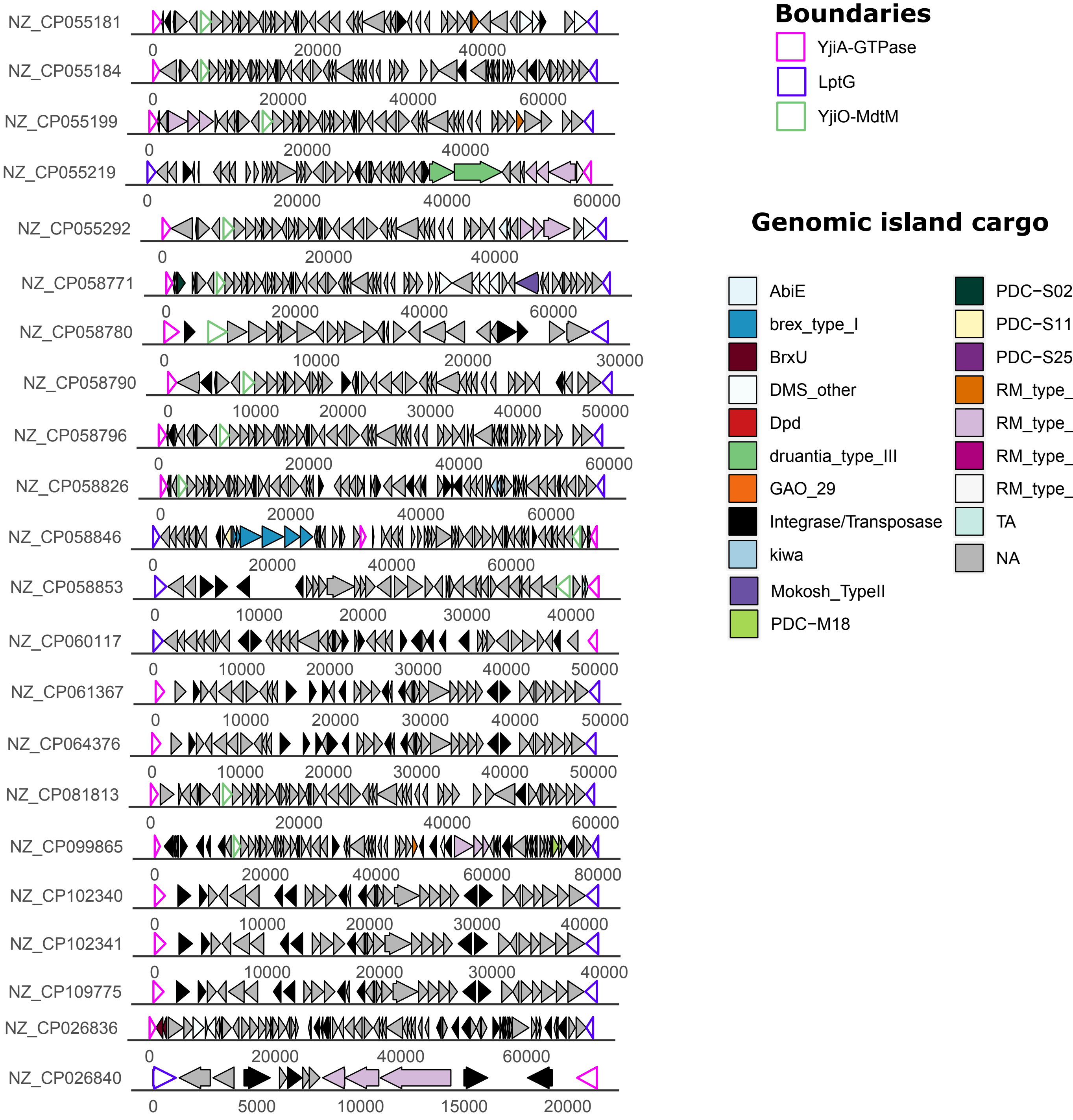


**Figure S8. LptG-YjiO-YjiA in *Shigella* spp accumulate numerous transposases and integrases.** Schematic representation of LptG-YjiO-YjiA islands in *Shigella* spp. Representative hotspots were chosen by filtering identical nucleotide sequences (90% sequence identity threshold) with fastANI. LptG, YjiO and YjiA boundaries are represented with coloured outlines. Anti-phage systems predicted by PADLOC are coloured as indicated in the legend. Integrases, transposases, toxin-antitoxin systems (TA), predicted T6SS-dependent effectors are also coloured according to the legend.

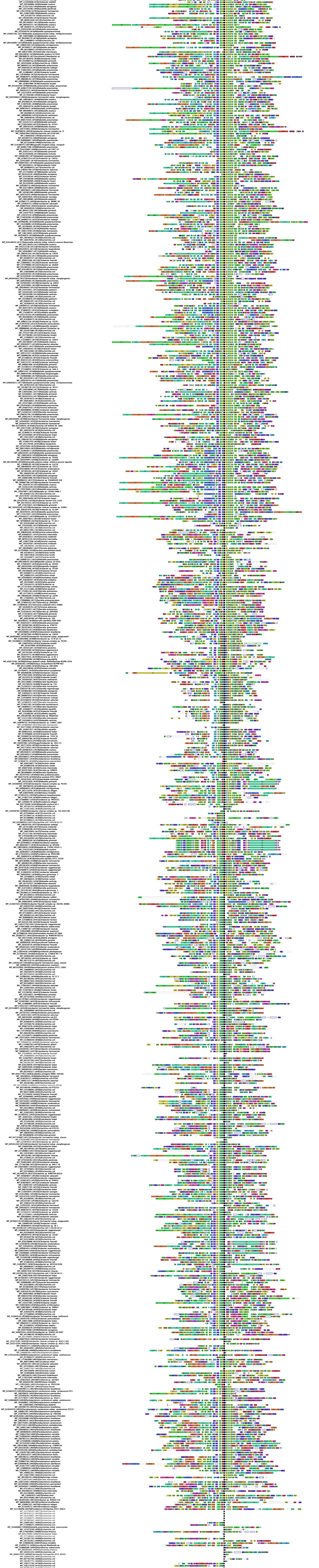

**Figure S9. SDIC4B homologues are often associated with BrxR.** FlaGs output showing the clustered genomic neighbourhood of SDCI4B homologues identified by PSI-BLAST.

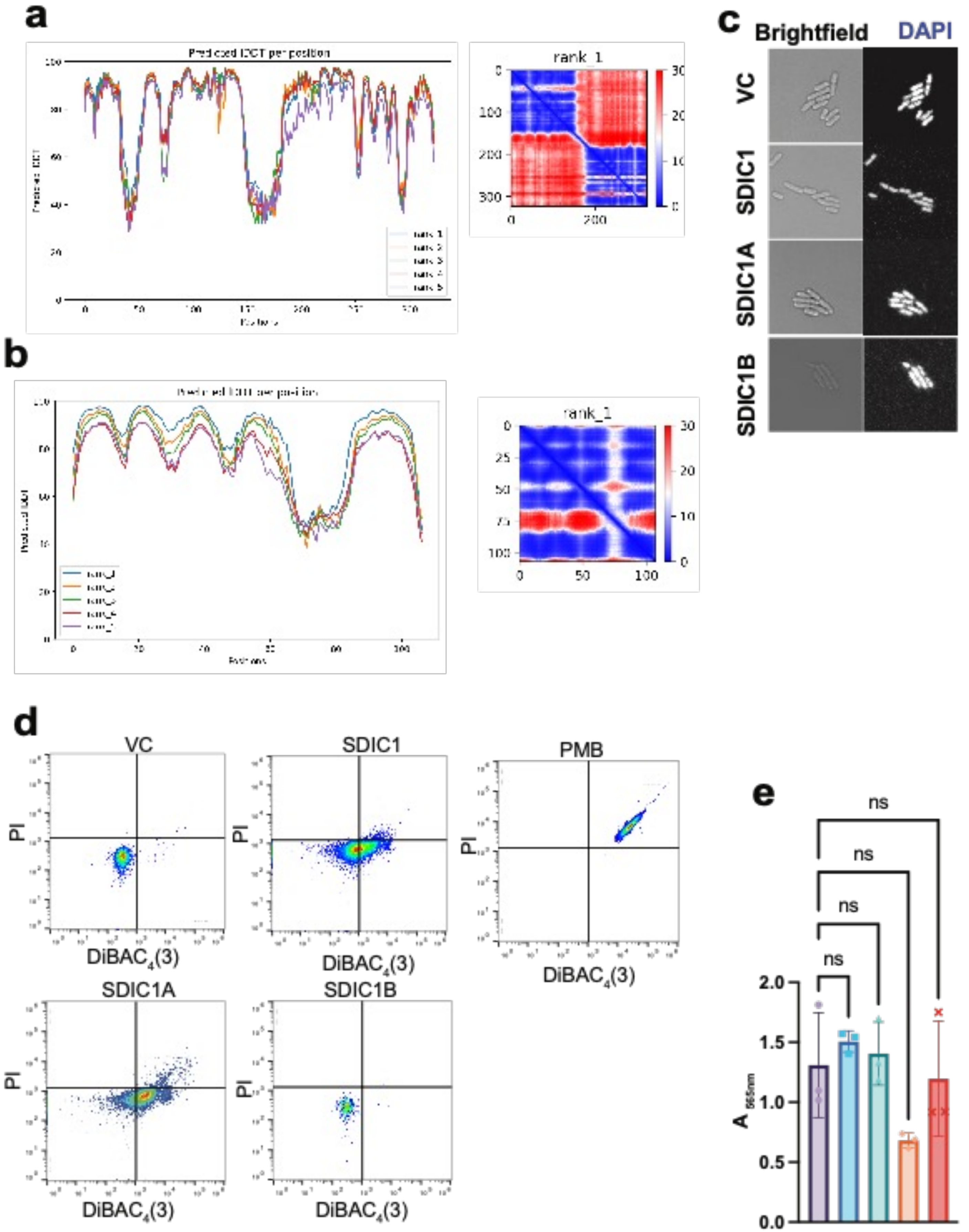

a
Predicted lDDT per position
Predicted lDDT
Positions
rank_1
b
Predicted lDDT per position
rank_1
rank_2
rank_3
rank_4
rank_5
Positions
rank_1
c
Brightfield
DAPI
VC
SDIC1
SDIC1A
SDIC1B
d
VC
SDIC1
PMB
SDIC1A
SDIC1B
PI
DiBAC4(3)
e
ns
ns
ns
ns
A565nm
VC
SDIC1
SDIC1B
SDIC1A
Shield II

**Figure S10. (a)** Local Distance Difference Test (lDDT) and Predicted Aligned Error (PAE) relative to SDIC1A predicted structure. **(b)** lDDT and PAE calculated for the predicted structure of SDIC1B. **(c)** Fluorescence microscopy analysis of *E. coli* MG1655 harbouring (pQE60-Tat) or the same plasmid encoding SDIC1, SDIC1A or SDIC1B when stained with DAPI, for DNA visualisation. **(d)** Representative replicate of flow cytometry analysis of *E. coli* MG1655 harbouring (pQE60-Tat) or the same plasmid encoding SDIC1, SDIC1A or SDIC1B when stained with propidium iodide (PI) and $DiBAC_4(3)$. Polymyxin B (PMB) was used as a control. **(e)** Whole cell measurement of $NAD^+$ levels for strains as in panel **b**. $NAD^+$ levels were measured using the EnzyChrom™ NAD/NADH Assay Kit (Universal Biologicals). Expression of the DNA-damaging anti-phage system Shield was used as a control to exclude $NAD^+$ depletion was a non-specific effect cell intoxication,
