## Supplementary Tables S28-29 for "Multi-conflict islands are a widespread trend within *Serratia* spp"

Table S28. Strains,phages and plasmids used in this study

| **Name** | **Description** | **Reference** |
| --- | --- | --- |
| **Strains** |  |  |
| *Serratia marcescens*  Db10 | Wild type | ^1^ |
| *Escherichia coli* |  |  |
| MG1655 | Wild type (model K-12 strain) | ^2^ |
| DH5α | Cloning strain, F– φ80lacZΔM15 Δ(lacZYA-argF)U169 recA1 endA1 hsdR17(rK–, mK+) phoA supE44 λ–thi-1 gyrA96 relA1 | New England Biolabs |
| **Phages** | **Description** | **Reference** |
| Emrys | Coliphage isolated in 50.784579, -1.162250 on *E.coli* DH5α as a host | This study |
| Lana | Coliphage isolated in 50.784657 -1.140652 on *E.coli* DH5α as a host | This study |
| Alwar | Coliphage isolated in 50.787564, -1.168700 on *E.coli* DH5α as a host | This study |
| phGM01 | Coliphage isolated from a waste water sample obtained from Guildford waste water treatment plant. Phages were enriched on *E.coli* DH5α as a host. | This study |
| phAvM | Coliphage isolated from a waste water sample obtained from Guildford waste water treatment plant. Phages were enriched on *E.coli* DH5α as a host. | This study |
| phNic | Coliphage isolated from a waste water sample obtained from Guildford waste water treatment plant. Phages were enriched on *E.coli* DH5α as a host. | This study |
| Riaz | Coliphage isolated from a waste water sample obtained from Guildford waste water treatment plant. Phages were enriched on *E.coli* DH5α as a host. | This study |
| Donnie | Coliphage isolated in 51.24355, -0.58543 on *E.coli* DH5α as a host | This study |
| Snape | *Serratia* phage isolated from a waste water sample obtained from Guildford waste water treatment plant. Phages were enriched on *S. marcescens* Db10 | This study |
| DpD | *Serratia* phage isolated from a waste water sample obtained from Guildford waste water treatment plant. Phages were enriched on *S. marcescens* Db10 | This study |
| Cap | *Serratia* phage isolated from a waste water sample obtained from Guildford waste water treatment plant. Phages were enriched on *S. marcescens* Db10 | This study |
| Kiryu | *Serratia* phage isolated from a waste water sample obtained from Guildford waste water treatment plant. Phages were enriched on *S. marcescens* Db10 | This study |
| Majima-phage | *Serratia* phage isolated from a waste water sample obtained from Guildford waste water treatment plant. Phages were enriched on *S. marcescens* Db10 | This study |
| Phagepool | *Serratia* phage isolated from a waste water sample obtained from Guildford waste water treatment plant. Phages were enriched on *S. marcescens* Db10 | This study |
| phGM1 | *Serratia* phage isolated from a waste water sample obtained from Guildford waste water treatment plant. Phages were enriched on *S. marcescens* Db10 | This study |
| phGM2 | *Serratia* phage isolated from a waste water sample obtained from Guildford waste water treatment plant. Phages were enriched on *S. marcescens* Db10 | This study |
| phGM3 | *Serratia* phage isolated from a waste water sample obtained from Guildford waste water treatment plant. Phages were enriched on *S. marcescens* Db10 | This study |
| phGM4 | *Serratia* phage isolated from a waste water sample obtained from Guildford waste water treatment plant. Phages were enriched on *S. marcescens* Db10 | This study |
| Olivia | *Serratia* phage isolated from a waste water sample obtained from Guildford waste water treatment plant. Phages were enriched on *S. marcescens* Db10 | This study |
| PM1259 | *Serratia* phage isolated from a waste water sample obtained from Guildford waste water treatment plant. Phages were enriched on *S. marcescens* Db10 | This study |
| Plaquezilla | *Serratia* phage isolated from a waste water sample obtained from Guildford waste water treatment plant. Phages were enriched on *S. marcescens* Db10 | This study |
| phGM7 | *Serratia* phage isolated from a waste water sample obtained from Guildford waste water treatment plant. Phages were enriched on *S. marcescens* Db10 | This study |
| Dante | *Serratia* phage isolated from a waste water sample obtained from Guildford waste water treatment plant. Phages were enriched on *S. marcescens* Db10 | This study |

| **Plasmids** | **Description** | **Reference** |
| --- | --- | --- |

| pQE60-Tat | Sequence of the constitutive Tat promoter cloned to replace the T5 promoter of pQE60 | This study/Genscript |
| --- | --- | --- |
| pGM248 | Coding sequence of SDIC3 (OCG46_RS13925 to OCG46_RS13900) from *Serratia entomophila* isolate (NZ_CAMKIA010000002.1) cloned in pQE60-Tat | This study/Genscript |
| pGM249 | Coding sequence of SDIC4 (AWY87_RS24860 and AWY87_RS12625) from *Serratia marcecens* strain 2880STDY5682973 (NZ_FCKZ01000002.1) cloned in pQE60-Tat | This study/Genscript |
| pGM250 | Coding sequence of SDIC4A (AWY87_RS24860) from *Serratia marcecens* strain 2880STDY5682973 (NZ_FCKZ01000002.1) cloned in pQE60-Tat | This study/Genscript |
| pGM251 | Coding sequence of SDIC4B (AWY87_RS12625) from *Serratia marcecens* strain 2880STDY5682973 (NZ_FCKZ01000002.1) cloned in pQE60-Tat | This study/Genscript |
| pGM252 | Coding sequence of SDIC5 (BSQ40_RS30855 to BSQ40_RS24190) from *Serratia fonticola* strain 5l (NZ_MQRH01000029.1) cloned in pQE60-Tat | This study/Genscript |
| pGM255 | Coding sequence of SDIC1 (AWX46_RS05495 and AWX46_RS05490) from *Serratia marcescens* strain 2880STDY5682900 (NZ_FCKY01000006.1) cloned in pQE60-Tat | This study/Genscript |
| pGM256 | Coding sequence of SDIC2 (AWX26_RS24455 to AWX26_RS24470) *from Serratia marcescens* strain 2880STDY5683020 (NZ_FCGS01000003.1) cloned in pQE60-Tat | This study/Genscript |
| pGM263 | Deletion of SDIC1A(AWX46_RS05495) from pGM255 | This study |
| pGM264 | Deletion of SDIC1B (AWX46_RS05490) from pGM255 | This study |
| pUT18 | Bacterial Two Hybrid plasmid (for fusion of target protein with C-terminal T18 fragment of CyaA; Amp^R^) | ^3^ |
| pT25 | Bacterial Two Hybrid plasmid (for fusion of target protein with N-terminal T25 fragment of CyaA; Cm^R^) | ^3^ |
| pGM266 | Coding sequence of SDIC1A cloned in pUT18 | This study |
| pGM267 | Coding sequence of SDIC1A cloned in pT25 | This study |
| pGM268 | Coding sequence of SDIC1B cloned in pUT18 | This study |
| pGM269 | Coding sequence of SDIC1B cloned in pT25 | This study |
| pGM118 | Coding sequence for ShdB II (AN400_RS26695) in pUT18 | ^4^ |
| pGM120 | Coding sequence for ShdB II (AN400_RS26695) in pT25 | ^4^ |
| pGM278 | Deletion of aa in SDIC1A_1-103_ in plasmid pGM255 | This study |
| pGM279 | Deletion of aa in SDIC1A_104-323_ in plasmid pGM255 | This study |
| pGM280 | Deletion of aa in SDIC1A_1-103_ in plasmid pGM264 | This study |
| pGM281 | Deletion of aa in SDIC1A_104-323_ in plasmid pGM264 | This study |

Table S29. Oligonucleotide primers and additional details for plasmid construction.

| **Plasmid** | **Sequence of relevant primers (5’-3’)**^a^ | **Description** |
| --- | --- | --- |
| pQE60-Tat | CTTGGACTCCTGTTGATAG | Forward primer to delete SDIC1 in pGM255 and generate empty pQE60-Tat by KLD |
|  | TTAATTTCTCCTCTTTAATG | Reverse primer to delete SDIC1 from pGM255 and generate empty pQE60-Tat by KLD |
| pGM263 | TTAATTTCTCCTCTTTAATGAATTC | Forward primer to delete SDIC1A in pGM255 by KLD. |
|  | GAATTCGCTAGCCCAAAAAAAC | Reverse primer to delete SDIC1A in pGM255 by KLD. |
| pGM264 | CTTGGACTCCTGTTGATAG | Forward primer to delete SDIC1B in pGM255 by KLD. |
|  | TGTTGCCTCCCGACTCTCAG | Reverse primer to delete SDIC1B in pGM255 by KLD. |
| pGM266 | AGTCGACCTGCAGGCATG | Forward primer to clone SDICA in pUT18 by NEBuilder HiFi DNA Assembly. Primer is used to linearise vector |
|  | CTAAGTAATATGGTGCACTC | Reverse primer to clone SDICA in pUT18 by NEBuilder HiFi DNA Assembly.  Primer is used to linearise vector |
|  | TGCATGCCTGCAGGTCGACTATGGCTACCCTCGTTTTTTC | Forward primer to clone SDICA in pUT18 by NEBuilder HiFi DNA Assembly. |
|  | GAGTGCACCATATTACTTAGGGTCATACGTTGTTTTATGGATTTG | Reverse primer to clone SDICA in pUT18 by NEBuilder HiFi DNA Assembly. |
| pGM267 | AGTCGACCCTGCAGCCCG | Forward primer to clone SDICA in pT25 by NEBuilder HiFi DNA Assembly. Primer is used to linearise vector |
|  | CTAGAGGATCCCCGGGTAC | Reverse primer to clone SDICA in pT25 by NEBuilder HiFi DNA Assembly.  Primer is used to linearise vector |
|  | GGCGGGCTGCAGGGTCGACTATGGCTACCCTCGTTTTTTC | Forward primer to clone SDICA in pT25 by NEBuilder HiFi DNA Assembly. |
|  | GGTACCCGGGGATCCTCTAGTCAGGTCATACGTTGTTTTATG | Reverse primer to clone SDICA in pT25 by NEBuilder HiFi DNA Assembly. |
| pGM268 | AGTCGACCTGCAGGCATG | Forward primer to clone SDICB in pUT18 by NEBuilder HiFi DNA Assembly. Primer is used to linearise vector |
|  | CTAAGTAATATGGTGCACTC | Reverse primer to clone SDICB in pUT18 by NEBuilder HiFi DNA Assembly.  Primer is used to linearise vector |
|  | TGCATGCCTGCAGGTCGACTATGGTACAACTCAATCTG | Forward primer to clone SDICB in pUT18 by NEBuilder HiFi DNA Assembly. |
|  | GAGTGCACCATATTACTTAGTCTCTCAGAACAAAACTTTTCCTG | Reverse primer to clone SDICB in pUT18 by NEBuilder HiFi DNA Assembly. |
| pGM269 | AGTCGACCCTGCAGCCCG | Forward primer to clone SDICB in pT25 by NEBuilder HiFi DNA Assembly. Primer is used to linearise vector |
|  | CTAGAGGATCCCCGGGTAC | Reverse primer to clone SDICB in pT25 by NEBuilder HiFi DNA Assembly.  Primer is used to linearise vector |
|  | GGCGGGCTGCAGGGTCGACTATGGTACAACTCAATCTG | Forward primer to clone SDICB in pT25 by NEBuilder HiFi DNA Assembly. |
|  | ggtacccggggatcctctagTTATCTCTCAGAACAAAACTTTTC | Reverse primer to clone SDICB in pT25 by NEBuilder HiFi DNA Assembly. |
| pGM278 | TTAATTTCTCCTCTTTAATGAATTC | Forward primer to delete SDIC1A_1-103_in pGM255 or pGM264 by KLD. |
|  | ATTTTGCGCAATTGTGCCTG | Forward primer to delete SDIC1A_1-103_in pGM255 or pGM264 by KLD. |
| pGM279 | TTAATTTCTCCTCTTTAATGAATTC | Forward primer to delete SDIC1A_1-103_in pGM255 or pGM264 by KLD. |
|  | ATTTTGCGCAATTGTGCCTG | Forward primer to delete SDIC1A_1-103_in pGM255 or pGM264 by KLD. |
| pGM280 | CACCGGGATGACTATCGCTTC | Forward primer to delete SDIC1A_1-103_in pGM255 or pGM264 by KLD. |
|  | GAGTCGGGAGGCAACAATG | Forward primer to delete SDIC1A_1-103_in pGM255 or pGM264 by KLD. |
| pGM281 | CACCGGGATGACTATCGCTTC | Forward primer to delete SDIC1A_104-323_in pGM255 by KLD. |
|  | CTTGGACTCCTGTTGATAG | Forward primer to delete SDIC1A_104-323_in pGM255 by KLD. |

* For the following plasmids the full-length insert was synthetised by GenScript and the genes of interest were cloned in pBAD18 by NEBuilder HiFi DNA Assembly

| **Plasmids synthetised by Genscript** | **Sequence (5’-3’)**^a^ | **Description** |
| --- | --- | --- |
| pGM248 | CTCGAGAAATGTCGGTTGGCGCAAAACACGCTGATTTTTTCATCGCTCAAGGCGGGCCGTGTAACGTATAATGCGGCTTTGTTTAATCATCATCTACCACAGAGGATTTCACACAGAATTCATTAAAGAGGAGAAATTAAATGCGATTGCCTTACGTAATATTTACAGCAGCTATACTGGCTTCTGCATCAACAGCACATGCTATTCCCAATATGTGGTCGAGTGGTTTTGGGATGGGCGTGACCGAATATATCATTACCAGCCCGGATAAGGTGGTGTTTAATCTGAACTGTACTGGCAATCCGGATGCGCAGAATATTCTGCAACACGGTGTTCATCTCACCCTGCCTGATGGAACATCAGTAAGTTCTCATGACGACGGGACGGAAATAACGGTCGTCATGGATAACAGTCAGTATCCGTTACCGTCATTCCTCGGCTGGCGAAACGGTGACAATGCCTGGGTGTCATTTATTGACGCACTGGGGCAGGCAGCAAATTTCGATGTTTATGTCAGCGATAAAAAAGTAGGCACTTTCAGTCCGGGTCTGAAGAATACGCAAAAAGAGCTGTCTGACCTCAGTGAGTGCCGGACCACACACTACAGCGATTAGGCTTGGACTCCTGTTGATAGATCCAGTAATGACCTCAGAACTCCATCTGGATTTGTTCAGAACGCTCGGTTGCCGCCGGGCGTTTTTTATTGGTGAGAATCCAAGCTAGCTTGGCGAGATTTTCAGGAGCTAAGGAAGCTAAAATGGAGAAAAAAATCACTGGATATACCACCGTTGATATATCCCAATGGCATCGTAAAGAACATTTTGAGGCATTTCAGTCAGTTGCTCAATGTACCTATAACCAGACCGTTCAGCTGGATATTACGGCCTTTTTAAAGACCGTAAAGAAAAATAAGCACAAGTTTTATCCGGCCTTTATTCACATTCTTGCCCGCCTGATGAATGCTCATCCGGAATTTCGTATGGCAATGAAAGACGGTGAGCTGGTGATATGGGATAGTGTTCACCCTTGTTACACCGTTTTCCATGAGCAAACTGAAACGTTTTCATCGCTCTGGAGTGAATACCACGACGATTTCCGGCAGTTTCTACACATATATTCGCAAGATGTGGCGTGTTACGGTGAAAACCTGGCCTATTTCCCTAAAGGGTTTATTGAGAATATGTTTTTCGTCTCAGCCAATCCCTGGGTGAGTTTCACCAGTTTTGATTTAAACGTGGCCAATATGGACAACTTCTTCGCCCCCGTTTTCACCATGCATGGGCAAATATTATACGCAAGGCGACAAGGTGCTGATGCCGCTGGCGATTCAGGTTCATCATGCCGTCTGTGATGGCTTCCATGTCGGCAGAATGCTTAATGAATTACAACAGTACTGCGATGAGTGGCAGGGCGGGGCGTAATTTTTTTAAGGCAGTTATTGGTGCCCTTAAACGCCTGGGGTAATGACTCTCTAGCTTGAGGCATCAAATAAAACGAAAGGCTCAGTCGAAAGACTGGGCCTTTCGTTTTATCTGTTGTTTGTCGGTGAACGCTCTCCTGAGTAGGACAAATCCGCCGCTCTAGAGCTGCCTCGCGCGTTTCGGTGATGACGGTGAAAACCTCTGACACATGCAGCTCCCGGAGACGGTCACAGCTTGTCTGTAAGCGGATGCCGGGAGCAGACAAGCCCGTCAGGGCGCGTCAGCGGGTGTTGGCGGGTGTCGGGGCGCAGCCATGACCCAGTCACGTAGCGATAGCGGAGTGTATACTGGCTTAACTATGCGGCATCAGAGCAGATTGTACTGAGAGTGCACCATATGCGGTGTGAAATACCGCACAGATGCGTAAGGAGAAAATACCGCATCAGGCGCTCTTCCGCTTCCTCGCTCACTGACTCGCTGCGCTCGGTCGTTCGGCTGCGGCGAGCGGTATCAGCTCACTCAAAGGCGGTAATACGGTTATCCACAGAATCAGGGGATAACGCAGGAAAGAACATGTGAGCAAAAGGCCAGCAAAAGGCCAGGAACCGTAAAAAGGCCGCGTTGCTGGCGTTTTTCCATAGGCTCCGCCCCCCTGACGAGCATCACAAAAATCGACGCTCAAGTCAGAGGTGGCGAAACCCGACAGGACTATAAAGATACCAGGCGTTTCCCCCTGGAAGCTCCCTCGTGCGCTCTCCTGTTCCGACCCTGCCGCTTACCGGATACCTGTCCGCCTTTCTCCCTTCGGGAAGCGTGGCGCTTTCTCATAGCTCACGCTGTAGGTATCTCAGTTCGGTGTAGGTCGTTCGCTCCAAGCTGGGCTGTGTGCACGAACCCCCCGTTCAGCCCGACCGCTGCGCCTTATCCGGTAACTATCGTCTTGAGTCCAACCCGGTAAGACACGACTTATCGCCACTGGCAGCAGCCACTGGTAACAGGATTAGCAGAGCGAGGTATGTAGGCGGTGCTACAGAGTTCTTGAAGTGGTGGCCTAACTACGGCTACACTAGAAGGACAGTATTTGGTATCTGCGCTCTGCTGAAGCCAGTTACCTTCGGAAAAAGAGTTGGTAGCTCTTGATCCGGCAAACAAACCACCGCTGGTAGCGGTGGTTTTTTTGTTTGCAAGCAGCAGATTACGCGCAGAAAAAAAGGATCTCAAGAAGATCCTTTGATCTTTTCTACGGGGTCTGACGCTCAGTGGAACGAAAACTCACGTTAAGGGATTTTGGTCATGAGATTATCAAAAAGGATCTTCACCTAGATCCTTTTAAATTAAAAATGAAGTTTTAAATCAATCTAAAGTATATATGAGTAAACTTGGTCTGACAGTTACCAATGCTTAATCAGTGAGGCACCTATCTCAGCGATCTGTCTATTTCGTTCATCCATAGTTGCCTGACTCCCCGTCGTGTAGATAACTACGATACGGGAGGGCTTACCATCTGGCCCCAGTGCTGCAATGATACCGCGAGACCCACGCTCACCGGCTCCAGATTTATCAGCAATAAACCAGCCAGCCGGAAGGGCCGAGCGCAGAAGTGGTCCTGCAACTTTATCCGCCTCCATCCAGTCTATTAATTGTTGCCGGGAAGCTAGAGTAAGTAGTTCGCCAGTTAATAGTTTGCGCAACGTTGTTGCCATTGCTACAGGCATCGTGGTGTCACGCTCGTCGTTTGGTATGGCTTCATTCAGCTCCGGTTCCCAACGATCAAGGCGAGTTACATGATCCCCCATGTTGTGCAAAAAAGCGGTTAGCTCCTTCGGTCCTCCGATCGTTGTCAGAAGTAAGTTGGCCGCAGTGTTATCACTCATGGTTATGGCAGCACTGCATAATTCTCTTACTGTCATGCCATCCGTAAGATGCTTTTCTGTGACTGGTGAGTACTCAACCAAGTCATTCTGAGAATAGTGTATGCGGCGACCGAGTTGCTCTTGCCCGGCGTCAATACGGGATAATACCGCGCCACATAGCAGAACTTTAAAAGTGCTCATCATTGGAAAACGTTCTTCGGGGCGAAAACTCTCAAGGATCTTACCGCTGTTGAGATCCAGTTCGATGTAACCCACTCGTGCACCCAACTGATCTTCAGCATCTTTTACTTTCACCAGCGTTTCTGGGTGAGCAAAAACAGGAAGGCAAAATGCCGCAAAAAAGGGAATAAGGGCGACACGGAAATGTTGAATACTCATACTCTTCCTTTTTCAATATTATTGAAGCATTTATCAGGGTTATTGTCTCATGAGCGGATACATATTTGAATGTATTTAGAAAAATAAACAAATAGGGGTTCCGCGCACATTTCCCCGAAAAGTGCCACCTGACGTCTAAGAAACCATTATTATCATGACATTAACCTATAAAAATAGGCGTATCACGAGGCCCTTTCGTCTTCAC | A Tat promoter was inserted to replace the original T5 promoter in plasmid pQE60. SDIC3 was cloned downstream of the Tat promoter. |
| pGM249 | CTCGAGAAATGTCGGTTGGCGCAAAACACGCTGATTTTTTCATCGCTCAAGGCGGGCCGTGTAACGTATAATGCGGCTTTGTTTAATCATCATCTACCACAGAGGATTTCACACAGAATTCATTAAAGAGGAGAAATTAAATGAAAGAATTGAAATTGTTGTCTTTACTGCCAGTCATATTGTTCCCCTTACTTATCGCAGGTAATGCTCACGCAGCACGTCCTTATAGCGGATTCCTGCCTGCAGCAGTGAATAATAAGAGTGGGAATATATTCTGGTGTGATGCCCACCGCCAGGTCAGAAAACTGTGTACAGCGACGGAAGTGGATTCAATGGGATGGTTTATTGTTGTTCGCTCGAAACCTTCCAGAGATTGTCCGGGGGGCCTTTGGGGGTGGATCACTCCCAGTAAAGACGATGTAGCAACAGGGTATACCATTACGCCAGGTAAAGAATTCAATCTTACCATGAAGGATATGTGCAGTCCGGATATGAAAGTGACGTTCCGCCCGAATAAAAAGGACCCTCGTTATGACGATCTCGTTGGGATTTATGCTGGGAAAGAAGCGTTTCGCTACAAACAAATAAAATAATCCAACTGTTTATTCCTTACTTTATTGTTTTGAACAACAAAAGCTTTATTACATGATTTGAGGACAACATTATGCGATTGCCTTACGTAATATTTACAGCAGCTATACTGGCTTCTGCATCAACAGCACATGCTATTCCCAATATGTGGTCGAGTGGTTTTGGGATGGGCGTGACCGAATATATCATTACCAGCCCGGATAAGGTGGTGTTTAATCTGAACTGTACTGGCAATCCGGATGCGCAGAATATTCTGCAACACGGTGTTCATCTCACCCTGCCTGATGGAACATCAGTAAGTTCTCATGACGACGGGACGGAAATAACGGTCGTCATGGATAACAGTCAGTATCCGTTACCGTCATTCCTCGGCTGGCGAAACGGTGACAATGCCTGGGTGTCATTTATTGACGCACTGGGGCAGGCAGCAAATTTCGATGTTTATGTCAGCGATAAAAAAGTAGGCACTTTCAGTCCGGGTCTGAAGAATACGCAAAAAGAGCTGTCTGACCTCAGTGAGTGCCGGACCACACACTACAGCGATTAGGCTTGGACTCCTGTTGATAGATCCAGTAATGACCTCAGAACTCCATCTGGATTTGTTCAGAACGCTCGGTTGCCGCCGGGCGTTTTTTATTGGTGAGAATCCAAGCTAGCTTGGCGAGATTTTCAGGAGCTAAGGAAGCTAAAATGGAGAAAAAAATCACTGGATATACCACCGTTGATATATCCCAATGGCATCGTAAAGAACATTTTGAGGCATTTCAGTCAGTTGCTCAATGTACCTATAACCAGACCGTTCAGCTGGATATTACGGCCTTTTTAAAGACCGTAAAGAAAAATAAGCACAAGTTTTATCCGGCCTTTATTCACATTCTTGCCCGCCTGATGAATGCTCATCCGGAATTTCGTATGGCAATGAAAGACGGTGAGCTGGTGATATGGGATAGTGTTCACCCTTGTTACACCGTTTTCCATGAGCAAACTGAAACGTTTTCATCGCTCTGGAGTGAATACCACGACGATTTCCGGCAGTTTCTACACATATATTCGCAAGATGTGGCGTGTTACGGTGAAAACCTGGCCTATTTCCCTAAAGGGTTTATTGAGAATATGTTTTTCGTCTCAGCCAATCCCTGGGTGAGTTTCACCAGTTTTGATTTAAACGTGGCCAATATGGACAACTTCTTCGCCCCCGTTTTCACCATGCATGGGCAAATATTATACGCAAGGCGACAAGGTGCTGATGCCGCTGGCGATTCAGGTTCATCATGCCGTCTGTGATGGCTTCCATGTCGGCAGAATGCTTAATGAATTACAACAGTACTGCGATGAGTGGCAGGGCGGGGCGTAATTTTTTTAAGGCAGTTATTGGTGCCCTTAAACGCCTGGGGTAATGACTCTCTAGCTTGAGGCATCAAATAAAACGAAAGGCTCAGTCGAAAGACTGGGCCTTTCGTTTTATCTGTTGTTTGTCGGTGAACGCTCTCCTGAGTAGGACAAATCCGCCGCTCTAGAGCTGCCTCGCGCGTTTCGGTGATGACGGTGAAAACCTCTGACACATGCAGCTCCCGGAGACGGTCACAGCTTGTCTGTAAGCGGATGCCGGGAGCAGACAAGCCCGTCAGGGCGCGTCAGCGGGTGTTGGCGGGTGTCGGGGCGCAGCCATGACCCAGTCACGTAGCGATAGCGGAGTGTATACTGGCTTAACTATGCGGCATCAGAGCAGATTGTACTGAGAGTGCACCATATGCGGTGTGAAATACCGCACAGATGCGTAAGGAGAAAATACCGCATCAGGCGCTCTTCCGCTTCCTCGCTCACTGACTCGCTGCGCTCGGTCGTTCGGCTGCGGCGAGCGGTATCAGCTCACTCAAAGGCGGTAATACGGTTATCCACAGAATCAGGGGATAACGCAGGAAAGAACATGTGAGCAAAAGGCCAGCAAAAGGCCAGGAACCGTAAAAAGGCCGCGTTGCTGGCGTTTTTCCATAGGCTCCGCCCCCCTGACGAGCATCACAAAAATCGACGCTCAAGTCAGAGGTGGCGAAACCCGACAGGACTATAAAGATACCAGGCGTTTCCCCCTGGAAGCTCCCTCGTGCGCTCTCCTGTTCCGACCCTGCCGCTTACCGGATACCTGTCCGCCTTTCTCCCTTCGGGAAGCGTGGCGCTTTCTCATAGCTCACGCTGTAGGTATCTCAGTTCGGTGTAGGTCGTTCGCTCCAAGCTGGGCTGTGTGCACGAACCCCCCGTTCAGCCCGACCGCTGCGCCTTATCCGGTAACTATCGTCTTGAGTCCAACCCGGTAAGACACGACTTATCGCCACTGGCAGCAGCCACTGGTAACAGGATTAGCAGAGCGAGGTATGTAGGCGGTGCTACAGAGTTCTTGAAGTGGTGGCCTAACTACGGCTACACTAGAAGGACAGTATTTGGTATCTGCGCTCTGCTGAAGCCAGTTACCTTCGGAAAAAGAGTTGGTAGCTCTTGATCCGGCAAACAAACCACCGCTGGTAGCGGTGGTTTTTTTGTTTGCAAGCAGCAGATTACGCGCAGAAAAAAAGGATCTCAAGAAGATCCTTTGATCTTTTCTACGGGGTCTGACGCTCAGTGGAACGAAAACTCACGTTAAGGGATTTTGGTCATGAGATTATCAAAAAGGATCTTCACCTAGATCCTTTTAAATTAAAAATGAAGTTTTAAATCAATCTAAAGTATATATGAGTAAACTTGGTCTGACAGTTACCAATGCTTAATCAGTGAGGCACCTATCTCAGCGATCTGTCTATTTCGTTCATCCATAGTTGCCTGACTCCCCGTCGTGTAGATAACTACGATACGGGAGGGCTTACCATCTGGCCCCAGTGCTGCAATGATACCGCGAGACCCACGCTCACCGGCTCCAGATTTATCAGCAATAAACCAGCCAGCCGGAAGGGCCGAGCGCAGAAGTGGTCCTGCAACTTTATCCGCCTCCATCCAGTCTATTAATTGTTGCCGGGAAGCTAGAGTAAGTAGTTCGCCAGTTAATAGTTTGCGCAACGTTGTTGCCATTGCTACAGGCATCGTGGTGTCACGCTCGTCGTTTGGTATGGCTTCATTCAGCTCCGGTTCCCAACGATCAAGGCGAGTTACATGATCCCCCATGTTGTGCAAAAAAGCGGTTAGCTCCTTCGGTCCTCCGATCGTTGTCAGAAGTAAGTTGGCCGCAGTGTTATCACTCATGGTTATGGCAGCACTGCATAATTCTCTTACTGTCATGCCATCCGTAAGATGCTTTTCTGTGACTGGTGAGTACTCAACCAAGTCATTCTGAGAATAGTGTATGCGGCGACCGAGTTGCTCTTGCCCGGCGTCAATACGGGATAATACCGCGCCACATAGCAGAACTTTAAAAGTGCTCATCATTGGAAAACGTTCTTCGGGGCGAAAACTCTCAAGGATCTTACCGCTGTTGAGATCCAGTTCGATGTAACCCACTCGTGCACCCAACTGATCTTCAGCATCTTTTACTTTCACCAGCGTTTCTGGGTGAGCAAAAACAGGAAGGCAAAATGCCGCAAAAAAGGGAATAAGGGCGACACGGAAATGTTGAATACTCATACTCTTCCTTTTTCAATATTATTGAAGCATTTATCAGGGTTATTGTCTCATGAGCGGATACATATTTGAATGTATTTAGAAAAATAAACAAATAGGGGTTCCGCGCACATTTCCCCGAAAAGTGCCACCTGACGTCTAAGAAACCATTATTATCATGACATTAACCTATAAAAATAGGCGTATCACGAGGCCCTTTCGTCTTCAC | A Tat promoter was inserted to replace the original T5 promoter in plasmid pQE60. SDIC4 was cloned downstream of the Tat promoter. |
| pGM250 | CTCGAGAAATGTCGGTTGGCGCAAAACACGCTGATTTTTTCATCGCTCAAGGCGGGCCGTGTAACGTATAATGCGGCTTTGTTTAATCATCATCTACCACAGAGGATTTCACACAGAATTCATTAAAGAGGAGAAATTAAATGAAAGAATTGAAATTGTTGTCTTTACTGCCAGTCATATTGTTCCCCTTACTTATCGCAGGTAATGCTCACGCAGCACGTCCTTATAGCGGATTCCTGCCTGCAGCAGTGAATAATAAGAGTGGGAATATATTCTGGTGTGATGCCCACCGCCAGGTCAGAAAACTGTGTACAGCGACGGAAGTGGATTCAATGGGATGGTTTATTGTTGTTCGCTCGAAACCTTCCAGAGATTGTCCGGGGGGCCTTTGGGGGTGGATCACTCCCAGTAAAGACGATGTAGCAACAGGGTATACCATTACGCCAGGTAAAGAATTCAATCTTACCATGAAGGATATGTGCAGTCCGGATATGAAAGTGACGTTCCGCCCGAATAAAAAGGACCCTCGTTATGACGATCTCGTTGGGATTTATGCTGGGAAAGAAGCGTTTCGCTACAAACAAATAAAATAAGCTTGGACTCCTGTTGATAGATCCAGTAATGACCTCAGAACTCCATCTGGATTTGTTCAGAACGCTCGGTTGCCGCCGGGCGTTTTTTATTGGTGAGAATCCAAGCTAGCTTGGCGAGATTTTCAGGAGCTAAGGAAGCTAAAATGGAGAAAAAAATCACTGGATATACCACCGTTGATATATCCCAATGGCATCGTAAAGAACATTTTGAGGCATTTCAGTCAGTTGCTCAATGTACCTATAACCAGACCGTTCAGCTGGATATTACGGCCTTTTTAAAGACCGTAAAGAAAAATAAGCACAAGTTTTATCCGGCCTTTATTCACATTCTTGCCCGCCTGATGAATGCTCATCCGGAATTTCGTATGGCAATGAAAGACGGTGAGCTGGTGATATGGGATAGTGTTCACCCTTGTTACACCGTTTTCCATGAGCAAACTGAAACGTTTTCATCGCTCTGGAGTGAATACCACGACGATTTCCGGCAGTTTCTACACATATATTCGCAAGATGTGGCGTGTTACGGTGAAAACCTGGCCTATTTCCCTAAAGGGTTTATTGAGAATATGTTTTTCGTCTCAGCCAATCCCTGGGTGAGTTTCACCAGTTTTGATTTAAACGTGGCCAATATGGACAACTTCTTCGCCCCCGTTTTCACCATGCATGGGCAAATATTATACGCAAGGCGACAAGGTGCTGATGCCGCTGGCGATTCAGGTTCATCATGCCGTCTGTGATGGCTTCCATGTCGGCAGAATGCTTAATGAATTACAACAGTACTGCGATGAGTGGCAGGGCGGGGCGTAATTTTTTTAAGGCAGTTATTGGTGCCCTTAAACGCCTGGGGTAATGACTCTCTAGCTTGAGGCATCAAATAAAACGAAAGGCTCAGTCGAAAGACTGGGCCTTTCGTTTTATCTGTTGTTTGTCGGTGAACGCTCTCCTGAGTAGGACAAATCCGCCGCTCTAGAGCTGCCTCGCGCGTTTCGGTGATGACGGTGAAAACCTCTGACACATGCAGCTCCCGGAGACGGTCACAGCTTGTCTGTAAGCGGATGCCGGGAGCAGACAAGCCCGTCAGGGCGCGTCAGCGGGTGTTGGCGGGTGTCGGGGCGCAGCCATGACCCAGTCACGTAGCGATAGCGGAGTGTATACTGGCTTAACTATGCGGCATCAGAGCAGATTGTACTGAGAGTGCACCATATGCGGTGTGAAATACCGCACAGATGCGTAAGGAGAAAATACCGCATCAGGCGCTCTTCCGCTTCCTCGCTCACTGACTCGCTGCGCTCGGTCGTTCGGCTGCGGCGAGCGGTATCAGCTCACTCAAAGGCGGTAATACGGTTATCCACAGAATCAGGGGATAACGCAGGAAAGAACATGTGAGCAAAAGGCCAGCAAAAGGCCAGGAACCGTAAAAAGGCCGCGTTGCTGGCGTTTTTCCATAGGCTCCGCCCCCCTGACGAGCATCACAAAAATCGACGCTCAAGTCAGAGGTGGCGAAACCCGACAGGACTATAAAGATACCAGGCGTTTCCCCCTGGAAGCTCCCTCGTGCGCTCTCCTGTTCCGACCCTGCCGCTTACCGGATACCTGTCCGCCTTTCTCCCTTCGGGAAGCGTGGCGCTTTCTCATAGCTCACGCTGTAGGTATCTCAGTTCGGTGTAGGTCGTTCGCTCCAAGCTGGGCTGTGTGCACGAACCCCCCGTTCAGCCCGACCGCTGCGCCTTATCCGGTAACTATCGTCTTGAGTCCAACCCGGTAAGACACGACTTATCGCCACTGGCAGCAGCCACTGGTAACAGGATTAGCAGAGCGAGGTATGTAGGCGGTGCTACAGAGTTCTTGAAGTGGTGGCCTAACTACGGCTACACTAGAAGGACAGTATTTGGTATCTGCGCTCTGCTGAAGCCAGTTACCTTCGGAAAAAGAGTTGGTAGCTCTTGATCCGGCAAACAAACCACCGCTGGTAGCGGTGGTTTTTTTGTTTGCAAGCAGCAGATTACGCGCAGAAAAAAAGGATCTCAAGAAGATCCTTTGATCTTTTCTACGGGGTCTGACGCTCAGTGGAACGAAAACTCACGTTAAGGGATTTTGGTCATGAGATTATCAAAAAGGATCTTCACCTAGATCCTTTTAAATTAAAAATGAAGTTTTAAATCAATCTAAAGTATATATGAGTAAACTTGGTCTGACAGTTACCAATGCTTAATCAGTGAGGCACCTATCTCAGCGATCTGTCTATTTCGTTCATCCATAGTTGCCTGACTCCCCGTCGTGTAGATAACTACGATACGGGAGGGCTTACCATCTGGCCCCAGTGCTGCAATGATACCGCGAGACCCACGCTCACCGGCTCCAGATTTATCAGCAATAAACCAGCCAGCCGGAAGGGCCGAGCGCAGAAGTGGTCCTGCAACTTTATCCGCCTCCATCCAGTCTATTAATTGTTGCCGGGAAGCTAGAGTAAGTAGTTCGCCAGTTAATAGTTTGCGCAACGTTGTTGCCATTGCTACAGGCATCGTGGTGTCACGCTCGTCGTTTGGTATGGCTTCATTCAGCTCCGGTTCCCAACGATCAAGGCGAGTTACATGATCCCCCATGTTGTGCAAAAAAGCGGTTAGCTCCTTCGGTCCTCCGATCGTTGTCAGAAGTAAGTTGGCCGCAGTGTTATCACTCATGGTTATGGCAGCACTGCATAATTCTCTTACTGTCATGCCATCCGTAAGATGCTTTTCTGTGACTGGTGAGTACTCAACCAAGTCATTCTGAGAATAGTGTATGCGGCGACCGAGTTGCTCTTGCCCGGCGTCAATACGGGATAATACCGCGCCACATAGCAGAACTTTAAAAGTGCTCATCATTGGAAAACGTTCTTCGGGGCGAAAACTCTCAAGGATCTTACCGCTGTTGAGATCCAGTTCGATGTAACCCACTCGTGCACCCAACTGATCTTCAGCATCTTTTACTTTCACCAGCGTTTCTGGGTGAGCAAAAACAGGAAGGCAAAATGCCGCAAAAAAGGGAATAAGGGCGACACGGAAATGTTGAATACTCATACTCTTCCTTTTTCAATATTATTGAAGCATTTATCAGGGTTATTGTCTCATGAGCGGATACATATTTGAATGTATTTAGAAAAATAAACAAATAGGGGTTCCGCGCACATTTCCCCGAAAAGTGCCACCTGACGTCTAAGAAACCATTATTATCATGACATTAACCTATAAAAATAGGCGTATCACGAGGCCCTTTCGTCTTCAC | A Tat promoter was inserted to replace the original T5 promoter in plasmid pQE60. SDIC4A was cloned downstream of the Tat promoter. |
| pGM251 | CTCGAGAAATGTCGGTTGGCGCAAAACACGCTGATTTTTTCATCGCTCAAGGCGGGCCGTGTAACGTATAATGCGGCTTTGTTTAATCATCATCTACCACAGAGGATTTCACACAGAATTCATTAAAGAGGAGAAATTAAATGCGATTGCCTTACGTAATATTTACAGCAGCTATACTGGCTTCTGCATCAACAGCACATGCTATTCCCAATATGTGGTCGAGTGGTTTTGGGATGGGCGTGACCGAATATATCATTACCAGCCCGGATAAGGTGGTGTTTAATCTGAACTGTACTGGCAATCCGGATGCGCAGAATATTCTGCAACACGGTGTTCATCTCACCCTGCCTGATGGAACATCAGTAAGTTCTCATGACGACGGGACGGAAATAACGGTCGTCATGGATAACAGTCAGTATCCGTTACCGTCATTCCTCGGCTGGCGAAACGGTGACAATGCCTGGGTGTCATTTATTGACGCACTGGGGCAGGCAGCAAATTTCGATGTTTATGTCAGCGATAAAAAAGTAGGCACTTTCAGTCCGGGTCTGAAGAATACGCAAAAAGAGCTGTCTGACCTCAGTGAGTGCCGGACCACACACTACAGCGATTAGGCTTGGACTCCTGTTGATAGATCCAGTAATGACCTCAGAACTCCATCTGGATTTGTTCAGAACGCTCGGTTGCCGCCGGGCGTTTTTTATTGGTGAGAATCCAAGCTAGCTTGGCGAGATTTTCAGGAGCTAAGGAAGCTAAAATGGAGAAAAAAATCACTGGATATACCACCGTTGATATATCCCAATGGCATCGTAAAGAACATTTTGAGGCATTTCAGTCAGTTGCTCAATGTACCTATAACCAGACCGTTCAGCTGGATATTACGGCCTTTTTAAAGACCGTAAAGAAAAATAAGCACAAGTTTTATCCGGCCTTTATTCACATTCTTGCCCGCCTGATGAATGCTCATCCGGAATTTCGTATGGCAATGAAAGACGGTGAGCTGGTGATATGGGATAGTGTTCACCCTTGTTACACCGTTTTCCATGAGCAAACTGAAACGTTTTCATCGCTCTGGAGTGAATACCACGACGATTTCCGGCAGTTTCTACACATATATTCGCAAGATGTGGCGTGTTACGGTGAAAACCTGGCCTATTTCCCTAAAGGGTTTATTGAGAATATGTTTTTCGTCTCAGCCAATCCCTGGGTGAGTTTCACCAGTTTTGATTTAAACGTGGCCAATATGGACAACTTCTTCGCCCCCGTTTTCACCATGCATGGGCAAATATTATACGCAAGGCGACAAGGTGCTGATGCCGCTGGCGATTCAGGTTCATCATGCCGTCTGTGATGGCTTCCATGTCGGCAGAATGCTTAATGAATTACAACAGTACTGCGATGAGTGGCAGGGCGGGGCGTAATTTTTTTAAGGCAGTTATTGGTGCCCTTAAACGCCTGGGGTAATGACTCTCTAGCTTGAGGCATCAAATAAAACGAAAGGCTCAGTCGAAAGACTGGGCCTTTCGTTTTATCTGTTGTTTGTCGGTGAACGCTCTCCTGAGTAGGACAAATCCGCCGCTCTAGAGCTGCCTCGCGCGTTTCGGTGATGACGGTGAAAACCTCTGACACATGCAGCTCCCGGAGACGGTCACAGCTTGTCTGTAAGCGGATGCCGGGAGCAGACAAGCCCGTCAGGGCGCGTCAGCGGGTGTTGGCGGGTGTCGGGGCGCAGCCATGACCCAGTCACGTAGCGATAGCGGAGTGTATACTGGCTTAACTATGCGGCATCAGAGCAGATTGTACTGAGAGTGCACCATATGCGGTGTGAAATACCGCACAGATGCGTAAGGAGAAAATACCGCATCAGGCGCTCTTCCGCTTCCTCGCTCACTGACTCGCTGCGCTCGGTCGTTCGGCTGCGGCGAGCGGTATCAGCTCACTCAAAGGCGGTAATACGGTTATCCACAGAATCAGGGGATAACGCAGGAAAGAACATGTGAGCAAAAGGCCAGCAAAAGGCCAGGAACCGTAAAAAGGCCGCGTTGCTGGCGTTTTTCCATAGGCTCCGCCCCCCTGACGAGCATCACAAAAATCGACGCTCAAGTCAGAGGTGGCGAAACCCGACAGGACTATAAAGATACCAGGCGTTTCCCCCTGGAAGCTCCCTCGTGCGCTCTCCTGTTCCGACCCTGCCGCTTACCGGATACCTGTCCGCCTTTCTCCCTTCGGGAAGCGTGGCGCTTTCTCATAGCTCACGCTGTAGGTATCTCAGTTCGGTGTAGGTCGTTCGCTCCAAGCTGGGCTGTGTGCACGAACCCCCCGTTCAGCCCGACCGCTGCGCCTTATCCGGTAACTATCGTCTTGAGTCCAACCCGGTAAGACACGACTTATCGCCACTGGCAGCAGCCACTGGTAACAGGATTAGCAGAGCGAGGTATGTAGGCGGTGCTACAGAGTTCTTGAAGTGGTGGCCTAACTACGGCTACACTAGAAGGACAGTATTTGGTATCTGCGCTCTGCTGAAGCCAGTTACCTTCGGAAAAAGAGTTGGTAGCTCTTGATCCGGCAAACAAACCACCGCTGGTAGCGGTGGTTTTTTTGTTTGCAAGCAGCAGATTACGCGCAGAAAAAAAGGATCTCAAGAAGATCCTTTGATCTTTTCTACGGGGTCTGACGCTCAGTGGAACGAAAACTCACGTTAAGGGATTTTGGTCATGAGATTATCAAAAAGGATCTTCACCTAGATCCTTTTAAATTAAAAATGAAGTTTTAAATCAATCTAAAGTATATATGAGTAAACTTGGTCTGACAGTTACCAATGCTTAATCAGTGAGGCACCTATCTCAGCGATCTGTCTATTTCGTTCATCCATAGTTGCCTGACTCCCCGTCGTGTAGATAACTACGATACGGGAGGGCTTACCATCTGGCCCCAGTGCTGCAATGATACCGCGAGACCCACGCTCACCGGCTCCAGATTTATCAGCAATAAACCAGCCAGCCGGAAGGGCCGAGCGCAGAAGTGGTCCTGCAACTTTATCCGCCTCCATCCAGTCTATTAATTGTTGCCGGGAAGCTAGAGTAAGTAGTTCGCCAGTTAATAGTTTGCGCAACGTTGTTGCCATTGCTACAGGCATCGTGGTGTCACGCTCGTCGTTTGGTATGGCTTCATTCAGCTCCGGTTCCCAACGATCAAGGCGAGTTACATGATCCCCCATGTTGTGCAAAAAAGCGGTTAGCTCCTTCGGTCCTCCGATCGTTGTCAGAAGTAAGTTGGCCGCAGTGTTATCACTCATGGTTATGGCAGCACTGCATAATTCTCTTACTGTCATGCCATCCGTAAGATGCTTTTCTGTGACTGGTGAGTACTCAACCAAGTCATTCTGAGAATAGTGTATGCGGCGACCGAGTTGCTCTTGCCCGGCGTCAATACGGGATAATACCGCGCCACATAGCAGAACTTTAAAAGTGCTCATCATTGGAAAACGTTCTTCGGGGCGAAAACTCTCAAGGATCTTACCGCTGTTGAGATCCAGTTCGATGTAACCCACTCGTGCACCCAACTGATCTTCAGCATCTTTTACTTTCACCAGCGTTTCTGGGTGAGCAAAAACAGGAAGGCAAAATGCCGCAAAAAAGGGAATAAGGGCGACACGGAAATGTTGAATACTCATACTCTTCCTTTTTCAATATTATTGAAGCATTTATCAGGGTTATTGTCTCATGAGCGGATACATATTTGAATGTATTTAGAAAAATAAACAAATAGGGGTTCCGCGCACATTTCCCCGAAAAGTGCCACCTGACGTCTAAGAAACCATTATTATCATGACATTAACCTATAAAAATAGGCGTATCACGAGGCCCTTTCGTCTTCAC | A Tat promoter was inserted to replace the original T5 promoter in plasmid pQE60. SDIC4B was cloned downstream of the Tat promoter. |
| pGM252 | CTCGAGAAATGTCGGTTGGCGCAAAACACGCTGATTTTTTCATCGCTCAAGGCGGGCCGTGTAACGTATAATGCGGCTTTGTTTAATCATCATCTACCACAGAGGATTTCACACAGAATTCATTAAAGAGGAGAAATTAAATGAAACACATCACAGCCCTGCTGCTGATAAGCAGCGGACTGATACTGGCATCGCTGTCGGCTGCAAATGCCCGTGCACCCGATGCCCTCGTACAAGCCCATCATCAGCGTTGCGGTAAACCGGTCTTCTATGCGCAGACCGCTAACGGCAAAAAAGAGGTGGAGATTTGCATCATTACGCCGTCGGTATCGTATTCGTTTGGTAAGACCGGTGCGGAGCATAAGGAAATGGACATTACCGTGCCGGCACACGCCACCGCTTACGCGTATCAGAACAATCAGGTTATCAGTCTTCAGGAATTCACGATCAGAAACGGCGACACCCATTACCAGGTGTCAGCCGGAACGAACGATGAAGGCAAGCCCTTTGCTTCTCTGGACGTTTACAAAGGGACGCCTGAGACGGGCAGACACCTGGCGAAAATCCAGCTCAACCCGAATACGGTCGTGAACAACATCAGCCACGCGCTTGCGGAAGAAGGCGTGGCGGGATCAGACAGTCTCTGAAAACTGGCATGTTAACTTACGGAAAATAACACAATGATAAACAAGAAATTAAAGGGAAATATTGTACCTCGTGGCGCATTTTGGCTCTGTACGCTGTTGTCTGCCCTGGCCCTGGCCCTGGCCCTGAGCGGCTGCGGTGACAAGAATGAGCGTGAATTTATTCGGGGCTGCAAGTCCGGTGGCGGTACTACCGCTATCTGTGGGTGTATCTGGGACGACCTGAAGACGAAATATACACACGGAGAGCTGGAGAAAATGAACCAGCAGTACGGTTATGCTCCTCCGCGCTTCATGGACAACATGCTGAACGCCGCCCAGCGGTGCAGAAAATAAGGACATAGAAAAATGGTACTCGTTATTAAATATTTCGCAATTGCCTTCGTCATCGCGCTTGTGGTCGTCCTGTTTAACGTTTTTGGAAACTCTGGCGAGATCCGCAGTTTCGGGCAGGGGATGGGGTATCTGTTCTGGATGACCCTGGGGCCCGGTGCCGGCATGGCCATCGGTGCCTTCTTGCGCCTGTGGCTGATGCCGGACAGCGTTTATACCACCGGCGGTGTGGGCGGTCTGCTGAAAGCCAGGCTGTTTTGGCTGATTGGCCCACAGAGCATCGGTTGGCTGGCCGGTCTGATCGCCGTGGGGAAGCAACTGATGTAACACCGTATTGTCGTCGACACGACAATCCAAAGATAACCCGCAGGGCAGTACGCCTCGCCCATTATTCAACTGATGAAGGAAAATCCAATGCGTTTATTTCATGCTGCCCTGGCGGCCATCGTGCTGGTCAGTGCTGCCACAGCACAGGCCATACCCAATATGTGGACCCGCGGTTTCGGCATGGGGGTGACTGAATACATCATCACCAGCCCCGAAAACGTGATGTTTAACCTCAACTGCACCATGAACCCGGATGAACAGAATATCTTACAACATCGCGTGCTCATCAGCCTGCCAGACGGTACTGGTGCAGATTCACGCGATGACAAAACAGCAATAACGATTGTGACGGATGACCAACAGTTCCCTTTACCCTATTCACTGGGCTGGCGAAATGGGGATAACGCCTGGATCCAGTTTATTGACGCGCTGGGTCACGCGGCAACGTTCGATGTTTATGTTAATGATAAAAAAGTAGGGCGTTTCAGTCCCGGTCTGAAAAATACGAAAAAAGAGCTGAATGACCTTGGCGGTTGCAGGAATACCGCAGGTTAAGCTTGGACTCCTGTTGATAGATCCAGTAATGACCTCAGAACTCCATCTGGATTTGTTCAGAACGCTCGGTTGCCGCCGGGCGTTTTTTATTGGTGAGAATCCAAGCTAGCTTGGCGAGATTTTCAGGAGCTAAGGAAGCTAAAATGGAGAAAAAAATCACTGGATATACCACCGTTGATATATCCCAATGGCATCGTAAAGAACATTTTGAGGCATTTCAGTCAGTTGCTCAATGTACCTATAACCAGACCGTTCAGCTGGATATTACGGCCTTTTTAAAGACCGTAAAGAAAAATAAGCACAAGTTTTATCCGGCCTTTATTCACATTCTTGCCCGCCTGATGAATGCTCATCCGGAATTTCGTATGGCAATGAAAGACGGTGAGCTGGTGATATGGGATAGTGTTCACCCTTGTTACACCGTTTTCCATGAGCAAACTGAAACGTTTTCATCGCTCTGGAGTGAATACCACGACGATTTCCGGCAGTTTCTACACATATATTCGCAAGATGTGGCGTGTTACGGTGAAAACCTGGCCTATTTCCCTAAAGGGTTTATTGAGAATATGTTTTTCGTCTCAGCCAATCCCTGGGTGAGTTTCACCAGTTTTGATTTAAACGTGGCCAATATGGACAACTTCTTCGCCCCCGTTTTCACCATGCATGGGCAAATATTATACGCAAGGCGACAAGGTGCTGATGCCGCTGGCGATTCAGGTTCATCATGCCGTCTGTGATGGCTTCCATGTCGGCAGAATGCTTAATGAATTACAACAGTACTGCGATGAGTGGCAGGGCGGGGCGTAATTTTTTTAAGGCAGTTATTGGTGCCCTTAAACGCCTGGGGTAATGACTCTCTAGCTTGAGGCATCAAATAAAACGAAAGGCTCAGTCGAAAGACTGGGCCTTTCGTTTTATCTGTTGTTTGTCGGTGAACGCTCTCCTGAGTAGGACAAATCCGCCGCTCTAGAGCTGCCTCGCGCGTTTCGGTGATGACGGTGAAAACCTCTGACACATGCAGCTCCCGGAGACGGTCACAGCTTGTCTGTAAGCGGATGCCGGGAGCAGACAAGCCCGTCAGGGCGCGTCAGCGGGTGTTGGCGGGTGTCGGGGCGCAGCCATGACCCAGTCACGTAGCGATAGCGGAGTGTATACTGGCTTAACTATGCGGCATCAGAGCAGATTGTACTGAGAGTGCACCATATGCGGTGTGAAATACCGCACAGATGCGTAAGGAGAAAATACCGCATCAGGCGCTCTTCCGCTTCCTCGCTCACTGACTCGCTGCGCTCGGTCGTTCGGCTGCGGCGAGCGGTATCAGCTCACTCAAAGGCGGTAATACGGTTATCCACAGAATCAGGGGATAACGCAGGAAAGAACATGTGAGCAAAAGGCCAGCAAAAGGCCAGGAACCGTAAAAAGGCCGCGTTGCTGGCGTTTTTCCATAGGCTCCGCCCCCCTGACGAGCATCACAAAAATCGACGCTCAAGTCAGAGGTGGCGAAACCCGACAGGACTATAAAGATACCAGGCGTTTCCCCCTGGAAGCTCCCTCGTGCGCTCTCCTGTTCCGACCCTGCCGCTTACCGGATACCTGTCCGCCTTTCTCCCTTCGGGAAGCGTGGCGCTTTCTCATAGCTCACGCTGTAGGTATCTCAGTTCGGTGTAGGTCGTTCGCTCCAAGCTGGGCTGTGTGCACGAACCCCCCGTTCAGCCCGACCGCTGCGCCTTATCCGGTAACTATCGTCTTGAGTCCAACCCGGTAAGACACGACTTATCGCCACTGGCAGCAGCCACTGGTAACAGGATTAGCAGAGCGAGGTATGTAGGCGGTGCTACAGAGTTCTTGAAGTGGTGGCCTAACTACGGCTACACTAGAAGGACAGTATTTGGTATCTGCGCTCTGCTGAAGCCAGTTACCTTCGGAAAAAGAGTTGGTAGCTCTTGATCCGGCAAACAAACCACCGCTGGTAGCGGTGGTTTTTTTGTTTGCAAGCAGCAGATTACGCGCAGAAAAAAAGGATCTCAAGAAGATCCTTTGATCTTTTCTACGGGGTCTGACGCTCAGTGGAACGAAAACTCACGTTAAGGGATTTTGGTCATGAGATTATCAAAAAGGATCTTCACCTAGATCCTTTTAAATTAAAAATGAAGTTTTAAATCAATCTAAAGTATATATGAGTAAACTTGGTCTGACAGTTACCAATGCTTAATCAGTGAGGCACCTATCTCAGCGATCTGTCTATTTCGTTCATCCATAGTTGCCTGACTCCCCGTCGTGTAGATAACTACGATACGGGAGGGCTTACCATCTGGCCCCAGTGCTGCAATGATACCGCGAGACCCACGCTCACCGGCTCCAGATTTATCAGCAATAAACCAGCCAGCCGGAAGGGCCGAGCGCAGAAGTGGTCCTGCAACTTTATCCGCCTCCATCCAGTCTATTAATTGTTGCCGGGAAGCTAGAGTAAGTAGTTCGCCAGTTAATAGTTTGCGCAACGTTGTTGCCATTGCTACAGGCATCGTGGTGTCACGCTCGTCGTTTGGTATGGCTTCATTCAGCTCCGGTTCCCAACGATCAAGGCGAGTTACATGATCCCCCATGTTGTGCAAAAAAGCGGTTAGCTCCTTCGGTCCTCCGATCGTTGTCAGAAGTAAGTTGGCCGCAGTGTTATCACTCATGGTTATGGCAGCACTGCATAATTCTCTTACTGTCATGCCATCCGTAAGATGCTTTTCTGTGACTGGTGAGTACTCAACCAAGTCATTCTGAGAATAGTGTATGCGGCGACCGAGTTGCTCTTGCCCGGCGTCAATACGGGATAATACCGCGCCACATAGCAGAACTTTAAAAGTGCTCATCATTGGAAAACGTTCTTCGGGGCGAAAACTCTCAAGGATCTTACCGCTGTTGAGATCCAGTTCGATGTAACCCACTCGTGCACCCAACTGATCTTCAGCATCTTTTACTTTCACCAGCGTTTCTGGGTGAGCAAAAACAGGAAGGCAAAATGCCGCAAAAAAGGGAATAAGGGCGACACGGAAATGTTGAATACTCATACTCTTCCTTTTTCAATATTATTGAAGCATTTATCAGGGTTATTGTCTCATGAGCGGATACATATTTGAATGTATTTAGAAAAATAAACAAATAGGGGTTCCGCGCACATTTCCCCGAAAAGTGCCACCTGACGTCTAAGAAACCATTATTATCATGACATTAACCTATAAAAATAGGCGTATCACGAGGCCCTTTCGTCTTCAC | A Tat promoter was inserted to replace the original T5 promoter in plasmid pQE60. SDIC5 was cloned downstream of the Tat promoter. |
| pGM255 | CTCGAGAAATGTCGGTTGGCGCAAAACACGCTGATTTTTTCATCGCTCAAGGCGGGCCGTGTAACGTATAATGCGGCTTTGTTTAATCATCATCTACCACAGAGGATTTCACACAGAATTCATTAAAGAGGAGAAATTAAATGGCTACCCTCGTTTTTTCCTATTCTCATGCAGATGAAGCCTTGCGTAACGAGCTCGAAACGCATCTATCACCGTTAAAACGCATGGGAACAATCAGCGCGTGGCATGACCGACGTATTGCTCCTGGGCAGGAGTTTGAGCATGAAATAGATCGCTATTTCGCCGAAGCCAATATTATTCTGTTGTTAGTCAGTAGCGACTTCATTGCATCGGATTATTGCTGGAATATTGAAATTAAGAATGCAATGGCGCGGCATGAACGAGGAGAAGCGATAGTCATCCCGGTGATTTTGCGCAATTGTGCCTGGCACAATCTTCCTTTTGGTAAGCTACTGGCCGCAACCAAAGATGGGAAACCGATAACTCAGTTTCCCAGTCATGATGACGGGTTTGTGCAAGTTGTTGATGCAGTATCAAAGGCCGCTGCCCGTATAGATACCAATAATTCAACCCAAGGGCTCCACGCTTCAGCTGCATTGGAGGCTCAACTATCACTGCCGGTAAATACAGTTTTACAACCCCGTTCTGGCAATCTCTCGCTTCCAAAAACATTCAGCGATCTTGATAGAGACCGGGCCAGCCGTGAAGGCTTTGAGTATGTTGCACGCTATTTCGAAAACTCACTCTTCGAGTTGAAACAACGGAACCCAGGTATTGAGGTGGAATTTCAACCGGTTGACGCTAGCGCATTCACCTGCGCGATTTATTTACATGGTACCCGAATGGGACAATGTGGGGTATGGCGTGGTGACAGACTCCATGGGGTCGGCTCAGTCTGCTTCAGTCATGACGGCATGGTTCGAAACAGCTACAACGAAAGTCTGAATCTAGCAGATAACGGTCAGACTCTGGGTTTTCGTACGATGATGGGGTTCACCGGGAACCGTCATAACACACTGCTGACAAATGAAGGAATGGCAGAGCATCTGTGGGATATGTTTTTCAAATCCATAAAACAACGTATGACCTGAGAGTCGGGAGGCAACAATGGTACAACTCAATCTGGTGAAAGTATTGCTGTTGAGCAATGAACGCACCGGCGATGTCCGCTATGAGATTTTCTCGAAGGAAGGTGAAGATCCTGGCTATCCGGAGAAAATCATCGTTTATCGTGAGGGAAATATCGGTGAGCATGGGGAGCGTGGCTGGATCAAGACTGACGATGTTATCAGTCTTGAACACTTGGGATTTCAACCCGGAAGGGGATTCCAGACAGCGATAACCTACCACATGCGCCCTGCACGAGATATGTGTTCAGCTATGGCTGAATGCCGACAACATTTCCAGGAAAAGTTTTGTTCTGAGAGATAAGCTTGGACTCCTGTTGATAGATCCAGTAATGACCTCAGAACTCCATCTGGATTTGTTCAGAACGCTCGGTTGCCGCCGGGCGTTTTTTATTGGTGAGAATCCAAGCTAGCTTGGCGAGATTTTCAGGAGCTAAGGAAGCTAAAATGGAGAAAAAAATCACTGGATATACCACCGTTGATATATCCCAATGGCATCGTAAAGAACATTTTGAGGCATTTCAGTCAGTTGCTCAATGTACCTATAACCAGACCGTTCAGCTGGATATTACGGCCTTTTTAAAGACCGTAAAGAAAAATAAGCACAAGTTTTATCCGGCCTTTATTCACATTCTTGCCCGCCTGATGAATGCTCATCCGGAATTTCGTATGGCAATGAAAGACGGTGAGCTGGTGATATGGGATAGTGTTCACCCTTGTTACACCGTTTTCCATGAGCAAACTGAAACGTTTTCATCGCTCTGGAGTGAATACCACGACGATTTCCGGCAGTTTCTACACATATATTCGCAAGATGTGGCGTGTTACGGTGAAAACCTGGCCTATTTCCCTAAAGGGTTTATTGAGAATATGTTTTTCGTCTCAGCCAATCCCTGGGTGAGTTTCACCAGTTTTGATTTAAACGTGGCCAATATGGACAACTTCTTCGCCCCCGTTTTCACCATGCATGGGCAAATATTATACGCAAGGCGACAAGGTGCTGATGCCGCTGGCGATTCAGGTTCATCATGCCGTCTGTGATGGCTTCCATGTCGGCAGAATGCTTAATGAATTACAACAGTACTGCGATGAGTGGCAGGGCGGGGCGTAATTTTTTTAAGGCAGTTATTGGTGCCCTTAAACGCCTGGGGTAATGACTCTCTAGCTTGAGGCATCAAATAAAACGAAAGGCTCAGTCGAAAGACTGGGCCTTTCGTTTTATCTGTTGTTTGTCGGTGAACGCTCTCCTGAGTAGGACAAATCCGCCGCTCTAGAGCTGCCTCGCGCGTTTCGGTGATGACGGTGAAAACCTCTGACACATGCAGCTCCCGGAGACGGTCACAGCTTGTCTGTAAGCGGATGCCGGGAGCAGACAAGCCCGTCAGGGCGCGTCAGCGGGTGTTGGCGGGTGTCGGGGCGCAGCCATGACCCAGTCACGTAGCGATAGCGGAGTGTATACTGGCTTAACTATGCGGCATCAGAGCAGATTGTACTGAGAGTGCACCATATGCGGTGTGAAATACCGCACAGATGCGTAAGGAGAAAATACCGCATCAGGCGCTCTTCCGCTTCCTCGCTCACTGACTCGCTGCGCTCGGTCGTTCGGCTGCGGCGAGCGGTATCAGCTCACTCAAAGGCGGTAATACGGTTATCCACAGAATCAGGGGATAACGCAGGAAAGAACATGTGAGCAAAAGGCCAGCAAAAGGCCAGGAACCGTAAAAAGGCCGCGTTGCTGGCGTTTTTCCATAGGCTCCGCCCCCCTGACGAGCATCACAAAAATCGACGCTCAAGTCAGAGGTGGCGAAACCCGACAGGACTATAAAGATACCAGGCGTTTCCCCCTGGAAGCTCCCTCGTGCGCTCTCCTGTTCCGACCCTGCCGCTTACCGGATACCTGTCCGCCTTTCTCCCTTCGGGAAGCGTGGCGCTTTCTCATAGCTCACGCTGTAGGTATCTCAGTTCGGTGTAGGTCGTTCGCTCCAAGCTGGGCTGTGTGCACGAACCCCCCGTTCAGCCCGACCGCTGCGCCTTATCCGGTAACTATCGTCTTGAGTCCAACCCGGTAAGACACGACTTATCGCCACTGGCAGCAGCCACTGGTAACAGGATTAGCAGAGCGAGGTATGTAGGCGGTGCTACAGAGTTCTTGAAGTGGTGGCCTAACTACGGCTACACTAGAAGGACAGTATTTGGTATCTGCGCTCTGCTGAAGCCAGTTACCTTCGGAAAAAGAGTTGGTAGCTCTTGATCCGGCAAACAAACCACCGCTGGTAGCGGTGGTTTTTTTGTTTGCAAGCAGCAGATTACGCGCAGAAAAAAAGGATCTCAAGAAGATCCTTTGATCTTTTCTACGGGGTCTGACGCTCAGTGGAACGAAAACTCACGTTAAGGGATTTTGGTCATGAGATTATCAAAAAGGATCTTCACCTAGATCCTTTTAAATTAAAAATGAAGTTTTAAATCAATCTAAAGTATATATGAGTAAACTTGGTCTGACAGTTACCAATGCTTAATCAGTGAGGCACCTATCTCAGCGATCTGTCTATTTCGTTCATCCATAGTTGCCTGACTCCCCGTCGTGTAGATAACTACGATACGGGAGGGCTTACCATCTGGCCCCAGTGCTGCAATGATACCGCGAGACCCACGCTCACCGGCTCCAGATTTATCAGCAATAAACCAGCCAGCCGGAAGGGCCGAGCGCAGAAGTGGTCCTGCAACTTTATCCGCCTCCATCCAGTCTATTAATTGTTGCCGGGAAGCTAGAGTAAGTAGTTCGCCAGTTAATAGTTTGCGCAACGTTGTTGCCATTGCTACAGGCATCGTGGTGTCACGCTCGTCGTTTGGTATGGCTTCATTCAGCTCCGGTTCCCAACGATCAAGGCGAGTTACATGATCCCCCATGTTGTGCAAAAAAGCGGTTAGCTCCTTCGGTCCTCCGATCGTTGTCAGAAGTAAGTTGGCCGCAGTGTTATCACTCATGGTTATGGCAGCACTGCATAATTCTCTTACTGTCATGCCATCCGTAAGATGCTTTTCTGTGACTGGTGAGTACTCAACCAAGTCATTCTGAGAATAGTGTATGCGGCGACCGAGTTGCTCTTGCCCGGCGTCAATACGGGATAATACCGCGCCACATAGCAGAACTTTAAAAGTGCTCATCATTGGAAAACGTTCTTCGGGGCGAAAACTCTCAAGGATCTTACCGCTGTTGAGATCCAGTTCGATGTAACCCACTCGTGCACCCAACTGATCTTCAGCATCTTTTACTTTCACCAGCGTTTCTGGGTGAGCAAAAACAGGAAGGCAAAATGCCGCAAAAAAGGGAATAAGGGCGACACGGAAATGTTGAATACTCATACTCTTCCTTTTTCAATATTATTGAAGCATTTATCAGGGTTATTGTCTCATGAGCGGATACATATTTGAATGTATTTAGAAAAATAAACAAATAGGGGTTCCGCGCACATTTCCCCGAAAAGTGCCACCTGACGTCTAAGAAACCATTATTATCATGACATTAACCTATAAAAATAGGCGTATCACGAGGCCCTTTCGTCTTCAC | A Tat promoter was inserted to replace the original T5 promoter in plasmid pQE60. SDIC1 was cloned downstream of the Tat promoter. |
| pGM256 | CTCGAGAAATGTCGGTTGGCGCAAAACACGCTGATTTTTTCATCGCTCAAGGCGGGCCGTGTAACGTATAATGCGGCTTTGTTTAATCATCATCTACCACAGAGGATTTCACACAGAATTCATTAAAGAGGAGAAATTAAATGAAGTGTATATCATTGCCACCGTATGCACCGACTTTGATAGAGTCAACTCGTGCAATTGGTTATACATTAGAAGCAGCAATAGCCGATATTATTGATAATAGTATCACAGCTCAGGCTTCGTGCACTGATATTTTCTTCTTCCCTACCGGGAATTCCTATATAGCCATATTGGACGATGGTTATGGAATGAATGCGGAAGAAATTGACATCGCAATGCGTTATGGCAGTCAAAATCCAAATTCTAAACGAACTGCAAATGATTTAGGCCGCTTTGGACTTGGTTTAAAAACAGCATCACTTTCGCAGTGTCGAACACTAACTGTAGTCAGCAAGCAAAGAAAGTGTATTGAAGCAAGGCGTTGGGATATTGATCATGTAATCAAAACTCAAGACTGGTCCCTTATCATCCTTGAGTCGGATGAAGAAATAAATAAAATCCCACGTATTGAGAAGTTGAAAGAAAAAGAATATGGAACTCTCATTGTATGGCAAAATCTTGACCGACTTAAAGTTGGAGAGCTTGATTTTGAACGTTCTATGGGAAAAAAAATGGATGACATGCGCAAACATCTATCATTGGTTTTTCACCGTTATATCAGTGGGGAGCCAAATCTGAAAAAACTACAGATAAGAATGAACAACACGCCTATCAGTCCTGCAGATCCATTTCTATCCCAACGAAACACGCAAGTCATGTCTGATGAGTCCATACTTTGTGAAGGCTCAAAAATTGTTATTCGTCCATATATACTACCACATATCTCTGATTTAACTAATCATGAAGTAGAATTGCTTGGGGGTAAAGAAGGTCTCAGAAAAAGCCAAGGTTTTTATGTGTATAGAAATAAAAGATTGCTGATTTGGGGAACTTGGTTTCGAATGATGCGTCAGGGAGAATGCTCTAAACTAGCTAGAGTTCAAATTGATATTCCGAATGAGTTAGATACTCTTTGGACACTAGATATAAAGAAATCAACAGCAATTCCGCCAGAAATAGTTCGTAACAATCTTGCTCCAATCATACAAAGTCTTGCAGAGAAAAGTAAAAGGACCTGGGAATTTCGTGGCAAACGCGAAATGGATGACTCGATTGAACATATTTGGCAACGATTTATAGGGAAAAGCGGTGGTTTTTATTATCAAATAAACCGCGATCACGTTCTGATAAAAGCGTTGATTAATTCATCACCGAAGTTGAAAGGAAATATGGAAAGTCTACTCAAATTGATTGAAGCAGGAATACCCTTCAATCAACTATATCTCGATTTAACATCTGAAAAACAAATTAAGAATGATATCGAAATAACTGATGTAGAAATTGATTCGATATTAAGAGGATTACTTGATCAATTTTCTACAAAAAGTGTGAAAAATGAAATGCTAGAGCAGTTAGCAATTACAGATCCTTTTATAAATTACCCACAGATAATTGCTCGCTATAAAGGAGAGTACTTAAATGGTTGCAACTAACATGGAGCAACTGGAAGGTATGATATCTTCCAGTACAAATAAAGAATACCCAGAGTATCCACCGACTGAAATTCAATTCGATGATATATCCGGAAAAATACGACAACTGCTTGAGCCTCTTTACCCTGTAACGGATGATGAATTCGACAAGATTAAGCGAAGATTAAAAGAAAAAGTTGTCGTAAAAATGGACCTAGGTGTTTTTATTACAGATAATAAACATCAACATAAATCATGGCTACCCGCTAGGCGTGCTGATTTAGATTTTTACTTCTGGAAGCGATACAAAATGTATCTGGAAGAAGTTAAAGGTTGGAACTCTCGTGTAACTGGTAGCTTAGACCGTGTATCTGATGAAATTGTGGATTTACTTGGTGATCCTAAAAGCAGTGATTCTTTTCAAAGAAGAGGTTTGGTCTTAGGAGATGTACAATCAGGAAAAACAGCAAACTATACAGCTATTTGTAATAAAGCAGCTGACACTGGATATAATGTAATAATTATCCTTGCAGGTACAATGGAGAATCTTCGTCAGCAAACACAGGAACGTCTTGATGCTGAATTCTCAGGAAGGAAGAGTGAATATTTTTTAAATCCAAAAGGAGATATTGAGAACATTCCTGTGGGGGTCGGTAAGTATGGACACGAAAAAAGAATAGAGTCATTTACTTCGGTTGTAAAAGATTTTGATAAAAATATATTACGTCAGTTAAGTCTTTCATTAAAAGGTGTAAATAACACCGTCATTTTTGTCATCAAAAAAAATAAAAGCATACTCAACAATCTTATCAGATGGTTAAAAAGTAATAATGCTGATGCAAAAGGTTTAATCTATAAGTCTTTATTATTAATTGATGATGAAGCAGATAATGCATCTGTAAATACAAATGACCCGGAAAAAGATCCAACTGCTATAAATAAAGCAATACGCGGCCTGCTACAATTATTCCGACAAGCATCTTATCTCGGCATTACGGCAACTCCTTATGCAAATATTTTTATCAATCCTGATAGTGTAGATGAGATGACTGGCGATGATTTATTTCCACGTGATTTTATTTATACACTTTCACCGCCTACTAATTATATTGGTGCTGAAGATATATTTGGAGGTGACGAACAATCACCCGCAAAATATAAGGATGCACTTAAATCAATCTATGCTAAAGAAATGGATGCTTTTTTCCCATTCAATCATAAGAAAGAACATGCAGTTACAGAGTTGCCACCCAGCATGATAGAAGCAATGGCTTACTTTTTACTGGCTACAGGCATTCGGAATATCAGGGGGGATGCAAATTCTCATCACTCAATGATGATACATACTAGCCGTTTTACCAATGTTCAAGACCAAATTAGAGATTTAGCTGGTGAATGGATAATTAGGGTTCGTTCGGATTTACAAAATTATGCTAGTTTATCGGAAAGTGAAAGTGAAAAGATTTCAAGTATTGCTTATTTAAAAGCTGTTTGGGAAAAACACGAACTTTCTAGTAAAGCAGGCAAGTTACAGAATCCAATGCCATGGCATCAATTCTTGACACAGCACCTTTATAAATCTGTTGCTCCAATCAATGTAAGAGCGGTAAACCAGAAAAGTGGGTCAACGAGCCTTGATTATTTTAATCATAAAGAAGATGGATTGCGAGTTATAGCTGTAGGCGGCAACAGTTTATCCAGGGGGCTAACACTTGAGGGGCTATGTGTTAGTTATTTTTACCGCCGTTCTAACATGTATGATACATTACTTCAAATGGGTAGATGGTTTGGTTATCGTCCTAATTATGACGATTTATTTAAGATTTGGATTAGTTCAGAAGCAATTGACTGGTATGGGTACATTACCGCAGCCGCTGAAGAACTTAAACTTGAAGTTGCGCGCATGAAAAATGCAAACCTCACTCCGATGGATTTTGGTTTGAAGGTTCGCCAAGATCCTGCGTCTCTAATCGTTACAGCAAAAAACAAAATGAGGGCTGCAACATTTGTTAAGCGCCCCATAACAGTTTCAGGGCGACTACTTGAGACCCCACGTCTAAAATCCGACACTAAAACCTTAACATATAATGAAGAAGCTTTTATAGATTTTGTTAATCGACTATCCGATGTTGGAGTGAAAGCTGAACAAAGATTTGAGAAACTCTTCTGGGAAAATGTTCACAAAGACGAAATAGAACAATTACTGCGAGATTTTAAAACACATCCTTGGCAACTAAACTTTCAAGGAGCAGCACTTGCTGACTTTATAAGAGATGATTCAAGCTTAGATTCTTGGGATGTTTATATAGCACAAGGGAGTGGAAATAAATTATATACACTCGATTGCGATAATGGGCAAATTGAAATTGCGCCTGAAATTCGTACTGTTAAGGCTTCTGGCTCTCAAATTTGCATTAGCGGAACTAAAGTGCGAGTAGGGGCAGGTGGTGCTACAAAGGTTGGTTTAAACGAAGAACAAAAGAAGGCCGCTGAGACCAAATTCAAAATGGACAATCCAAAAGCACATCATGTTCCTGATAAAGCGTATCTTTGTGTACCCCGAAACCCTATTTTAATTCTTCATGTAATAGAGGTGGATAAAGAAAAGAGCCAGATTGAAGATAATATTAAAGTCCCTGACCATCTTTTTGCTTTGGGGATAGGAATACCGGCTAATGGTAAAGAGAAAACAGCGAATTATATGGTCAATCCCATTGAGCTTCGTGGTTATAATGATTTTATTGAAGATGAAGGTAATGAAGAATGACTTTAACTCCTGAAGTACTTCGTCAGCAATGGGAAGATATAGATTACAAGGATGGTGGTTTTTTACAGATAAACATCCAACATTCACTAGAATGGTATATTGGATATCAATCTATTAGCCAGAGGACTTTACTATTATTATGTAATATGGATATCGATTCTATCGAGTCCTCTAAGTCTATACTTGTGAGTCGTCGTCGTAGAGAAGTTGATAATCGTTGGATACTAACATTTGAGTTGTTACGCAATGAACAACAAGATGTTTTTGCAATTCTTTGTTGTGACATTATAAATTACTCATCCTTTGTTGCAGATGAACAGGAGGCTCTAAGTTTAGTCATAGCACGTTATAAACAATGGGCTAAATTGCTTGAATCACAAAAACAAGGAGTAATGGATGAGCATATGCGCAAGGGATTATTAGGTGAACTACTCTTCCTAGAAAAATATATGGATAGTTGTCCCTCTATTCTATATGCAATTAATGGATGGTCAGGTCCAGAAGGTTCAGATCAAGATTTTATCTATTCTGAAGGATGGTACGAAATTAAGAGCATTGGTATTTCATCATCAAATGTCACCATTTCATCACTTGAGCAATTAGATTGCGATGAATTAGGGGAACTTGTGATAATGAGAATTGATAAAGTGCCACCAAATAAACCTAATGCAGTTTCTTTAAATGAATTAGTAAACCGAATAAAGGATAAACTATCCTTTAATCCAGAAGCGCTCGACATATTTCAACAAAAGCTAGTTTCTTATGGGTATATAGAACTGCAGGAATACTCAGAGACAAAATACCACTTTTCAAAAACTCAAAGGTATCTTGTTAGTGAATCATTCCCAAGGCTTATAAAAAAAAATGTACCAAATGAAATAGTCTCTTCAAACTATGAACTAAATCTACCATCTTTGAATAAATGGTTAAAAGTATAGAAAATGGATGCAAAGGATTTTAACAAAGATTTTATGGAAGAAGTGAAAGTCAATGCTTCAGTGACAGGAGATGGGTCATGTGCGTCTTTTGTATCTACTTTCGCTCATTACTTGCAAGAGGCTGATTTCTTGCACAATTTTACACCATCTTATTTTGAAGGGACTGGAAAATATAATCGAAAACTCCGTGTTGATGGTTATGCATATGATGAATTCGATAACACAATGAATCTAATAATTGCTGATTATGATAGTTCTGAAAGCGAACAATCCTTATCAAAAACTCAAGCAAAGCAATTGCATAATCGTTTACTAAATTTTGTACAAGACGCAAGAAACGTTGGTCTTCAACGTAGTGTAGAAATGAGTAGGCCTTGCTGGGACTTAATCGACTTGATAAAGGAAAAAAAAATTGAGGTGCGAAAATATAGGTTTATGATCTTTTCAAACTCTATCATTAACAAAAATATCGTCACTTTAGAATCATCTAAAATAGATGACGTATTGACTGAATGCCAAATATGGAATATTGACCGCTTATTCAAGATATGTGCATCTGATTCAGGAAGACATATTGTCGAAATTAATTTCAAAGATTACACTCTCAACGGAATTCCTTGTTTAGATGCAAGTGAAACAATTACAAAAGATTTTAAAAGTTACCTCTGTATAATTCCAGGTTCAGTATTAGCAGATGTGTACGATGAATATGGCAGTTTACTTTTAGAAGGTAATGTCCGATCTTTTTTATCTACTAAGGTTGCTGTTAACAAAAAGATTCGAACAACTATTATTGAATCCCCTGAAAGATTCTTTGCTTATAATAATGGGATTTCAGCAACTGCTATGAATGTGTCGATTGAGTCAACTGCTGATGGCCAGCGACTCATAGCTGCGAGTGACTTTCAGATAATAAATGGAGGGCAAACTACAGCCTCCCTTTCAAATACAAGACATAAAGATAAGTATGACTTGAGTGCGATTTTTGTACAGATGAAACTGACGGTAATTGAAAAAATCCCTGAAGAAGATGCGACTCTACTCATTCAAGACATATCACGATCCTCTAATAGCCAAAATAAAGTAAGCGATGCTGATTTCTTTTCAACCCACCCATTCCATATATGGATAGAACGTTGTTCTCAACAATTATACGCAAAAGCCATTGATGGCTCTCAATATGACACAAAATGGTTCTATGAGAGAGCGCGAGGTCAATATTTTCAAAAACAAATGCACTTAAGTCGAGCGGACAAGAAAAAATTTCTTCTACAGAATCCCAAAAGTCAGTTAATTACTAAAACTGATCTTTCAAAGGTCCGAAATTCTTGGTCTGGTTTTCCACATATTGTCAGTCGTGGTGCACAGACTAATTTCACAGACTTTGCTGACAAAATGACTAAAGTATGGAGTAATGAAAAAAATACATTACAGTCAGGAAATAAATATTTTCAAGAAACGGTCGCATTGATATTAATGTTTAAGTACATGGAAAGAATGATACCGCATCAATCATGGTACTCACAGGGATATCGTGCAAACATTATAACTTACACAATTGCTTTTCTACATAAACTAATAAAAAATAATTATAATACACGAACTCTAGACTTAATTGCAATATGGACACGACAGACTGTACCTAATATTGTCCAGGAAATATTAACTGAGTTAGCAGAAAAGGTTTATTTTAAACTTACAGATCCTTCACGTAGTGTAGAAAATGTAACACAGTGGTGTAAACGTGAAGATTGCTGGAAAAGCGTTCAGAGTATTTCCTATAAACTTCCAAATGAAATAGAAAATTATCTCACTGAATCTAATGATCAATATTTAGAGATTGAAATGGCAAAATCAAGCAACGAATGAGCTTGGACTCCTGTTGATAGATCCAGTAATGACCTCAGAACTCCATCTGGATTTGTTCAGAACGCTCGGTTGCCGCCGGGCGTTTTTTATTGGTGAGAATCCAAGCTAGCTTGGCGAGATTTTCAGGAGCTAAGGAAGCTAAAATGGAGAAAAAAATCACTGGATATACCACCGTTGATATATCCCAATGGCATCGTAAAGAACATTTTGAGGCATTTCAGTCAGTTGCTCAATGTACCTATAACCAGACCGTTCAGCTGGATATTACGGCCTTTTTAAAGACCGTAAAGAAAAATAAGCACAAGTTTTATCCGGCCTTTATTCACATTCTTGCCCGCCTGATGAATGCTCATCCGGAATTTCGTATGGCAATGAAAGACGGTGAGCTGGTGATATGGGATAGTGTTCACCCTTGTTACACCGTTTTCCATGAGCAAACTGAAACGTTTTCATCGCTCTGGAGTGAATACCACGACGATTTCCGGCAGTTTCTACACATATATTCGCAAGATGTGGCGTGTTACGGTGAAAACCTGGCCTATTTCCCTAAAGGGTTTATTGAGAATATGTTTTTCGTCTCAGCCAATCCCTGGGTGAGTTTCACCAGTTTTGATTTAAACGTGGCCAATATGGACAACTTCTTCGCCCCCGTTTTCACCATGCATGGGCAAATATTATACGCAAGGCGACAAGGTGCTGATGCCGCTGGCGATTCAGGTTCATCATGCCGTCTGTGATGGCTTCCATGTCGGCAGAATGCTTAATGAATTACAACAGTACTGCGATGAGTGGCAGGGCGGGGCGTAATTTTTTTAAGGCAGTTATTGGTGCCCTTAAACGCCTGGGGTAATGACTCTCTAGCTTGAGGCATCAAATAAAACGAAAGGCTCAGTCGAAAGACTGGGCCTTTCGTTTTATCTGTTGTTTGTCGGTGAACGCTCTCCTGAGTAGGACAAATCCGCCGCTCTAGAGCTGCCTCGCGCGTTTCGGTGATGACGGTGAAAACCTCTGACACATGCAGCTCCCGGAGACGGTCACAGCTTGTCTGTAAGCGGATGCCGGGAGCAGACAAGCCCGTCAGGGCGCGTCAGCGGGTGTTGGCGGGTGTCGGGGCGCAGCCATGACCCAGTCACGTAGCGATAGCGGAGTGTATACTGGCTTAACTATGCGGCATCAGAGCAGATTGTACTGAGAGTGCACCATATGCGGTGTGAAATACCGCACAGATGCGTAAGGAGAAAATACCGCATCAGGCGCTCTTCCGCTTCCTCGCTCACTGACTCGCTGCGCTCGGTCGTTCGGCTGCGGCGAGCGGTATCAGCTCACTCAAAGGCGGTAATACGGTTATCCACAGAATCAGGGGATAACGCAGGAAAGAACATGTGAGCAAAAGGCCAGCAAAAGGCCAGGAACCGTAAAAAGGCCGCGTTGCTGGCGTTTTTCCATAGGCTCCGCCCCCCTGACGAGCATCACAAAAATCGACGCTCAAGTCAGAGGTGGCGAAACCCGACAGGACTATAAAGATACCAGGCGTTTCCCCCTGGAAGCTCCCTCGTGCGCTCTCCTGTTCCGACCCTGCCGCTTACCGGATACCTGTCCGCCTTTCTCCCTTCGGGAAGCGTGGCGCTTTCTCATAGCTCACGCTGTAGGTATCTCAGTTCGGTGTAGGTCGTTCGCTCCAAGCTGGGCTGTGTGCACGAACCCCCCGTTCAGCCCGACCGCTGCGCCTTATCCGGTAACTATCGTCTTGAGTCCAACCCGGTAAGACACGACTTATCGCCACTGGCAGCAGCCACTGGTAACAGGATTAGCAGAGCGAGGTATGTAGGCGGTGCTACAGAGTTCTTGAAGTGGTGGCCTAACTACGGCTACACTAGAAGGACAGTATTTGGTATCTGCGCTCTGCTGAAGCCAGTTACCTTCGGAAAAAGAGTTGGTAGCTCTTGATCCGGCAAACAAACCACCGCTGGTAGCGGTGGTTTTTTTGTTTGCAAGCAGCAGATTACGCGCAGAAAAAAAGGATCTCAAGAAGATCCTTTGATCTTTTCTACGGGGTCTGACGCTCAGTGGAACGAAAACTCACGTTAAGGGATTTTGGTCATGAGATTATCAAAAAGGATCTTCACCTAGATCCTTTTAAATTAAAAATGAAGTTTTAAATCAATCTAAAGTATATATGAGTAAACTTGGTCTGACAGTTACCAATGCTTAATCAGTGAGGCACCTATCTCAGCGATCTGTCTATTTCGTTCATCCATAGTTGCCTGACTCCCCGTCGTGTAGATAACTACGATACGGGAGGGCTTACCATCTGGCCCCAGTGCTGCAATGATACCGCGAGACCCACGCTCACCGGCTCCAGATTTATCAGCAATAAACCAGCCAGCCGGAAGGGCCGAGCGCAGAAGTGGTCCTGCAACTTTATCCGCCTCCATCCAGTCTATTAATTGTTGCCGGGAAGCTAGAGTAAGTAGTTCGCCAGTTAATAGTTTGCGCAACGTTGTTGCCATTGCTACAGGCATCGTGGTGTCACGCTCGTCGTTTGGTATGGCTTCATTCAGCTCCGGTTCCCAACGATCAAGGCGAGTTACATGATCCCCCATGTTGTGCAAAAAAGCGGTTAGCTCCTTCGGTCCTCCGATCGTTGTCAGAAGTAAGTTGGCCGCAGTGTTATCACTCATGGTTATGGCAGCACTGCATAATTCTCTTACTGTCATGCCATCCGTAAGATGCTTTTCTGTGACTGGTGAGTACTCAACCAAGTCATTCTGAGAATAGTGTATGCGGCGACCGAGTTGCTCTTGCCCGGCGTCAATACGGGATAATACCGCGCCACATAGCAGAACTTTAAAAGTGCTCATCATTGGAAAACGTTCTTCGGGGCGAAAACTCTCAAGGATCTTACCGCTGTTGAGATCCAGTTCGATGTAACCCACTCGTGCACCCAACTGATCTTCAGCATCTTTTACTTTCACCAGCGTTTCTGGGTGAGCAAAAACAGGAAGGCAAAATGCCGCAAAAAAGGGAATAAGGGCGACACGGAAATGTTGAATACTCATACTCTTCCTTTTTCAATATTATTGAAGCATTTATCAGGGTTATTGTCTCATGAGCGGATACATATTTGAATGTATTTAGAAAAATAAACAAATAGGGGTTCCGCGCACATTTCCCCGAAAAGTGCCACCTGACGTCTAAGAAACCATTATTATCATGACATTAACCTATAAAAATAGGCGTATCACGAGGCCCTTTCGTCTTCAC | A Tat promoter was inserted to replace the original T5 promoter in plasmid pQE60. SDIC2 was cloned downstream of the Tat promoter. |
